# *In vitro* characterization of the colibactin-activating peptidase ClbP enables development of a fluorogenic activity probe

**DOI:** 10.1101/559385

**Authors:** Matthew R. Volpe, Matthew R. Wilson, Carolyn A. Brotherton, Ethan S. Winter, Sheila E. Johnson, Emily P. Balskus

**Affiliations:** Department of Chemistry & Chemical Biology, Harvard University, 12 Oxford Street, Cambridge, Massachusetts 02138, United States

## Abstract

The gut bacterial genotoxin colibactin has been linked to the development of colorectal cancer. In the final stages of colibactin’s biosynthesis, an inactive precursor (pre-colibactin) undergoes proteolytic cleavage by ClbP, an unusual inner-membrane-bound periplasmic peptidase, to generate the active genotoxin. This enzyme presents an opportunity to monitor and modulate colibactin biosynthesis, but its active form has not been studied *in vitro* and limited tools exist to measure its activity. Here, we describe the *in vitro* biochemical characterization of catalytically active, full-length ClbP. We elucidate its substrate preferences and use this information to develop a fluorogenic activity probe. This tool will enable the discovery of ClbP inhibitors and streamline identification of colibactin-producing bacteria.

---

The trillions of microorganisms living on and in the human body, known as the human microbiota, have been implicated in many diseases, including cancer. In particular, commensal gut bacteria harboring the *pks* island are linked to colorectal cancer (CRC).^1–3^ The *pks* island encodes a hybrid nonribosomal peptide synthetase-polyketide synthase (NRPS-PKS) assembly line that synthesizes the small molecule genotoxin colibactin.^4^ Prior work has shown that *pks*^*+*^ *E. coli* induce DNA double-strand breaks in eukaryotic cells and increase tumor load in mouse models of CRC in an inflammation-dependent fashion.^2,5,6^ While *pks*^*+*^ organisms are present in about 20% of healthy patients, they are more prevalent in the guts of those with inflammatory bowel disease (40%) and CRC (55%–67%).^2,5^ Our understanding of colibactin’s role in carcinogenesis is limited, as the active molecule has never been isolated or structurally characterized and cannot be readily detected. We sought to develop a fluorogenic probe that would allow us to measure the activity of the colibactin biosynthetic pathway in a more direct manner than current sequencing-based methods, which detect the *pks* genes. Such a tool could also accelerate the identification of small molecules that can inhibit colibactin production.

We chose the colibactin biosynthetic enzyme ClbP as a target for probe development. This periplasmic peptidase proteolytically activates colibactin in the final stages of genotoxin assembly. It removes an *N*-myristoyl-D-Asn ‘prodrug scaffold’ from an inactive precursor, precolibactin (Figure 1). Based on the structures of candidate precolibactins isolated from *clbP* deletion mutants, it has been hypothesized that cleavage by ClbP triggers the formation of an α,β-unsaturated iminium-conjugated cyclopropane which serves as an electrophilic warhead for DNA alkylation.^7,8^ Previous studies revealed that ClbP possesses three C-terminal transmembrane (TM) helices anchored in the inner membrane as well as a soluble periplasmic peptidase domain.^9^ Prior *in vitro* studies of ClbP used the truncated peptidase domain (ClbP_pep_), but this construct does not rescue genotoxicity in a *clbP* mutant and cannot process precolibactin-like synthetic substrates in whole cells.^10,11^ In fact, deletion of even a single TM helix blocks genotoxicity completely.^9^ Challenges in obtaining the full-length enzyme (ClbP_FL_) have hampered the discovery and characterization of ClbP inhibitors, with previous efforts limited to *in silico* screening with ClbP_pep_ and indirect phenotypic measurements of activity.^12^ We therefore began our work by characterizing ClbP_FL_ *in vitro*.

**Figure 1.**
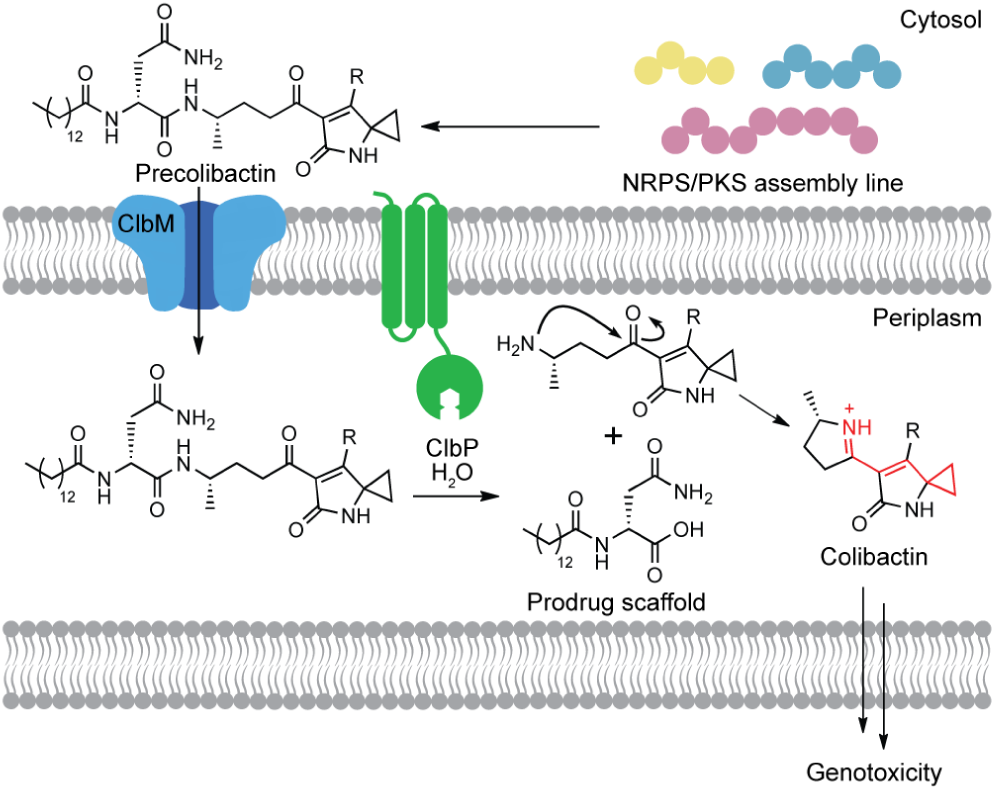
ClbP is an essential part of colibactin’s prodrug resistance mechanism. Precolibactin is synthesized by the NRPS-PKS assembly line and tailoring enzymes encoded by the *pks* island before hydrolysis by ClbP in the periplasm. Cyclization of the free amine onto an adjacent ketone enhances the electrophilicity of the cyclopropane warhead. ‘R’ = undefined, see Figure S1.

We cloned and overexpressed C-terminal His_10_-tagged ClbP_FL_, as well as an inactive mutant lacking the catalytic serine nucleophile (ClbP_FL_-S95A), in *E. coli* (Figure S2). We obtained purified enzyme by separating the cell components using ultracentrifugation, solubilizing the membranes in an n-dodecyl-β-D-maltoside (DDM)-containing buffer, and performing immobilized metal affinity chromatography. ClbP_FL_ cleaved synthetic precolibactin mimic **1** rapidly *in vitro*, while ClbP_FL_-S95A and ClbP_pep_ showed no activity by LC-MS (Figure 2A). We used *o*-phthaldialdehyde (OPA) derivatization of the amine product to determine Michaelis–Menten kinetic parameters for this transformation (Figure 2B). Compared to the related AmpC family of β-Lactamases, ClbP_FL_’s catalytic efficiency is low (Table S1), but this may be due to use of a non-native substrate.^13^

**Figure 2.**
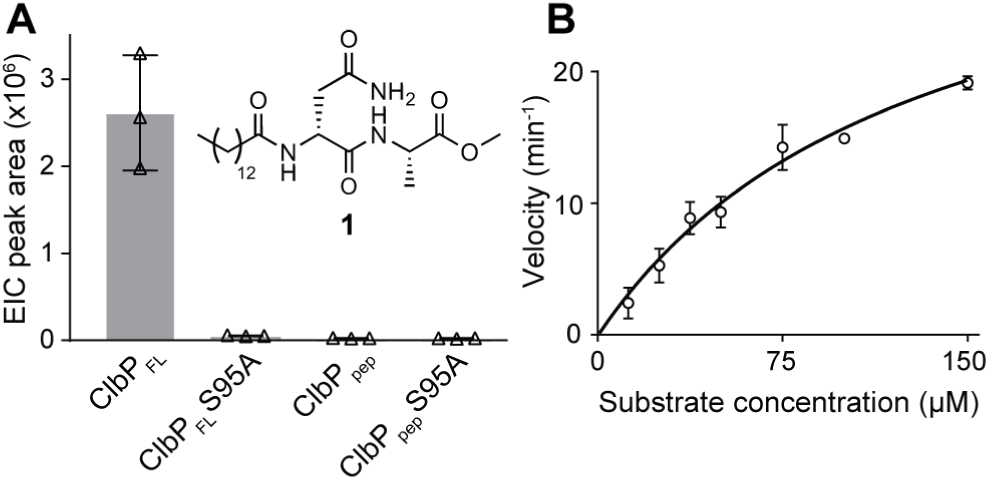
ClbP_FL_ is catalytically active *in vitro*. (a) LC-MS detection of the prodrug scaffold ([M+H]^+^: 343.2597 *m/z*) produced by ClbP-mediated hydrolysis of 100 µM **1** *in vitro*. Assays were carried out in 50 mM Tris pH 8.0, 200 mM NaCl, and 0.1 µM enzyme. 0.02% w/v DDM was added for all assays involving ClbP_FL_. Error bars represent 1 standard deviation (SD), n=3. (b) Kinetic analysis of ClbP acting on substrate **1** fit to the Michaelis–Menten model (*k*_cat_ = 36 ± 4 min^−1^, K_M_ = 130 ± 26 µM, *k*_cat_/K_M_ = 4600 ± 1100 M^−1^s^−1^). Errors are SD, n=3. Substrate is insoluble at concentrations >200 µM under these conditions.

To better understand substrate recognition by ClbP, we conducted a structure-activity relationship (SAR) study using **1** as our starting point. We first varied the *N*-acyl substituent of the prodrug scaffold. We expected that ClbP would accommodate changes at this position since *E. coli* overexpressing ClbP_FL_ can hydrolyze synthetic precolibactin mimics bearing shorter fatty acyl chains.^11^ Thus, we synthesized a series of precolibactin mimics with varied *N*-acyl chain length, polarity, and steric bulk (Table 1, entries **2**–**5**). LC-MS assays confirmed that ClbP_FL_ hydrolyzed all of these substrates at the expected peptide linkage (Table 1, Figure S3). We determined kinetic parameters for a subset of substrates. The higher *k*_cat_ and lower K_M_ values observed for more hydrophobic, long-chain substrates suggests acyl chain length is an important, though non-essential, recognition feature (Table S1, Figure S4). These observations, together with the fact that ClbP_pep_ cannot process **1** but retains some weak activity toward smaller substrates,^10^ suggest that the TM helices could play a role in substrate binding by interacting with the hydrophobic acyl substituent.

**Table 1.**
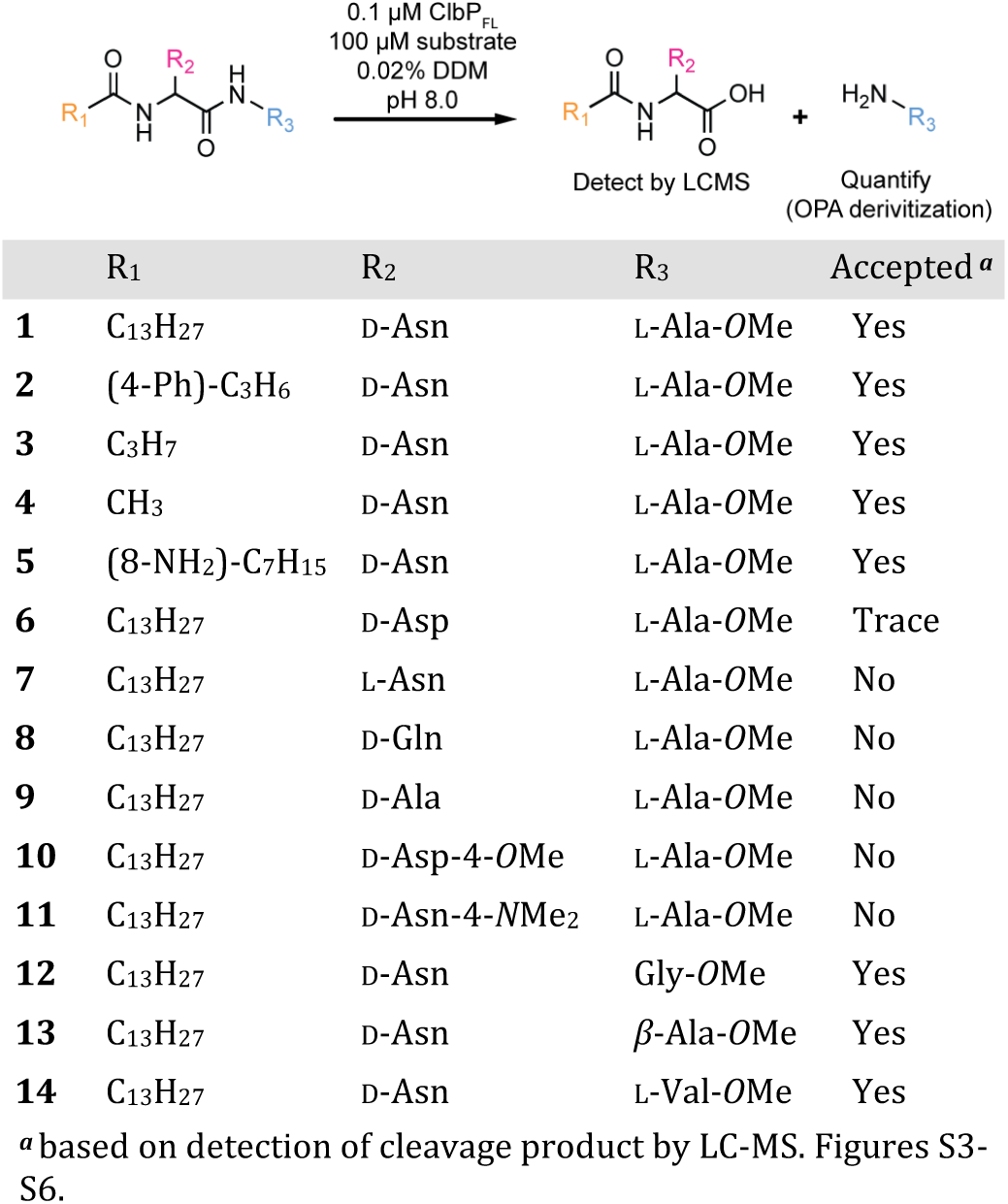
SAR study: Substrates and Approach

|  | R<sub>1</sub> | R<sub>2</sub> | R<sub>3</sub> | Accepted<sup>a</sup> |
| --- | --- | --- | --- | --- |
| 1 | C<sub>13</sub>H<sub>27</sub> | D-Asn | L-Ala-OMe | Yes |
| 2 | (4-Ph)-C<sub>3</sub>H<sub>6</sub> | D-Asn | L-Ala-OMe | Yes |
| 3 | C<sub>3</sub>H<sub>7</sub> | D-Asn | L-Ala-OMe | Yes |
| 4 | CH<sub>3</sub> | D-Asn | L-Ala-OMe | Yes |
| 5 | (8-NH<sub>2</sub>)-C<sub>7</sub>H<sub>15</sub> | D-Asn | L-Ala-OMe | Yes |
| 6 | C<sub>13</sub>H<sub>27</sub> | D-Asp | L-Ala-OMe | Trace |
| 7 | C<sub>13</sub>H<sub>27</sub> | L-Asn | L-Ala-OMe | No |
| 8 | C<sub>13</sub>H<sub>27</sub> | D-Gln | L-Ala-OMe | No |
| 9 | C<sub>13</sub>H<sub>27</sub> | D-Ala | L-Ala-OMe | No |
| 10 | C<sub>13</sub>H<sub>27</sub> | D-Asp-4-OMe | L-Ala-OMe | No |
| 11 | C<sub>13</sub>H<sub>27</sub> | D-Asn-4-NMe<sub>2</sub> | L-Ala-OMe | No |
| 12 | C<sub>13</sub>H<sub>27</sub> | D-Asn | Gly-OMe | Yes |
| 13 | C<sub>13</sub>H<sub>27</sub> | D-Asn | β-Ala-OMe | Yes |
| 14 | C<sub>13</sub>H<sub>27</sub> | D-Asn | L-Val-OMe | Yes |

<sup>a</sup>based on detection of cleavage product by LC-MS. Figures S3–S6.

We next tested the ability of ClbP_FL_ to hydrolyze substrates bearing different amino acids within the prodrug scaffold (Table 1, Entries **6**–**11**). When ClbP_FL_ was incubated with substrates containing L-Asn, D-Gln, or D-Ala (**7**–**9**), no cleavage products were detected by LC-MS (Figure S5). The D-Asp containing substrate (**6**) was processed at low levels (>10% activity relative to **1**), but we observed no hydrolysis of the corresponding D-Asp methyl ester (**10**) and D-Asn dimethylamide (**11**) substrates (Figure S5). These results indicate that this amino acid is a key recognition motif. Two closely related peptidases, ZmaM and XcnG, also hydrolyze substrates containing D-Asn at the same position *in vivo*, suggesting that this feature is evolutionarily conserved.^14,15^

We next examined ClbP’s selectivity for the second amino acid position of precolibactin mimics. Although all candidate precolibactins characterized to date have L-alanine at this location, ClbB_NRPS_, the module responsible for incorporating this building block, accepts other amino acids *in vitro*.^11^ We incubated ClbP_FL_ with substrates **12**–**14** (Table 1) and detected cleavage of all substrates by LC-MS (Figure S6). The observation that ClbP_FL_ accommodates variation at this position is in agreement with the exceptionally large groove around the catalytic residues seen in crystal structure of ClbP_pep_.^9^ The native precolibactin substrate is likely a much larger molecule than **12**–**14**, which may explain ClbP’s promiscuity toward these substrates.

Using this information, we sought to develop a fluorogenic probe for ClbP activity. Based on the key recognition elements we identified, we designed a three-component probe that would incorporate a large hydrophobic acyl substituent, the key D-Asn residue, and a caged fluorophore connected to the prodrug scaffold by a self-cleaving linker. This three-component approach has been used successfully with more canonical types of peptidases.^16–18^ Our initial target, **15**, was accessed in 9 steps from commercially available materials (Methods). Notably, the activation of **15** is analogous to precolibactin activation: hydrolysis by ClbP reveals a primary amine, which can undergo a rapid 5-*exo*-trig cyclization to produce the active species (Figure 3A). While ClbP_FL_ processed **15** *in vitro* and in live *E. coli*, its low solubility and membrane permeability likely led to poor performance in some cell-based assays (Figure S7).

**Figure 3.**
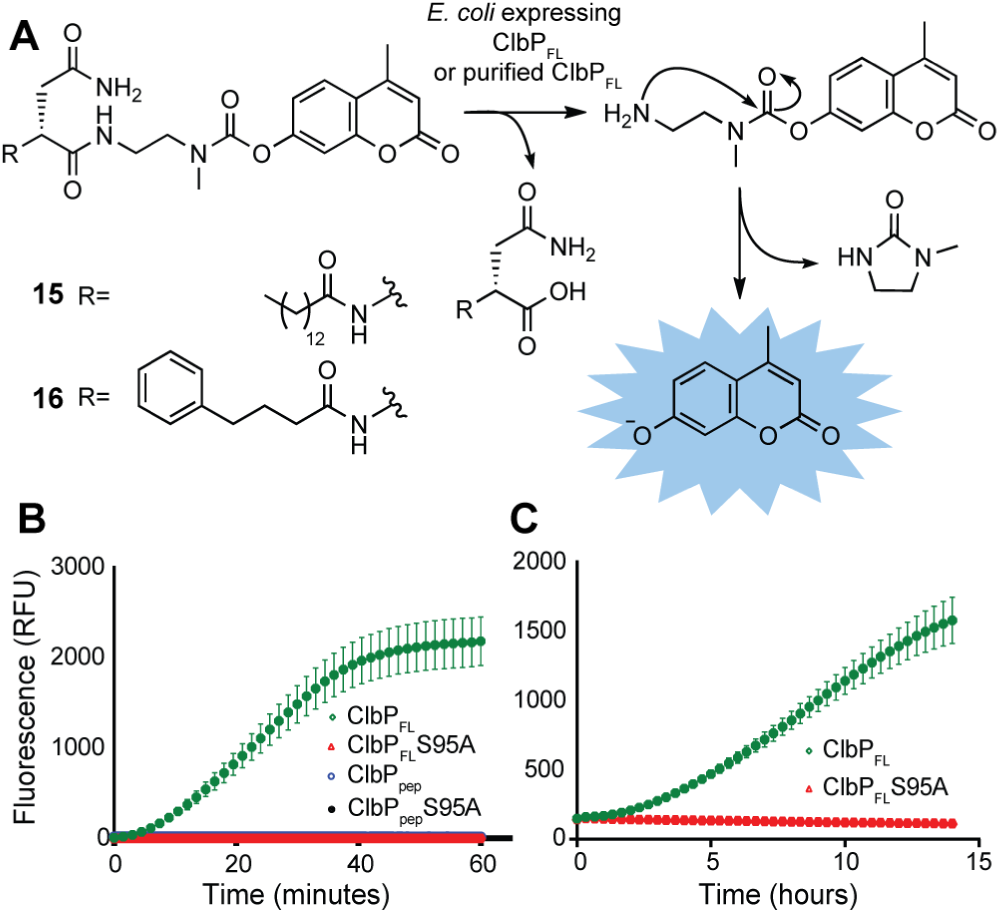
Fluorogenic probes monitor ClbP activity in whole cells and *in vitro*. (a) Mechanism of activation of fluorogenic probes. (b) **16** is activated by ClbP_FL_ but not ClbP_pep_ or ClbP_FL_-S95A *in vitro.* Points represent mean fluorescence of 6 reactions for each enzyme ± SD. (c) **16** can be activated by whole cells. Points represent mean fluorescence of 6 cultures of *E. coli* BL21 expressing each enzyme ± SD.

Because our SAR study indicated that ClbP_FL_ hydrolyzed the 4-phenylbutyryl-containing substrate **2** with similar catalytic efficiency to substrate **1** (*k*_cat_/K_M_ = 5400 ± 1800 vs 4600 ± 1100 M^−1^s^−1^, Table S1), we synthesized an analogous probe containing the same modification (**16**, Figure 3) in hopes of improving probe solubility and performance. **16** was cleaved rapidly by ClbP_FL_ *in vitro* and showed negligible background activity with the S95A mutant and ClbP_pep_ (Figure 3B). It also exhibited excellent stability in DMSO stock solutions, *in vitro* assay conditions, and LB and MEGA growth media. In an *in vitro* assay containing 50 µM **16**, cleavage by ClbP_FL_ results in a >100-fold increase in fluorescence relative to the control in less than 30 minutes (Figure 3B). Probe **16** also shows robust and consistent cleavage in a whole-cell assay format with *E. coli* over-expressing ClbP (Figure 3C).

We next assessed whether our probe could detect ClbP activity in wild-type *pks*^*+*^ strains. For this assay to provide a reliable read out, it must be sensitive enough to be activated by native levels of ClbP. Indeed, we reliably detected activation of probe **16** after overnight incubation with the *pks*^*+*^ strains *E. coli* CCR20 and *E. coli* Nissle 1917, as well as with a strain heterologously expressing the *pks* island (BW25113 BAC*pks*) (Figure 4A).

**Figure 4.**
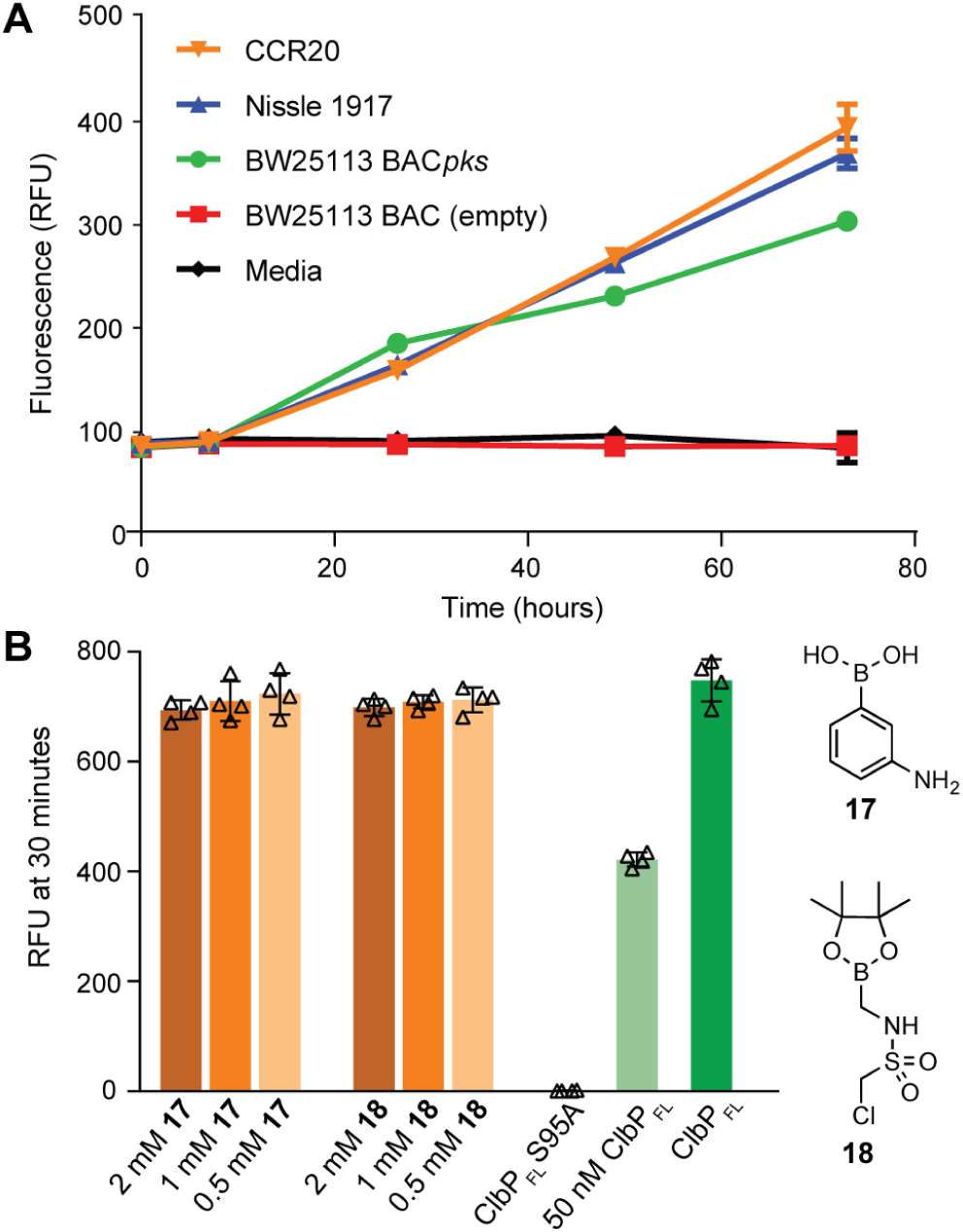
Probe **16** can identify *pks*^*+*^ organisms and assess inhibition by small molecules. (a) Wild-type *pks*^*+*^ *E. coli* strains were grown in MEGA medium under anaerobic conditions with 100 µM **16** (Error bars are SD, n=3, RFU values measure whole culture). (b) Purified ClbP_FL_ and the indicated concentration of **17** or **18** were incubated for 1 hour and fluorescence of each reaction was measured 30 minutes after addition of **16** (Error bars are SD, n=4, 0.1 µM enzyme except where indicated.) Differences between treatment groups and control were not significant (Student’s two-tailed *t* test, *P* > 0.05).

Finally, we used probe **16** to test the inhibition of ClbP by small molecules. Boronic acids **17** and **18** were identified previously by *in silico* docking with the ClbP_pep_ crystal structure.^12^ These molecules block the genotoxicity of *pks*^*+*^ *E. coli* toward mammalian cells in tissue culture, but their effect on the catalytic activity of ClbP has not been reported. When ClbP_FL_ was incubated *in vitro* with either **17** or **18** and probe **16**, we observed a small decrease in the fluorescent signal produced, but this change was not statistically significant. This assay was also run under the same conditions with half the amount of ClbP_FL_ present. This control showed approximately 50% activity, confirming that the assay is sensitive to modest changes in the activity of the enzyme (Figure 4B). We also used a more sensitive LC-MS assay to evaluate the inhibition of ClbP_FL_ with substrate **1.** In this format, we observed weak but statistically significant inhibition of ClbP_FL_ (10–15% inhibition at 2 mM **17**, Figure S8). The weakness of this effect suggests that these molecules abrogate genotoxicity through other cellular targets, and there remains a need for potent ClbP inhibitors.

Despite over a decade of research, the colibactin genotoxin remains elusive, and evidence of its carcinogenicity relies on correlations and models that cannot fully recapitulate the complexity of the gut microbiota. To better understand this intricate pathway, we have characterized the essential colibactin-activating peptidase ClbP *in vitro* and developed a fluorogenic probe for its activity. Our SAR study provides a starting point for the rational design of ClbP inhibitors, while our fluorogenic assay will allow the use of high-throughput screening toward the same goal. Such inhibitors would enable more detailed studies of colibactin’s effects in physiologically-relevant animal models and in complex microbiotas. The fact that our probe can detect ClbP activity in wild-type *pks*^*+*^ strains also opens up the possibility of activity-based diagnostics to detect these organisms.

ClbP is the most well-characterized enzyme in a larger family of prodrug-activating peptidases. Though homologs ZmaM and XcnG have been studied, ClbP is the only member of this family to be purified and characterized *in vitro*. Qian and coworkers have shown that additional peptidases that likely recognize *N*-acyl-D-Asn motifs are encoded in cryptic gene clusters.^19^ Our fluorogenic probe may be adaptable to these uncharacterized homologs, enabling the direct monitoring of other secondary metabolite pathways. Overall, this work shows how detailed characterization of biosynthetic enzymes can enable the development of innovative tools to interrogate natural products.

## AUTHOR INFORMATION

### Notes

The authors declare the following competing financial interest: a provisional patent has been filed on the fluorogenic probes described here which lists M.R.V., M.R.W. and E.P.B. as co-inventors (U.S. Provisional Application No.: 62/719,325).

## ACKNOWLEDGMENT

The authors thank F. Prati for providing compound **18**, R. Müller and R. Bonnet for providing bacterial strains, Y. Jiang and P. Boudreau for input on experimental design, J. Velilla Garcia and R. Gaudet for plasmids and helpful discussion, C. Chittim and M. Bollenbach for comments on the manuscript, and D. Kahne and coworkers for use of equipment. The authors acknowledge financial support from the National Institutes of Health (R01CA208834) and the Damon Runyon-Rachleff Innovation Award. M.R.W acknowledges support from the American Cancer Society-New England Division Postdoctoral Fellowship (PF-16-122-01-CDD). S.E.J acknowledges support from The Amgen Scholars Program.

