## Supplemental Information for "*In vitro* characterization of the colibactin-activating peptidase ClbP enables development of a fluorogenic activity probe"

**Supplementary Information**

### 1. Supplemental discussion

The genotoxin colibactin and its inactive precursor, precolibactin, have not yet been fully characterized. There have, however, been numerous reports of candidate precolibactins which have been isolated from *pks*$^+$ strains of *E. coli* which do not express ClbP (Figure S1).[1–7] Some candidate precolibactins are believed to be derived from on-pathway biosynthetic intermediates, while others are believed to be shunt metabolites. The mechanism of colibactin activation presented in Figure 1 has been supported by the structures of several of these candidate precolibactins as well as by the preparation of synthetic model precolibactins,[4] biochemical studies of ClbP activity in whole cells,[8] and the identification of putative colibactin-derived DNA adducts.[9] Based on the structures of candidate precolibactins identified to date, as well as the biosynthetic enzymes annotated in the *pks* island, the 'R' group as presented in Figure 1 likely incorporates a glycine extender unit, one or more thiazole or thiazoline rings, and at least one structural motif derived from the unusual PKS building block aminomalonate.

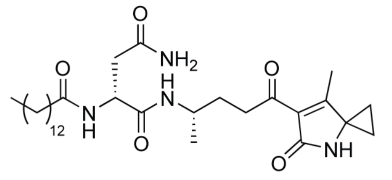

**Precolibactin '547'**
Vizcaino *et al* 2015, Brotherton *et al* 2015, Bian *et al* 2015

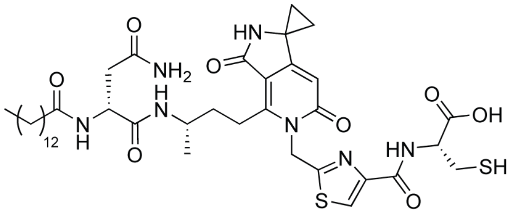

**Precolibactin '816'**
Vizcaino *et al* 2015, Healy *et al* 2016

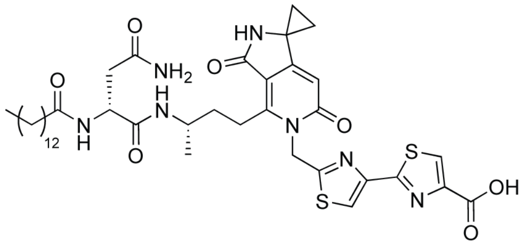

**Precolibactin '796'**
Li *et al* 2015, Zha *et al* 2016

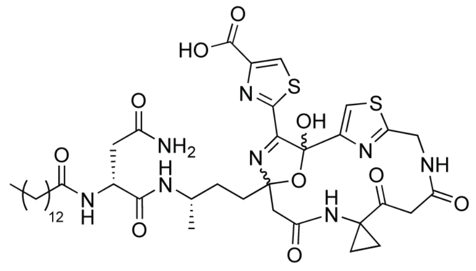

**Precolibactin '887'**
Li *et al* 2016

**Figure S1 – Structures of selected candidate precolibactins.** These compounds are named according to the mass by which they were first identified *via* mass spectrometry for clarity. These metabolites, all of which have been isolated from strains of *pks*$^+$ bacteria which do not possess a functional copy of ClbP, possess specific structural features that may be present in the final genotoxin. Metabolites containing a pyridone group and larger macrocycles (Precolibactins '816', '796', and '887' above) cannot cyclize upon cleavage by ClbP as shown in Figure 1, which suggests that they are shunt products from the biosynthetic pathway that accumulate in the absence of ClbP.

### 2. General materials and methods

Oligonucleotide primers were synthesized by Integrated DNA Technologies (Coralville, IA). Recombinant plasmid DNA was purified with a Qiaprep Kit from Qiagen (Hilden, Germany). Gel extraction of DNA fragments and restriction endonuclease clean up were performed using an Illustra GFX PCR DNA and Gel Band Purification Kit from GE Healthcare. DNA sequencing was performed by Eton Bioscience, Inc. (Boston, MA). Nickel-nitrilotriacetic acid agarose (Ni-NTA) resin was purchased from Qiagen. SDS-PAGE gels were purchased from BioRad (Hercules, CA). Purified protein concentrations were determined by absorbance measurements at 280 nanometers taken on a NanoDrop 2000 UV-Vis Spectrophotometer (ThermoFisher Scientific, Waltham, MA) using extinction coefficients calculated based on protein sequences using the ExPASY online tool (https://web.expasy.org/protparam/). Luria-Bertani (LB) and Terrific Broth (TB) media were obtained from EMD Millipore (Darmstadt, Germany) or Alfa Aesar (Tewksbury, MA). Isopropyl β-*D*-1-thiogalactopyranoside (IPTG) was purchased from Teknova (Hollister, CA) and Dodecyl maltoside (DDM) was purchased from Chem Impex International Inc (Wood Dale, IL). All other chemicals were purchased from Sigma-Aldrich (St. Louis, MO) unless otherwise noted.

Optical densities of *E. coli* cultures were determined with a DU 730 Life Sciences UV/Vis spectrophotometer (Beckman Coulter, Indianapolis, IN) by measuring absorbance at 600 nm. Ultracentrifugation was performed on a Beckman-Coulter Optima XE-90 Ultracentrifuge fitted with a type 45 Ti rotor in 70 mL polycarbonate tubes. Solvents and formic acid used for LC-MS were B & J Brand High Purity Solvents (Honeywell Research Chemicals). High-resolution LC-MS analyses of enzyme assays and synthetic compounds were performed on an Agilent 6530 Q-TOF Mass Spectrometer fitted with a dual-spray electrospray ionization (ESI) source. The capillary voltage was set to 3.5 kV, the fragmentor voltage to 175 V, the skimmer voltage to 65 V, and the Oct 1 RF to 750 V. The drying gas temperature was maintained at 275 °C with a flow rate of 8 L/min and a nebulizer pressure of 35 psi. A standard calibrant mix was introduced continuously during all experiments via the dual-spray ESI source. For analytical liquid chromatography, unless otherwise noted, experiments were performed using an Agilent Technologies 1200 series LC on a Hypersil GOLDaq C18 reverse phase column (50 x 3 mm, Thermo Scientific) with the following elution conditions: a gradient from 35% solvent A:65% solvent B to 100% solvent A over 5 minutes, holding at 100% A for 2 minutes, followed by a gradient back to 35 % solvent A over 1 minute and holding at 35% solvent A for 3.5 minutes (solvent A: acetonitrile + 0.1% formic acid; solvent B: water + 0.1% formic acid; flow rate = 0.4 mL/minute; injection volume = 10 μL). All experiments were performed in positive ion mode, and the masses detected corresponded to $[M+H]^+$ ions unless otherwise noted. Masses were found within 5 ppm of the expected $[M+H]^+$ masses.

NMR chemical shifts are reported in parts per million downfield from tetramethylsilane using the solvent resonance as internal standard for $^1H$ ($CDCl_3$ = 7.26 ppm, DMSO-$d_6$ = 2.50 ppm) and $^{13}C$ ($CDCl_3$ = 77.25 ppm, DMSO-$d_6$ = 39.52 ppm). Data are reported as follows: chemical shift, integration multiplicity (s = singlet, d = doublet, t = triplet, q = quartet, quint = quintet, sext = sextet m = multiplet), coupling constant, integration, and assignment. All solvents for synthesis were obtained from Sigma-Aldrich, with the exception of methanol (EMD Millipore). All NMR solvents were purchased from Cambridge Isotope Laboratories (Tewksbury, MA). NMR spectra were collected in the Magnetic Resonance Laboratory in Harvard University Department of Chemistry and Chemical Biology and visualized and processed using MestreNova, version 11.0.2-18153 (Mestrelab Research S.L., Escondido, CA) Optical rotation data were obtained using a 1 mL cell with a 0.5 dm path length on a Jasco P-2000 polarimeter equipped with a sodium (589 nm, D) lamp.

Preparative HPLC purification was run on a Dionex Ultimate 3000 instrument (Thermo Scientific) using Hypersil GOLDaQ column (250 mm x 20 mm, 5 µm particle size, Thermo Scientific). Standard LC gradient conditions: 50% Solvent A for 2.5 minutes, gradient to 95% Solvent A over 7.5 minutes, hold at 95% Solvent A for 11.5 minutes, gradient to 50% solvent A over 1 minute, hold at 50% solvent A for 2.5 minutes (solvent A: HPLC-grade acetonitrile (VWR, HiPerSolv-Chromanorm) + 0.1% formic acid; solvent B: water + 0.1% formic acid; flow rate: 8 mL/minute; injection volume: 200 to 400 µL). All solvents were degassed by sonication prior to use. Compounds which were purified using different conditions are noted in the synthetic methods section below for that compound.

Strains of *E. coli* DH10B harboring pBeloBAC11-*pks*, as well as *E. coli* CCR20 were obtained from the Bonnet lab, Laboratoire de Bactériologie Clinique, Centre Hospitalier de Clermont-Ferrand, Clermont-Ferrand F-63003, France. The strain *E. coli* Nissle 1917 was obtained from the Müller lab, Helmholtz Institute for Pharmaceutical Research Saarland, Campus E8.1, 66123 Saarbrücken, Germany.

### 3. Cloning, overexpression, and purification of $ClbP_{FL}$, $ClbP_{FL}$S95A, $ClbP_{pep}$, and $ClbP_{pep}$S95A

**Mutagenesis to extend histidine tag:** Plasmids for the expression of $ClbP_{FL}$ and $ClbP_{FL}$-S95A have been previously reported.[10] These plasmids (pET29b-$ClbP_{FL}$-$CHis_6$ and pET29b-$ClbP_{FL}$-S95A-$CHis_6$ were used as templates for site-directed mutagenesis with the following primers: Forward - CCACCATCACCATCACTGAGATCCGGCTGCTAACAAAGCCCGAAAG, Rev - CTCAGTGATGGTGATGGTGGTGGTGGTGGTGGTGCTCGAGCTC. Reactions (50 µL total volume) were performed with 35.5 µL $H_2O$, 5 µL 10X Pfu Ultra II buffer, 2.5 µL of NEB dNTP mix (10 mM each dATP, dCTP, dGTP, dTTP), 1 µL Pfu Ultra II, 1 µL template at 75 ng/µL, 2.5 µL 10 µM forward primer, 2.5 µL 10 µM reverse primer. For PCR, reaction mixtures were heated to 95 °C for 30 seconds and then were cycled 20 times at 95 °C for 30 seconds, 50 °C for 1 minute, 68 °C for 7 minutes, followed by a final extension at 68 °C for 8 minutes. Successful amplification was confirmed by running agarose gel electrophoresis, and template DNA was removed by digestion for 1 hour at 37 °C with 1 µL of DpnI. Digested reactions were used to transform chemically competent *E. coli* DH5α cells, and plasmids were isolated from cultures of single colonies. Mutagenesis was confirmed by sequencing isolated plasmid DNA.

**Transformation of *E. coli* for expression of $ClbP_{FL}$-$CHis_{10}$:** OverExpress C41(DE3) chemically competent cells were obtained from Lucigen (Madison, WI) and transformed with the constructs described above following the manufacturer's protocol. Following 1 hour of recovery in SOC medium after heat shock, each transformation was plated on to fresh LB agar plates containing 50 µg/ml kanamycin and incubated at 37 °C overnight.

**Transformation of BW25113 cells with BAC for *pks* activity assays:** The pBeloBAC11-*pks* BAC was isolated from *E. coli* DH10B harboring pBeloBAC11-*pks* using a BAC DNA Miniprep Kit (Zymo Research, Irvine, CA). The empty pBeloBAC11 BAC was isolated from *E. coli* K12 ER2420 harboring pBeloBAC11 (New England Biolabs, Ipswich, MA) using a Plasmid Miniprep Kit (Qiagen). To generate BW25113 *E. coli* strains harboring colibactin biosynthesis genes, electrocompetent BW25113 *E. coli* were electroporated (BioRad MicroPulser, 1.8 kV, 0.1 cm cuvette) with pBeloBAC11-*pks*, pBeloBAC11-*pks*Δ*clbP*, or pBeloBAC11. The transformed *E. coli* was then selected on LB agar plates containing either 25 µg/mL chloramphenicol or 25 µg/mL chloramphenicol and 50 µg/mL kanamycin.

**Large scale overexpression and purification of $ClbP_{FL}$ and $ClbP_{FL}$-S95A:**

A 50 mL starter culture of either *E. coli* C41 pET-29b-$ClbP_{FL}$-$CHis_{10}$ or *E. coli* C41 pET-29b-$ClbP_{FL}$-S95A-$CHis_{10}$ was inoculated from a single colony from an LB + kanamycin agar plate and grown overnight at 37 °C in LB medium (Lennox, Alfa Aesar) supplemented with 50 µg/mL kanamycin. Overnight cultures were diluted 1:100 into 2 L of TB medium containing 50 µg/mL kanamycin. Cultures were incubated at 37 °C with shaking at 175 rpm, moved to 15 °C at $OD_{600}$ = 0.5-0.8 and then induced with 500 µM IPTG. Cultures were incubated with shaking at 15 °C for approximately 20 h. Cells from the 2 L culture were harvested by centrifugation (6,000 x *g* for 15 min at 4 °C). Cell pellets were flash frozen in liquid $N_2$ and stored at –80 °C for a maximum of 2 days. Pellets were thawed and resuspended in lysis buffer (20 mM Sodium phosphate, pH 8.0, 500 mM NaCl, 55 mM imidazole, 5 mM $MgCl_2$, 0.25 mg/mL Lysozyme, 10 µg/mL DNase I). Cells were lysed in 50 mL conical tubes on ice using a Branson Digital sonifier (Fisher Scientific, Hampton, NH) [10 second pulse at 25% amplitude with 20 seconds of rest for a total pulse time of 5 minutes, pausing to mix tube by inversion after 2.5 minutes]. Unbroken cells were removed by centrifugation (2,000 x *g* for 10 minutes at 4 °C). Supernatants were transferred to ultracentrifugation tubes, and membranes were isolated by centrifugation at 35,000 RPM for 70 minutes at 4 °C (average RCF for a Beckman-Coulter Type 45 Ti rotor at 35,000 RPM is approximately 95,000 x *g*). The supernatant was removed, and the membrane pellet was resuspended with a detergent buffer (20 mM Sodium phosphate, pH 8.0, 500 mM NaCl, 55 mM imidazole, 1.5% w/v DDM + Pierce™ Protease Inhibitor (Thermo Scientific)) using an IKA T18 Basic Ultra-Turrax homogenizer (IKA Works, Inc., Wilmington, NC). The homogenate was nutated for 3 hours at 4 °C to dissolve membranes. The homogenate was centrifuged to remove insoluble cell debris (95,000 x *g* for 35 minutes at 4 °C) and the supernatant containing solubilized cell membranes was collected. Ni-NTA resin (~1 mL per L of expression culture) was added to a gravity-flow column and washed with 3 column volumes of wash buffer (50 mM Tris, pH 8.0, 200 mM NaCl, 55 mM imidazole, 0.02% w/v DDM). Resin was resuspended in a minimal volume of wash buffer and added to the clarified membrane homogenate. The mixture was then incubated on a nutating mixer at 4 °C for 2 hours before collecting the resin by centrifugation (4,000 x *g* 10 minutes at 4 °C). The resin was resuspended in a minimal volume of the residual homogenate and transferred by pipette to a gravity flow column fitted with a plastic stopcock. The resin was then washed with 2 column volumes of wash buffer and the protein eluted using a stepwise gradient of increasing imidazole in the wash buffer (55 mM, 100 mM, 200 mM, 450 mM). Fractions were collected for each column volume of wash buffer and analyzed by SDS-PAGE (4–15% Tris-HCl gel) for the presence of the desired protein. Fractions containing the desired protein were pooled and dialyzed twice against 200 volumes of storage buffer for each round (50 mM Tris, pH 8.0, 200 mM NaCl, 0.02% w/v DDM), first for 4 hours, then overnight. The dialyzed protein was then concentration in 100 kDa MWCO* Amicon® Ultra spin filters (Corning, Corning, NY). Concentrated protein was then flash frozen as 10-15 µL droplets in liquid $N_2$ and stored at –80 °C. Yield: 1.4 - 1.8 mg/L culture

*Although the mass of $ClbP_{FL}$ is only ~50 kDa, a larger molecular weight cut-off (MWCO) is necessary for this step because membranes with lower MWCOs do not permit DDM to pass

through efficiently, resulting in a significant over concentration of the detergent. Analysis of the protein concentrate and flow through by $A_{280}$ measurements showed that $ClbP_{FL}$ is completely retained by a 100 kDa MWCO filter, which is likely due to the larger effective size of the complex formed by the enzyme and DDM micelle.

**Large scale overexpression and purification of $ClbP_{pep}$ and $ClbP_{pep}$-S95A:**
$ClbP_{pep}$ and $ClbP_{pep}$-S95A were purified as previously reported using the same plasmids and expression strains, with the modifications described below.[10]

A 50 mL starter culture of *E. coli* BL21 pET-29b-$ClbP_{pep}$ or pET-29b-$ClbP_{pep}$-S95A was inoculated from frozen stock and grown overnight at 37 °C in LB medium supplemented with 50 µg/ml kanamycin. Overnight cultures were diluted 1:100 into 2 L of LB medium containing 50 µg/mL kanamycin. Cultures were incubated at 37 °C with shaking at 175 rpm. Protein expression was induced with 500 µM IPTG at $OD_{600}$ = 0.6-0.7, and cultures were moved to 15 °C and incubated for ~ 20 h. Cells from the 2 L culture were harvested by centrifugation (6,000 x *g* for 15 min) and resuspended in Tris/Sucrose buffer (30 mM Tris-HCl, pH 8.0, 20 wt% sucrose), 80 mL buffer per gram of cell pellet. To this mixture was added EDTA (1 mM final concentration) dropwise with stirring. This mixture was allowed to stir at 4 °C for 10 min and was then centrifuged (6,000 x *g* for 20 min at 4 °C). The supernatant was removed, and the cell pellet was resuspended in 400 mL of 5 mM $MgSO_4$. The suspension was allowed to stir at 4 °C for 10 min, and was then centrifuged (11,000 x *g* for 20 min at 4 °C). The supernatant was incubated with 3 mL of Ni-NTA resin and 5 mM imidazole for 2 h at 4 °C. This mixture was then poured onto a glass column and the solution eluted to give ~ 2 mL of Ni-NTA resin in a 5 mM imidazole solution. Protein was eluted from the column using a stepwise imidazole gradient (25 mM, 50 mM, 75 mM, 100 mM, 125 mM, 150 mM, 200 mM) in elution buffer (50 mM Tris-HCl, pH 8.0, 200 mM NaCl), collecting 2 mL fractions. SDS–PAGE analysis (4–15% Tris-HCl gel) was employed to ascertain the presence and purity of protein in each fraction. Fractions containing the desired protein were combined and dialyzed twice against 2 L of storage buffer (50 mM Tris-HCl, pH 8.0, 200 mM NaCl, 10% (v/v) glycerol). This procedure afforded yields of 0.27 mg/L for $CHis_6$-tagged $ClbP_{pep}$, and 0.21 mg/L for $CHis_6$-tagged $ClbP_{pep}$-S95A.

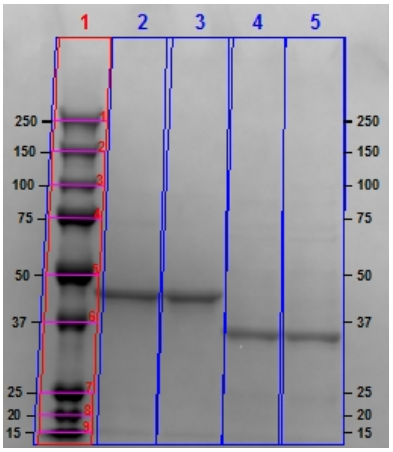

**Figure S2** – $ClbP_{FL}$ and $ClbP_{pep}$ analyzed by SDS-PAGE (4-12% Bis-Tris NuPAGE gel, ThermoFisher). Lanes: 1- Precision Plus Protein™ All Blue standard (BioRad), 2- $ClbP_{FL}$-$CHis_{10}$, 2- $ClbP_{FL}$-S95A-$CHis_{10}$, 3- $ClbP_{pep}$-$CHis_6$, 4- $ClbP_{pep}$-S95A-$CHis_6$. Numbers in black indicate masses of standards in kDa.

### 4. Biochemical characterization of $ClbP_{FL}$

#### LC-MS assays for substrate cleavage

For all *in vitro* assays involving $ClbP_{FL}$ and $Clb_{FL}$-S95A "Tris assay buffer" refers to a buffer composed of 50 mM Tris, 200 mM NaCl, 0.02% w/v DDM, pH 8.0. Likewise, "phosphate assay buffer" refers to 50 mM sodium phosphate, 200 mM NaCl, 0.02% w/v DDM, pH 8.0.

**SAR Study:** Assays to determine which substrates were accepted and hydrolyzed were conducted in standard Tris assay buffer. Substrates were prepared as 10 mM stock solutions in DMSO, then diluted to 120 µM in standard Tris assay buffer. $ClbP_{FL}$-$CHis_{10}$ and $ClbP_{FL}$-S95A-$CHis_{10}$ were defrosted from –80 °C stocks on ice, then diluted to 0.6 µM in Tris assay buffer. 25 µL of enzyme was deposited in individual wells of a 96-well plate in triplicate. Reactions were initiated by the addition of 125 µL of the appropriate substrate in Tris assay buffer to a final concentration of 0.1 µM enzyme, 100 µM substrate, 1% DMSO and incubated at 25 °C. A 20 µL sample of each reaction mixture was immediately diluted into 180 µL of cold LC-MS-grade methanol as the "t0" time point. Reactions were then allowed to incubate at room temperature for 5 hours, at which point another 20 µL of each reaction mixture was diluted into 180 µL of cold LC-MS methanol and mixed by pipetting ("t5"). 96-well plates containing the methanol-quenched reactions were sealed with adhesive foil and stored at –20 °C overnight. Plates were then centrifuged at 3,880 x *g* at 4 °C for 15 minutes and 20 µL of the supernatant was diluted again into 180 µL of LC-MS-grade methanol in an Agilent 96-well LC-MS autosampler plate. Samples were analyzed by LC-MS following the procedures outlined in the general materials and methods, with the exception of substrates **3**, **4**, and **5** for which the following chromatography method was used: 0% solvent A in solvent B for 30 seconds, a gradient from 0% A to 50% A over 9.5 minutes, followed by a gradient to 90% A over 2 minutes, holding at 90% A for 3 minutes, a gradient back to 0% A over 1 minute, hold at 0% A for 3 minutes (solvent A = acetonitrile + 0.1% formic acid; solvent B = water +0.1% formic acid. Flow rate = 0.4 mL/minute, injection volume = 10 µL). Relative activity was determined by measuring the difference in Extracted Ion Chromatogram (EIC) peak area for the expected mass of the prodrug scaffold fragment (± 5 ppm) between the "t5" and "t0" samples and comparing this value for each substrate to substrate **1** which was used as a control in each sample group. For comparing substrates **1-5**, a standard curve for each substrate was used to quantify the molar consumption of substrate, since the significant chemical variation in $R_1$ position and different LC conditions make direct comparison of the EIC signal more prone to error. All substrates were tested in triplicate.

***In vitro* kinetics assays with $ClbP_{FL}$:** Assays were run in triplicate in 1.5 mL deep-well plates. $ClbP_{FL}$-$CHis_{10}$ was defrosted from –80 °C stocks on ice, then buffer exchanged into phosphate assay buffer using Zeba Spin 40K MWCO desalting columns (ThermoFisher Scientific). The enzyme was then diluted to 0.6 µM in phosphate assay buffer and 50 µL of this stock was deposited in the appropriate wells. The substrate of interest was prepared in a 2-fold dilution series in phosphate assay buffer. Reactions were initiated by the addition of 250 µL of substrate stock to each assay well to a final concentration of 0.1 µM ClbP, 12.5 – 200 µM substrate, 3% DMSO and incubated at 25 °C. Reaction mixtures were pipetted briefly to mix. At each time point, 40 µL of each reaction was quenched by adding to 80 µL of cold methanol in the wells of a standard 96-well plate. Standard curves were prepared in triplicate by dissolving 1 – 100 µM L-alanine methyl ester in phosphate assay buffer with 0.1 µM ClbP and 3% DMSO and following the same quenching and sample preparation steps. Plates containing the quenched reaction samples were sealed and spun down at 3,880 x *g* at 4 °C for 15 minutes. An opaque black 384-well flat-bottom plate was prepared by combining 3 mL of the derivatization reagent ("Phthaldialdehyde Reagent Solution, Incomplete", Millipore-Sigma) with 3 µL of β-mercaptoethanol and adding 50 µL of this mixture to each well for measurement. 50 µL of each supernatant from the quenched reactions and standard curves were then transferred to the appropriate wells, pipetted once to mix, and then fluorescence was immediately measured on a plate reader (Bio-Tek Synergy HTX multimode plate reader, measurement settings – 360/40 nm excitation filter, 440/20 nm emission filter, detector gain: 35). Fluorescence was measured every 30 seconds for 10 minutes and the maximum value achieved for each well was used for calculations. Initial rate measurements are based on 3 independent replicates at each substrate concentration and 5 time points spaced 1 minute apart.

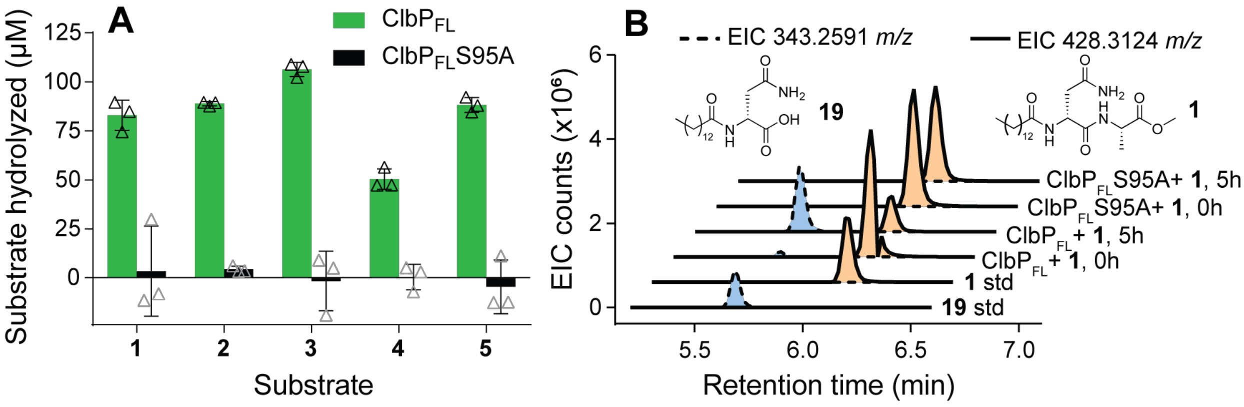

**Figure S3** – Cleavage of substrates with varying $R_1$ groups (**1**–**5**) by $ClbP_{FL}$. (a) Bars represent average amount of substrate consumed across 3 replicates after 5 hours. Concentrations were determined by comparing peak areas of the $[M+H]^+$ ions of each substrates to a standard curve generated for each substrate (see methods below. Triangles represent individual replicate values, error bars = 1 SD). (b) Representative EICs for $[M+H]^+$ ion masses of substrate **1** and its $ClbP_{FL}$ cleavage product, **19** from a typical reaction at the start of the reaction ("0h") and after 5 hours at room temperature ("5h"). **1**: 428.3124 *m/z*; **19**: 343.2597 *m/z*

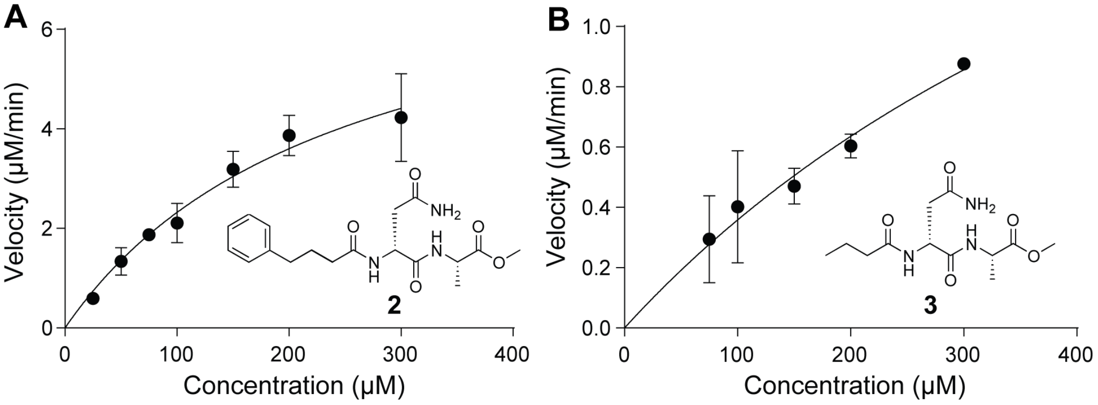

**Figure S4** –Kinetics of $ClbP_{FL}$-mediated cleavage of substrates (a) **2** and (b) **3**. Each data point is an average of 3 biological replicates with error bars representing 1 SD. Solid lines are non-linear regression fits of data points to the Michaelis-Menten model.

**Table S1** – Kinetic parameters for substrates **1**, **2**, and **3**.

| | $k_{cat}$ (min$^{-1}$) | $K_M$ (μM) | $k_{cat}/K_M$ (M$^{-1}$s$^{-1}$) |
|---|---|---|---|
| **1** | 36 ± 4 | 130 ± 26 | 4600 ± 1100 |
| **2** | 80 ± 13 | 246 ± 70 | 5400 ± 1800 |
| **3** | 28 ± 15 | 680 ± 470 | 690 ± 600 |

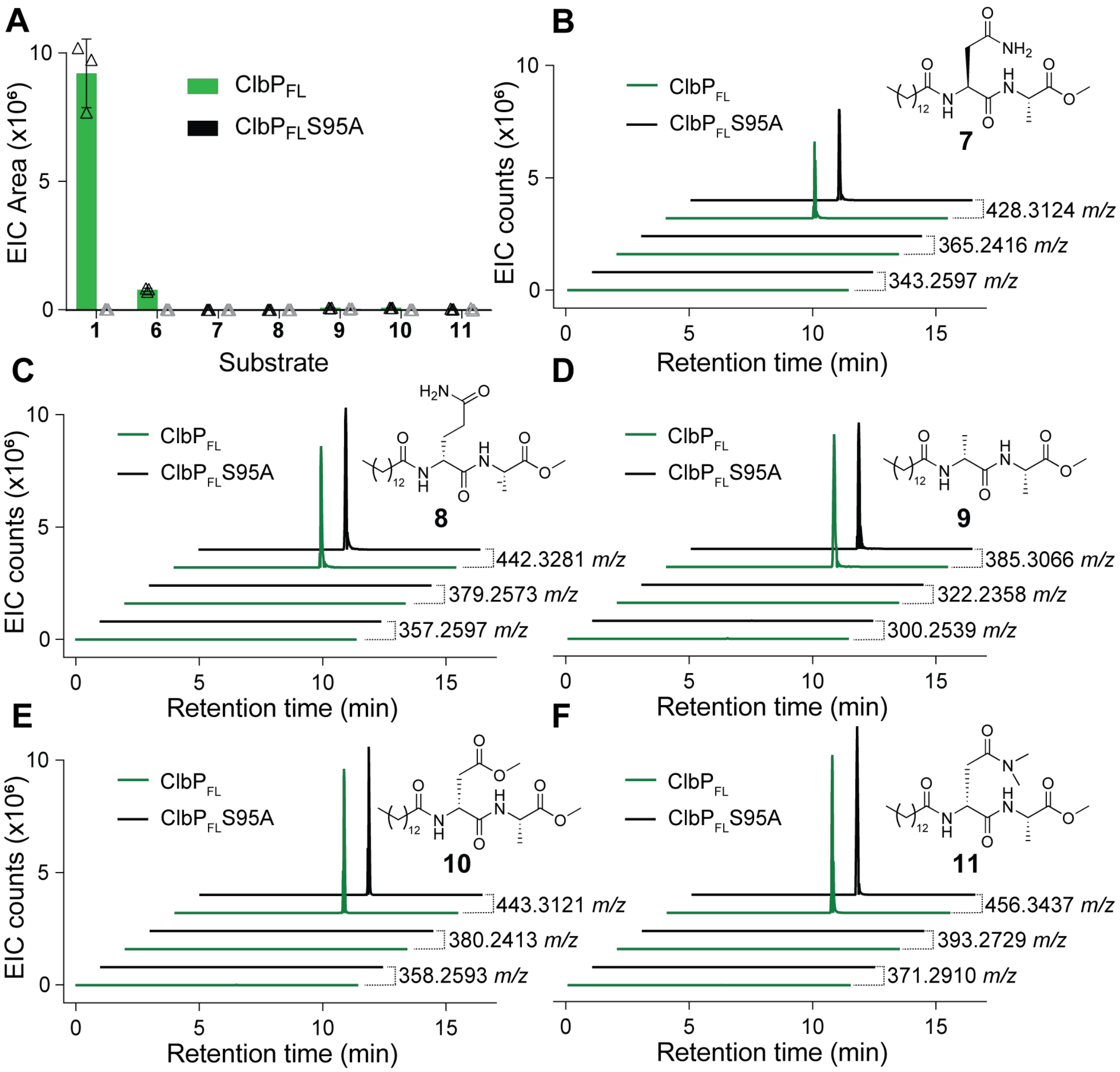

**Figure S5** – Cleavage of substrates with varying $R_2$ groups (**1, 6–11**) by $ClbP_{FL}$. (a) Bars represent average EIC peak areas for to the $[M+H]^+$ ions of the expected cleavage products: **1**: 343.2597 *m/z*; **6**: 344.2437 *m/z*; **7**: 343.2597 *m/z*; **8**: 357.2753 *m/z*; **9**: 300.2539 *m/z*; **10**: 358.2593 *m/z*; **11**: 371.2910 *m/z*; All substrates were tested in triplicate; triangles show individual replicate values (error bars = 1 SD) (b-f) Representative EIC traces (1 replicate each) for masses of substrates and expected cleavage products ($[M+H]^+$ and $[M+Na]^+$ for products, $[M+H]^+$ for substrates) for each of the substrates which showed no cleavage by $ClbP_{FL}$ *in vitro* (substrates **7** – **11**). All EICs are for reactions quenched after incubation at room temperature for 5 hours (compare to Figure S2b).

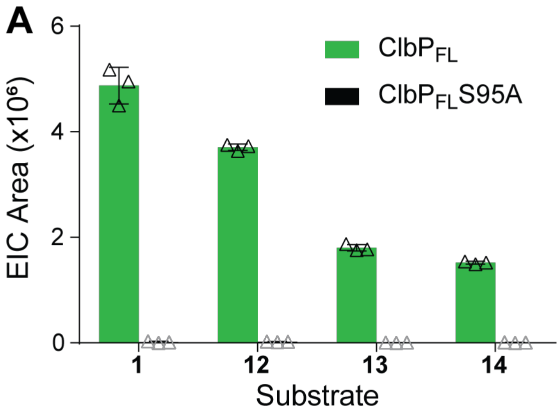

**Figure S6** – Cleavage of substrates with varying $R_3$ groups (**12**–**14**) by $ClbP_{FL}$. Bars represent average EIC peak areas for the $[M+H]^+$ ion of the expected cleavage product, **19** (343.2597 *m/z*) which is the same for these four substrates. All substrates were tested in triplicate; triangles show individual replicate values (error bars = 1 SD).

### 5. ClbP activity assays with fluorogenic probes

**Initial validation of probes 15 *in vitro* (Figure S6a):** Probes **15** was prepared as 10 mM stock solutions in DMSO and 1 µL of the appropriate stock was added to individual wells of an opaque black 96-well plate. $ClbP_{FL}$-$CHis_{10}$ and $ClbP_{FL}$-S95A-$CHis_{10}$ were defrosted from –80 °C stocks on ice, then diluted to 0.2 µM in Tris assay buffer. To initiate reactions, 99 µL of either $ClbP_{FL}$ or $ClbP_{FL}$-S95A were transferred to the 96-well plate via multichannel pipette and mixed (n = 6 for each enzyme). Activity was monitored by measuring the fluorescence of each well once per minute for 2 hours using a plate reader (Bio-Tek Synergy HTX multimode plate reader, measurement settings – 360/40 nm excitation filter, 440/20 nm emission filter, detector gain: 35).

**Validation of probes 15 and 16 whole-cells overexpressing $ClbP_{FL}$ (Figure 3c and Figure S6b):** A 5 mL starter culture of *E. coli* BL21 pET-29b-$ClbP_{FL}$ or pET-29b-$ClbP_{FL}$-S95A was inoculated from a frozen stock and grown overnight at 37 °C in LB medium supplemented with 50 µg/ml kanamycin. Overnight cultures were diluted 1:100 into 5 mL M9 minimal medium supplemented with 1% w/v Casamino acids (“M9+CAA”) containing 50 µg/mL kanamycin. Cultures were incubated at 37 °C with shaking at 175 rpm. Protein expression was induced with 500 µM IPTG at $OD_{600}$ = 0.5 and cultures were moved to 15 °C and incubated with shaking at 175 RPM for ~ 4 h. The $OD_{600}$ of each culture was measured again. Cultures were pelleted at 4000 x *g* for 10 minutes at 4 °C and resuspended in enough fresh M9+CAA media (containing 50 µg/mL kanamycin and 500 µM IPTG) to normalize the $OD_{600}$ to 0.5. Resuspended cultures were added to a black 96 well plate, 120 µL per well. Probe **15** or **16** was prepared in a master mix of 1 mM probe, 50 µg/mL kanamycin, and 500 µM IPTG in M9+CAA media. 30 µL of master mix was added to each well to achieve a final concentration of 200 µM probe (2% DMSO). Plates were incubated at room temperature in a plate reader overnight while taking fluorescence measurements at regular intervals (Bio-Tek Synergy HTX multimode plate reader, measurement settings – 360/40 nm excitation filter, 440/20 nm emission filter, detector gain: 35).

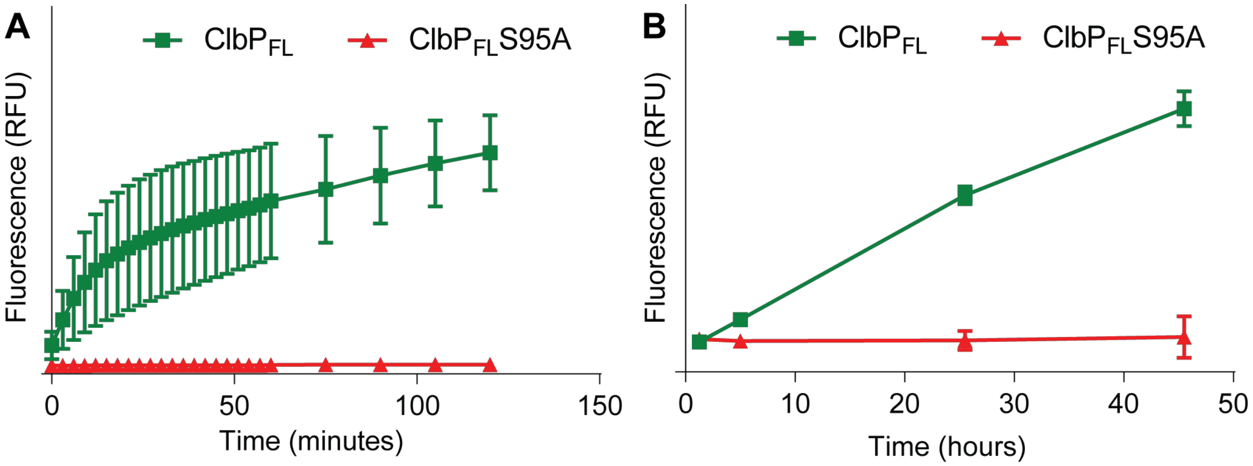

**Figure S7** – (a) detection of ClbP$_{FL}$ activity *in vitro* using probe **15**. (b) Detection of ClbP$_{FL}$ activity in *E. coli* BL21 pET29b_ClbP$_{FL}$ using probe **15**. Data points are averages of 6 biological replicates (error bars = 1 SD).

**Comparing activity of ClbP$_{FL}$ and ClbP$_{pep}$ *in vitro* (Figure 3b):** Probe **16** was prepared as a 10 mM stock solution in DMSO then diluted to 250 µM in standard Tris assay buffer, and 5 µL of this stock was added to each well of an opaque black 384-well flat-bottom plate. ClbP$_{FL}$-CHis$_{10}$ and ClbP$_{FL}$-S95A-CHis$_{10}$ were defrosted from –80 °C stocks on ice, then diluted to 0.125 µM in Tris assay buffer. Reactions were initiated by added 20 µL of either the ClbP$_{FL}$ or ClbP$_{FL}$-S95A to the wells via multichannel pipette and pipetting once to mix (n = 6 for each enzyme). Assays for ClbP$_{pep}$ and ClbP$_{pep}$-S95A activity were conducted in parallel following the same procedure, but in buffers that did not contain DDM. All reactions contained 0.1 µM enzyme, 50 µM substrate, and 0.5% DMSO and were run at 25 °C. Activity was monitored by measuring the fluorescence of each well once per minute for 2 hours using a plate reader (Bio-Tek Synergy HTX multimode plate reader, measurement settings – 360/40 nm excitation filter, 440/20 nm emission filter, detector gain: 35).

**Detection of ClbP activity in *pks*$^+$ organisms (Figure 4a)**: For anaerobic experiments, MEGA media[11] was autoclaved for 20 minutes at 120 °C, sparged with sterile $N_2$ for 1 hour, and equilibrated in a Coy vinyl anaerobic chamber (Coy Labs, Grass Lake, MI) overnight under a 95% $N_2$/5% $H_2$ atmosphere. All anaerobic experiments were performed in this environment. Note that the standard formulation of MEGA medium includes resazurin as a colorimetric oxygen indicator, but this was omitted to avoid interfering with the fluorescence signal. A 5 mL starter culture of *E. coli* BW25113 BAC*pks*, BW25113 BAC vector, Nissle 1917, or CCR20 was inoculated from a frozen stock and grown overnight at 37 °C in MEGA medium under anaerobic conditions without shaking. Overnight cultures were diluted 1:100 into 1 mL of MEGA medium containing 100 µM **16** (added as a 10 mM stock in DMSO) in triplicate in a 1.5 mL 96-well plate. Cultures were incubated anaerobically at 37 °C. At each time point, 100 µL of each culture was transferred to an opaque black 96-well plate, removed from the anaerobic chamber, and the fluorescence was measured immediately (Bio-Tek Synergy HTX multimode plate reader, measurement settings – 360/40 nm excitation filter, 440/20 nm emission filter, detector gain: 35).

**Testing inhibition of ClbP$_{FL}$ by small molecules using the fluorescence-based assay (Figure 4b)**: Boronic acids **17** and **18** were prepared as 50 mM stocks in DMSO, while probe **16** was prepared as a 10 mM stock in DMSO. ClbP$_{FL}$-CHis$_{10}$ was defrosted on ice from –80 °C stocks, diluted to 0.125 µM in Tris assay buffer, and incubated with the appropriate concentration of **17, 18**, or DMSO (controls) for 1 hour at ambient temperature in order to allow for inhibitor binding (n = 3 for each concentration). Separately, 250 µM **1** in Tris assay buffer was deposited in the wells of an opaque black 384-well flat-bottom plate (5 µL/well). Reactions were initiated by transferring 20 µL of the enzyme/inhibitor or enzyme/DMSO mix to the 384 well plate (final assay conditions: 0.1 µM enzyme, 50 µM probe **16**, 0-2 mM inhibitor, 4.5% DMSO, 25 °C). Activity was monitored by measuring the fluorescence of each well once per minute for 4 hours using a plate reader (Bio-Tek Synergy HTX multimode plate reader, measurement settings – 360/40 nm excitation filter, 440/20 nm emission filter, detector gain: 35).

**Testing inhibition of ClbP$_{FL}$ by small molecules using the LC-MS-based assay (Figure S7)**: Boronic acids **17** (purchased from Sigma Aldrich) and **18** (gifted from Professor Fabio Prati, see Synthetic Methods section for details) were prepared as 50 mM stocks in DMSO, while substrate **1** was prepared as a 10 mM stock in DMSO. ClbP$_{FL}$-CHis$_{10}$ was defrosted on ice from –80 °C stocks, diluted to 0.12 µM in Tris assay buffer, and incubated with the appropriate concentration of **17**, **18**, or DMSO (controls) for 1 hour at ambient temperature in order to allow for inhibitor binding. Separately, 300 µM **1** in Tris assay buffer was deposited in the wells of a 96-well plate (25 µL). Reactions were initiated by transferring 125 µL of the enzyme/inhibitor or enzyme/DMSO mix to the 96-well plate (final assay conditions: 0.1 µM enzyme, 50 µM substrate **1**, 0-2 mM inhibitor, 4.5% DMSO, 25 °C). Time points were taken at 0, 5, and 10 minutes as described above (see "SAR study") and LC-MS samples were prepared in the same manner. For each condition, a reaction velocity was determined by measuring the increase in EIC peak area for the expected mass of the prodrug scaffold fragment (±5 ppm) independently across three replicates.

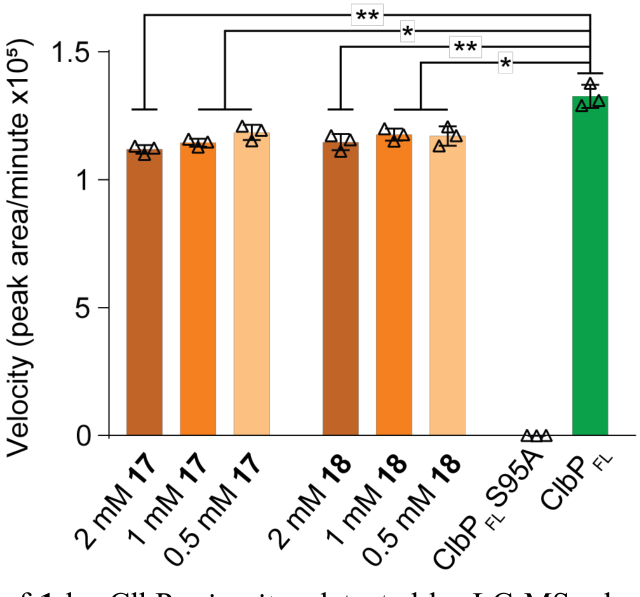

**Figure S8** – Cleavage of **1** by ClbP$_{FL}$ *in vitro* detected by LC-MS when treated with **17** or **18.** Reaction velocities correspond to the increase in EIC peak area for the $[M+H]^+$ ion of **19** (343.3597 *m/z*). Experiments were performed in triplicate at each concentration for each molecule (triangle represent individual replicates). Bars represent the slope of the linear fit of increase in the EIC peak area of **19** of over the first 10 minutes of the reaction of ClbP$_{FL}$ *in vitro* with **1**. Error bars represent SD of the slope values. * : $P < 0.05$; ** : $P < 0.01$; Student's two-tailed *t* test.

### 6. Chemical synthesis procedures and characterization data

Compounds **1**, **14**, **19**, and intermediate **28** were synthesized as previously described.[10] NMR spectra and high-resolution masses for all compounds were identical to those previously reported. Inhibitor **18** was provided as a gift from Professor Fabio Prati (Unimore, Via Università 4, 41121 Modena, Italy) and was prepared as previously described.[12] Its purity was confirmed by $^1$H NMR spectroscopy.

† **Note on NMR spectra:** For many of the substrates below which contain a D-asparagine residue, a peak annotated as a doublet of doublets is reported at or near 2.33 ppm with coupling constants of approximately $J$ = 15.3 and 8.1 Hz, a 1H integration, and an asymmetric splitting pattern. This peak is actually one of two peaks originating from the two diastereotopic protons on the β-carbon of the D-asparagine side chain. The other 1H proton signal overlaps with the DMSO-$d_6$ solvent residual for some substrates. When possible, $^1$H NMR spectra were recorded in $CDCl_3$ or $CD_3OD$ to avoid this issue, but many of these substrates are poorly soluble in anything other than DMSO. For several example substrates, 2D NMR spectroscopy experiments ($^1$H-$^{13}$C HSQC, HMBC or $^1$H-$^1$H COSY) were used to confirm that there is a second peak overlapping with the DMSO signal and that the connectivity is as described (see substrates **3**, **5**, **6**, **11**, **12**, **15**, and **16** for example spectra). The chemical shift and splitting patterns for these peaks are also in line with computational predictions for this peak. This overlap results in the total integration of the spectrum lacking 1 H when compared to the molecular formula. In all cases where this overlap occurs, the peak is denoted by a †.

#### General Procedure A: HATU coupling of amino acids

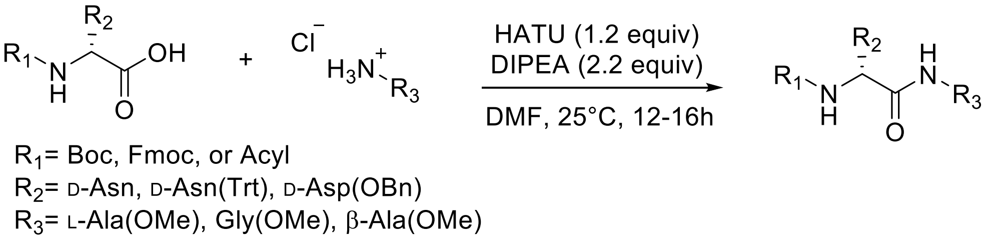

To an oven-dried round bottom flask equipped with a stir bar was added a Boc-protected, Fmoc-protected, or $N$-acylated amino acid (1.0 equiv), the HCl-salt of the methyl ester-protected amino acid (1.2 equiv), and hexafluorophosphate azabenzotriazole tetramethyluronium (HATU, 1.2 equiv, Oakwood Chemicals or Chem-Impex International, Inc.) under $N_2$. Anhydrous $N,N$-dimethylformamide (DMF, 0.1–0.5 M) was added and the reaction mixture stirred at room temperature until all components were dissolved. Diisopropylethylamine (DIPEA, 2.2 equiv) was added dropwise, and the reaction mixture was stirred at room temperature overnight. The reaction was quenched by the addition of 1 reaction volume of aqueous 1 M HCl. The reaction mixture was diluted with 5 reaction volumes of ethyl acetate (EtOAc) and the organic layer was washed with 1 M HCl. The aqueous layer was extracted a second time with ethyl acetate. The combined organic layers were washed with 5% aqueous LiCl, saturated aqueous sodium bicarbonate, water, and brine. The organic layer was dried over $Na_2SO_4$, filtered, and concentrated *in vacuo* to give the final product. The crude product was either used directly in the subsequent step or was purified as described for the individual compounds listed below.

**General Procedure B: HATU coupling of dipeptides to acyl group**

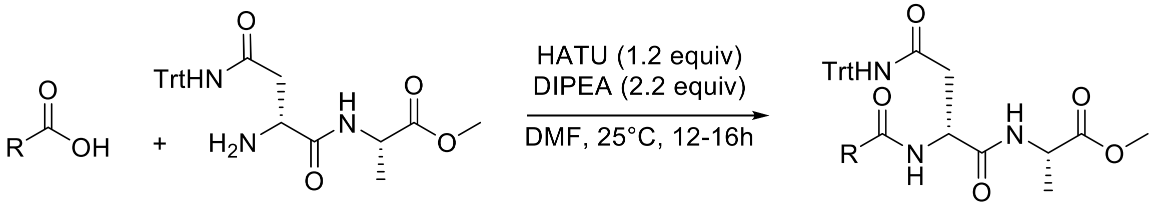

R = Acyl

To an oven-dried round bottom flask equipped with a stir bar was added a dipeptide methyl ester (1.0 equiv), the appropriate carboxylic acid (1.2 equiv), and hexafluorophosphate azabenzotriazole tetramethyluronium (HATU, 1.2 equiv, Oakwood Chemicals or Chem-Impex International, Inc.) under $N_2$. Anhydrous DMF (0.1–0.5 M) was added and the reaction mixture was stirred at room temperature until all components were dissolved. DIPEA (2.2 equiv) was added dropwise, and the reaction mixture was stirred at room temperature overnight. The reaction was quenched by the addition of 1 reaction volume of aqueous 1 M HCl. The reaction mixture was then diluted with 5 reaction volumes of EtOAc and the organic layer was washed with 1 M HCl. The aqueous layer was extracted a second time with ethyl acetate. The combined organic layers were washed with 5% aqueous LiCl, saturated aqueous $NaHCO_3$, water, and brine. The organic layer was dried over $Na_2SO_4$ and concentrated *in vacuo* to give the crude product. The crude product was used directly in the subsequent step.

**General Procedure C: Acyl chloride coupling**

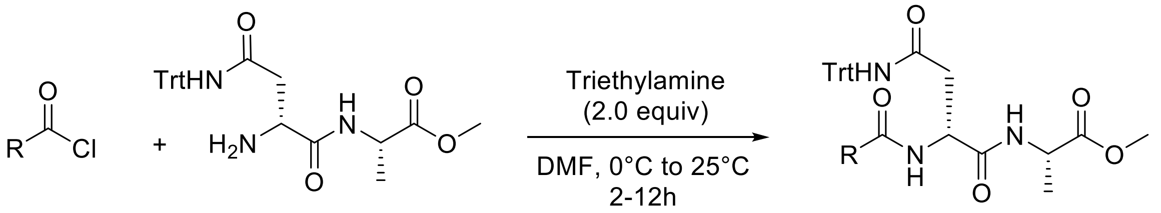

R = Acyl

To an oven-dried round bottom flask equipped with a stir bar was added the dipeptide methyl ester (1.0 equiv) and anhydrous DMF (0.4 M) under dry $N_2$. Triethylamine (TEA, 2.0 equiv) was added and the mixture was subsequently cooled to 0 °C. In a separate oven-dried round bottom flask, the acyl chloride (2.0 equiv) was diluted in anhydrous DMF (0.8 M) under dry $N_2$ and cooled to 0 °C. The acyl chloride solution was added dropwise to the flask containing the dipeptide via syringe. The reaction mixture was warmed to room temperature and stirred for at least 2 hours and up to 12 hours. Work up conditions varied for each substrate and are described for the individual compounds listed below. The crude product was used directly in the subsequent step.

**General Procedure D: Trityl deprotection**

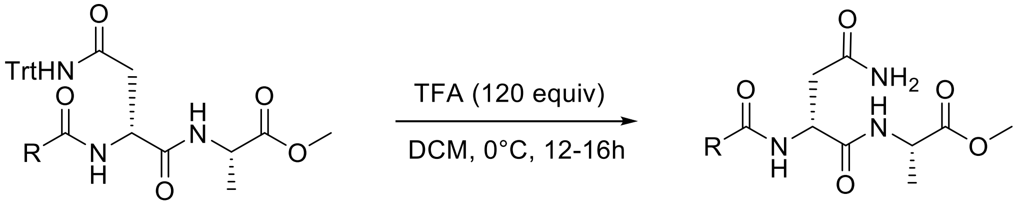

R = Acyl

To a round bottom flask was added the trityl-protected peptide and dichloromethane (DCM, 0.05–0.2 M). Once the starting material was dissolved, the reaction mixture was cooled to 0 °C and trifluoroacetic acid (TFA, 120 equiv) was added. The flask was sealed and kept at 4 °C overnight. Toluene (1 reaction volume) was added, and the reaction mixture was concentrated *in vacuo*. This procedure was repeated twice more to remove any residual TFA. The crude residue was dissolved in DMSO and purified using preparative HPLC (conditions are specified for each substrate below).

**General Procedure E: PyBOP coupling**

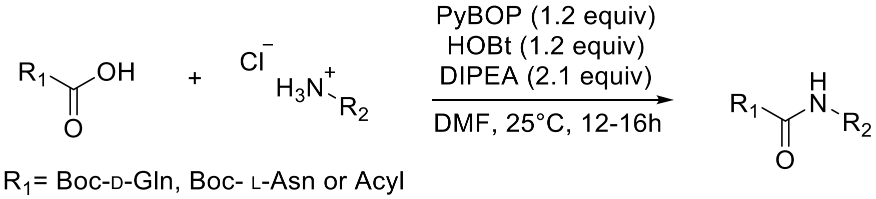

To an oven-dried round bottom flask equipped with a stir bar was added a Boc-protected amino acid or carboxylic acid (1.0 equiv), the HCl-salt of the methyl ester-protected amino acid (1.0 equiv) and dry DMF (0.1–0.3 M). The starting materials were dissolved at room temperature and HOBt (Sigma Aldrich, 1.2 equiv) and PyBOP (Chem-Impex International, Inc) were added. The flask was then cooled to 0 °C and DIPEA (2.1 equiv) was added dropwise. The reaction mixture was warmed to room temperature overnight. The reaction mixture was then diluted with 5 reaction volumes of EtOAc or DCM and quenched by the addition of 1 reaction volume of aqueous 1 M HCl. The aqueous and organic layers were separated and the aqueous layer was extracted 2 more times with the same solvent (EtOAc or DCM). The combined organic layers were washed with saturated aqueous $NaHCO_3$ and brine. The organic layer was dried over $Na_2SO_4$, filtered, and then concentrated *in vacuo.* The crude product was either used directly in the subsequent step or was purified as described for the individual compounds listed below.

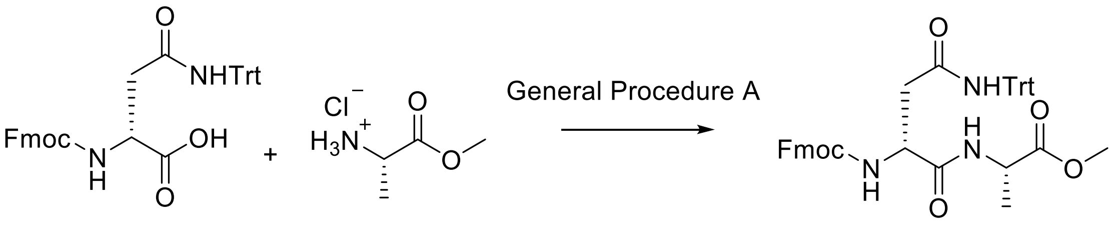

**20**

Dipeptide **20** was accessed by coupling *N*α-Fmoc-*N*γ-Trt-D-asparagine (Chem-Impex International, 2.13 g) and L-alanine methylester hydrochloride (Chem-Impex International, 547 mg) according to General Procedure A. This intermediate was purified by flash column chromatography (silica, isocratic 6:2:1 EtOAc:Hexanes:DCM) affording **20** as a white solid (2.2 g, 85% yield). $^{1}$H NMR (600 MHz, $CDCl_3$): δ (ppm) = 7.76 (d, *J* = 7.6 Hz, 2H), 7.58 (t, *J* = 7.7 Hz, 2H), 7.40 (t, *J* = 7.5 Hz, 2H), 7.34 – 7.22 (m, 15H)*, 7.19 (d, *J* = 7.6 Hz, 6H), 6.90 (s, 1H), 6.40 (d, *J* = 7.9 Hz, 1H), 4.57 (s, 1H), 4.46 (q, *J* = 7.1 Hz, 1H), 4.44 – 4.38 (m, 1H), 4.35 (t, *J* = 9.1 Hz, 1H), 4.21 (t, *J* = 7.2 Hz, 1H), 3.71 (d, *J* = 1.1 Hz, 3H), 3.21 – 3.03 (m, 1H), 2.59 (dd, *J* = 15.7, 6.1 Hz, 1H), 1.35 (d, *J* = 7.1 Hz, 3H). $^{13}$C NMR (400 MHz, DMSO-$d_6$): δ (ppm) = 172.8, 171.1, 168.66, 155.6, 144.7, 143.8, 140.7, 128.6, 127.6, 127.4, 127.1, 126.3, 125.33, 125.28, 120.1,

69.4, 65.8, 51.9, 47.7, 46.7, 17.1‡. HRMS (ESI): calcd for $C_{42}H_{40}N_3O_6$ $[M+H]^+$, 682.2917; found, 682.2945.

*Additional 3H integration value due to overlap with solvent residual with expected 12H multiplet
‡An additional $^{13}C$ peak is predicted at a shift of 80.7 ppm, corresponding to the quaternary carbon of the trityl protecting group. This signal could not be detected, likely due to the poor signal produced by this nuclide and the limited solubility of **20**.

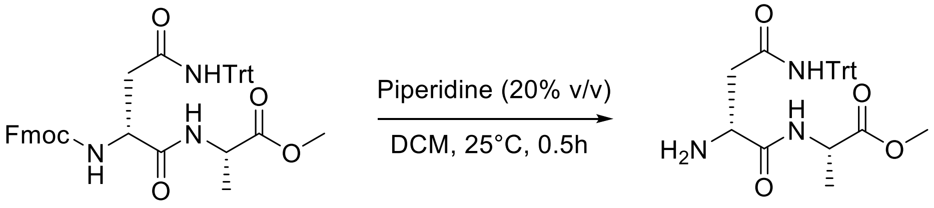

**20** **21**

Fmoc-protected dipeptide **20** (105 mg) was dissolved in DCM (1.5 mL, 0.1 M) in a round bottom flask. Piperidine (0.38 mL) was added and the reaction mixture was stirred at room temperature under an atmosphere of $N_2$ for approximately 30 minutes until thin-layer chromatography (TLC) indicated that the reaction was complete. The reaction mixture was diluted with 5 reaction volumes of DCM and then quenched by the addition of 5 reaction volumes of saturated aqueous ammonium chloride. The aqueous layer was extracted again with DCM, and the combined organic layers were washed with saturated aqueous $NH_4Cl$, $NaHCO_3$, and brine. The organic layer was dried over $Na_2SO_4$, filtered, and then concentrated *in vacuo*. The resulting residue was purified by flash column chromatography (gradient from $CHCl_3$ to $CHCl_3$:MeOH 9:1 on neutral alumina) affording **21** as a thick clear oil (73 mg, 70% yield). $^1H$ NMR (500 MHz, $CDCl_3$): δ (ppm) = 7.82 (d, $J$ = 7.6 Hz, 1H), 7.59 (s, 1H), 7.36 – 7.18 (m, 17H), 4.47 (quint, $J$ = 7.3 Hz, 1H), 3.71 (s, 3H), 3.66 (dd, $J$ = 8.1, 4.1 Hz, 1H), 2.73 (dd, $J$ = 15.1, 4.1 Hz, 1H), 2.62 (dd, $J$ = 15.1, 8.2 Hz, 1H), 1.38 (d, $J$ = 7.2 Hz, 3H). $^{13}C$ NMR (400 MHz, DMSO-$d_6$): δ (ppm) = 173.2*, 172.4, 168.7, 168.6*, 167.8, 144.6§, 144.4, 144.38*, 143.7§, 128.6*, 128.5, 128.2§, 127.9§, 127.54, 127.47*, 127.0§, 126.5, 126.4*, 69.64, 69.56*, 52.3*, 52.1, 47.9, 47.8*, 36.2, 17.1, 17.0*.‡ HRMS (ESI): calcd for $C_{27}H_{30}N_3O_4$ $[M+H]^+$, 460.2236; found, 460.2233.

* denotes a rotamer

§ denotes a peak arising from residual dibenzofulvene, the side product of Fmoc-deprotection

‡An additional peak is predicted at a shift of 80.7 ppm, corresponding to the quaternary carbon of the trityl protecting group. This signal could not be detected, likely due to the poor signal produced by this nuclide.

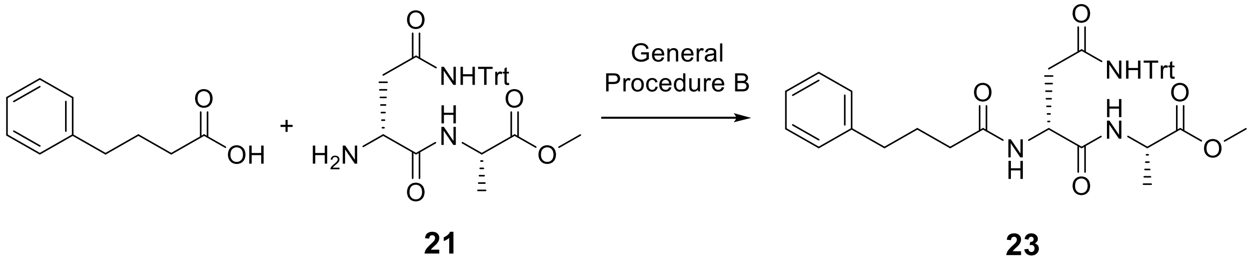

Dipeptide **21** (122 mg) was coupled to 4-phenylbutanoic acid (48 mg) according to General Procedure B. The product **23** was isolated as a white solid (145 mg, 90% yield). Crude **23** was used in the next step of the synthesis (trityl deprotection) without further purification.

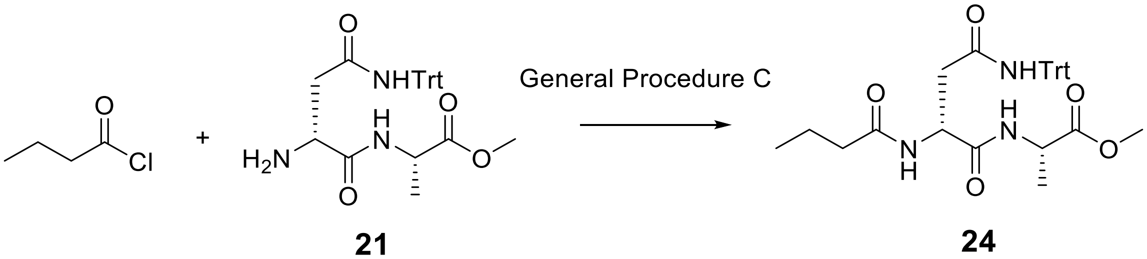

Dipeptide **21** (172 mg) was coupled to butyryl chloride (77 µL) following General Procedure C. The reaction mixture was diluted with 5 reaction volumes of DCM and washed with 5 reaction volumes of aqueous 0.1 M HCl. The aqueous layer was extracted again with DCM and the combined organic layers were washed with saturated aqueous $NaHCO_3$, water, and brine. The organic layer was dried over $Na_2SO_4$ and was then concentrated *in vacuo* to afford crude **24** as a light-yellow solid (101 mg, 51% yield). This material was used in the next step of the synthesis (trityl deprotection) without further purification.

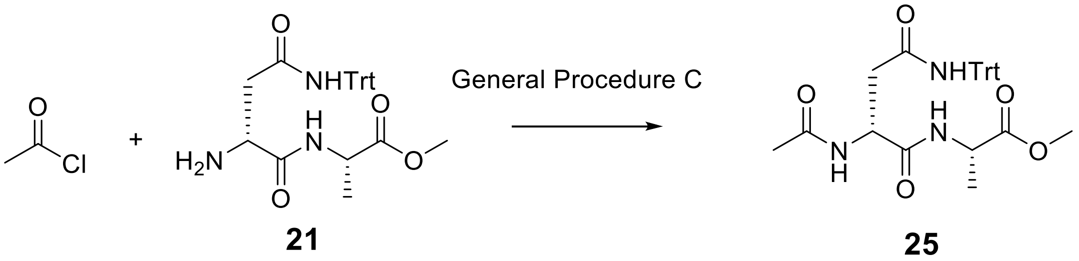

Dipeptide **21** (53 mg) was coupled to acetyl chloride (9 µL) following General Procedure C. The reaction mixture was diluted with EtOAc and washed with aqueous 0.1 M HCl. The aqueous layer was extracted again with EtOAc, and the combined organic layers were washed with saturated aqueous $NaHCO_3$, water, and brine. The organic layer was dried over $Na_2SO_4$ and was then concentrated *in vacuo* to afford crude **25** as a white solid (56.4 mg, 97% yield). Crude **25** was used in the next step of the synthesis (trityl deprotection) without further purification.

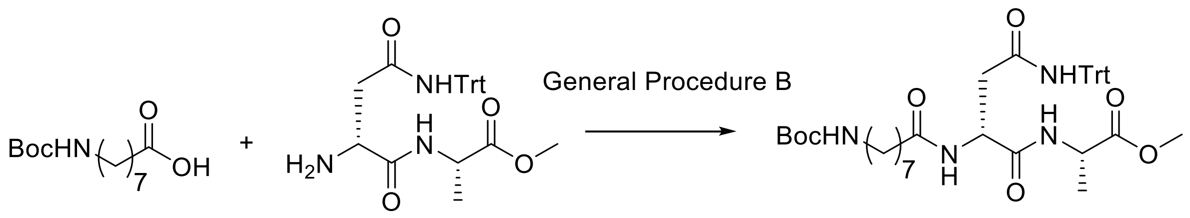

**21** **26**

Dipeptide **21** (50 mg) was coupled to 8-aminocaproic acid (28.2 mg) following General Procedure B. The crude product, **26**, was isolated as an off-white solid (yield reported for next step). Crude **26** was used in the next step of the synthesis (trityl deprotection) without further purification.

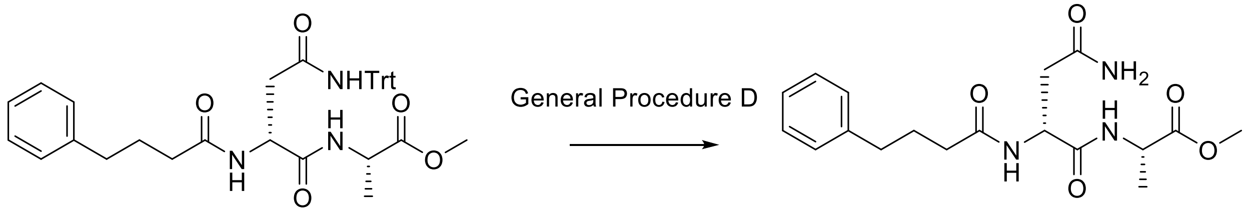

**23** **2**

**23** was deprotected to afford **2** according to General Procedure D. The initial product of the deprotection was a waxy off-white solid. This solid was dissolved in a minimal volume of DMSO and purified by preparative HPLC following the procedure described in the General Materials and Methods. Fractions containing product were pooled and concentrated *in vacuo* to afford **2** as a white solid (38.2 mg, 57.9% yield over two steps). $^{1}$H NMR (500 MHz DMSO-$d_6$): δ (ppm) = 8.10 (d, $J$ = 7.3 Hz, 1H), 7.97 (d, $J$ = 8.1 Hz, 1H), 7.31 – 7.14 (m, 6H), 6.86 (s, 1H), 4.60 (dt, $J$ = 8.1, 5.8 Hz, 1H), 4.25 (quint, $J$ = 7.2 Hz, 1H), 3.59 (s, 3H), 2.55 (t, $J$ = 7.5 Hz, 2H), 2.33† (dd, $J$ = 15.3, 8.1 Hz, 1H), 2.13 (t, $J$ = 7.4 Hz, 2H), 1.78 (quint, $J$ = 7.5 Hz, 2H), 1.25 (d, $J$ = 7.3 Hz, 3H). $^{13}$C NMR (500 MHz, DMSO-$d_6$): δ (ppm) = δ 172.8, 171.9, 171.2, 171.0, 141.9, 128.3, 128.2, 125.7, 51.8, 49.4, 47.6, 37.3, 34.7, 34.5, 27.1, 17.1. HRMS (ESI): calcd for $C_{18}H_{26}N_3O_5$ $[M+H]^+$, 364.1872; found, 364.1870. $[\alpha]_D^{22}$ = + 7.4° ($c$ = 0.84, DMSO)

† see “Note on NMR spectra” at the beginning of this section

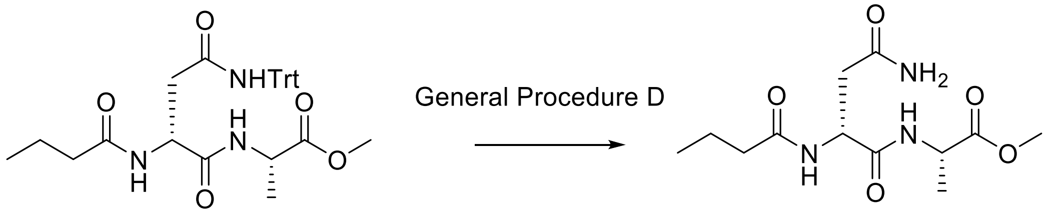

**24** **3**

**24** was deprotected to afford **3** according to General Procedure D. The initial product of the deprotection was an off-white solid. This solid was dissolved in a minimal volume of DMSO and purified by preparative HPLC using the following gradient conditions: 10% Solvent A for 2 minutes, gradient to 95% Solvent A over 7 minutes, hold at 95% Solvent A for 4 minutes, gradient to 10% solvent A over 1 minute, hold at 10% solvent A for 2 minutes (solvent A: HPLC-grade acetonitrile + 0.1% formic acid; solvent B: water + 0.1% formic acid; flow rate: 10 mL/minute; injection volume: 200 to 400 μL). Fractions were pooled and concentrated *in vacuo* to afford **3** as a white solid (30 mg, 27% yield over two steps). $^{1}$H NMR (600 MHz DMSO-$d_6$): δ (ppm) = 8.12 (d, $J$ = 7.3 Hz, 1H), 8.01 (d, $J$ = 8.0 Hz, 1H), 7.29 (s, 1H), 6.85 (s, 1H), 4.59 (dt, $J$ = 8.1, 5.8 Hz, 1H), 4.25 (quint, $J$ = 7.2 Hz, 1H), 3.61 (s, 3H), 2.48 (dd, $J$ = 15.2, 5.9 Hz, 1H), 2.34 (dd, $J$ = 15.2, 5.9 Hz, 1H), 2.08 (t, $J$ = 7.3 Hz, 2H), 1.50 (sext, 2H), 1.25 (d, $J$ = 7.2 Hz, 3H), 0.85 (t, $J$ = 7.4 Hz, 3H).$^{13}$C NMR (500 MHz, DMSO-$d_6$) δ = 172.8, 172.0, 171.3, 171.0, 51.9, 49.4, 47.6, 37.4, 37.1, 18.6, 17.1, 13.5. HRMS (ESI): calcd for $C_{12}H_{22}N_3O_5$ [M+H]$^+$, 288.1559; found, 288.1546. $[\alpha]_D^{23}$ = + 23.4° ($c$ = 1.0, DMSO)

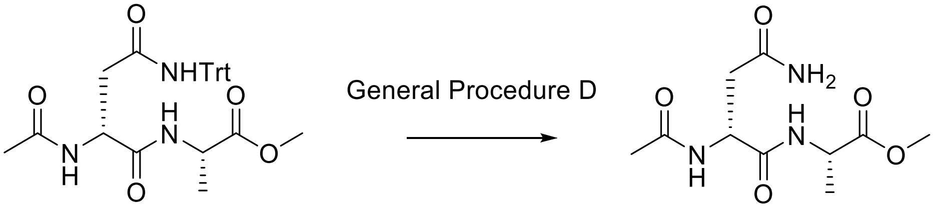

**25** **4**

**25** was deprotected to afford **4** according to General Procedure D. The initial product of the deprotection was an off-white solid. This solid was dissolved in a minimal volume of DMSO and purified by preparative HPLC using the following gradient conditions: 5% Solvent A for 2 minutes, gradient to 40% Solvent A over 8 minutes, gradient to 95% Solvent A over 5 minutes, hold at 95% Solvent A for 2 minutes, gradient to 5% solvent A over 1 minute, hold at 5% solvent A for 2 minutes (solvent A: HPLC-grade acetonitrile + 0.1% formic acid; solvent B: water + 0.1% formic acid; flow rate: 6 mL/minute; injection volume: 200 to 400 µL). Fractions were pooled and concentrated *in vacuo* to afford **4** as a white solid (24 mg, 17% yield over two steps). $^{1}$H NMR (500 MHz DMSO-$d_6$): δ (ppm) = 8.17 (d, $J$ = 7.3, 1H), 8.04 (d, $J$ = 7.6 Hz, 1H), 7.29 (s, 1H), 6.87 (s, 1H), 4.57 (dt, $J$ = 8.0, 5.7 Hz, 1H), 4.24 (quint, $J$ = 7.2 Hz, 1H), 3.61 (s, 3H), 2.47 (dd, $J$ = 15.5, 6.1 Hz, 1H), 2.32 (dd, $J$ = 15.2, 8.1 Hz, 1H),1.83 (s, 3H), 1.25 (d, $J$ = 7.2 Hz, 3H).$^{13}$C NMR (500 MHz, DMSO-$d_6$) δ = 172.8, 171.3, 171.0, 169.2, 51.9, 49.5, 47.6, 37.5, 22.6, 17.01. HRMS (ESI): calcd for $C_{10}H_{18}N_3O_5$ [M+H]$^+$, 260.1246; found, 260.1235. $[\alpha]_D^{21.5}$ = + 15.0° ($c$ = 0.2, DMSO)

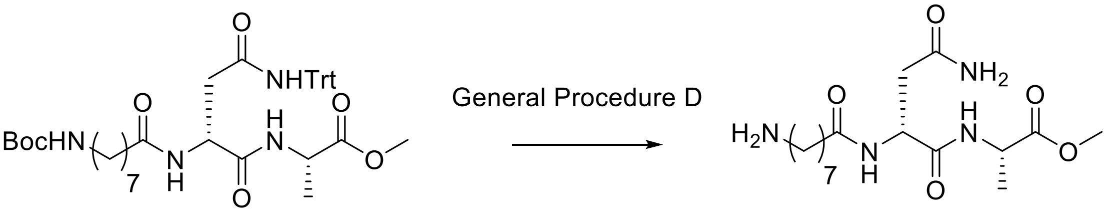

**26** **5**

**26** was deprotected to afford **5** according to General Procedure D. The initial product of the deprotection was an off-white solid. This solid was dissolved in a minimal volume of DMSO and purified by preparative HPLC using the following gradient conditions: 5% Solvent A for 2 minutes, gradient to 45% Solvent A over 7 minutes, hold for 2 minutes, gradient to 60% solvent A over 2 minutes, hold for 3 minutes, gradient to 95% solvent A over 2 minutes, hold for 1 minute, gradient to 5% solvent A over 1 minute, hold for 3 minutes (solvent A: HPLC-grade acetonitrile + 0.1% formic acid; solvent B: water + 0.1% formic acid; flow rate: 7 mL/minute; injection volume: 200 to 400 µL). Fractions were pooled and concentrated *in vacuo* to afford **5** as a white solid (34 mg, 40% yield over two steps). $^{1}$H NMR (400 MHz DMSO-$d_6$): δ (ppm) = 8.20 – 7.94 (m, 2H), 7.32 (s, 1H), 6.86 (s, 1H), 4.57 (d, $J$ = 7.1 Hz, 1H), 4.24 (p, $J$ = 7.3 Hz, 1H), 3.61 (s, 3H), 2.79 – 2.65 (m, 2H), 2.34† (dd, $J$ = 15.8, 7.9 Hz, 1H), 2.10 (t, $J$ = 7.4 Hz, 2H), 1.59 – 1.40 (m,

4H), 1.40 – 1.05 (m, 11H).[13]C NMR (500 MHz, DMSO-$d_6$). δ = 172.8, 172.1, 171.3, 171.1, 51.9, 49.5, 47.6, 37.4, 35.1, 28.3, 27.8, 25.8, 25.0, 17.1. HRMS (ESI): calcd for $C_{16}H_{31}N_4O_5$ $[M+H]^+$, 359.2294; found, 359.2278. $[\alpha]_D^{21.5}$ = + 18.5° (*c* = 0.57, DMSO)

14 distinct $^{13}$C signals are observed for this substrate (16 are expected), which is likely due to the close overlap of peaks from the acyl chain in the 25-29 ppm range. 2D HSQC data is included for this substrate in the next section.

† see "Note on NMR spectra" at the beginning of this section

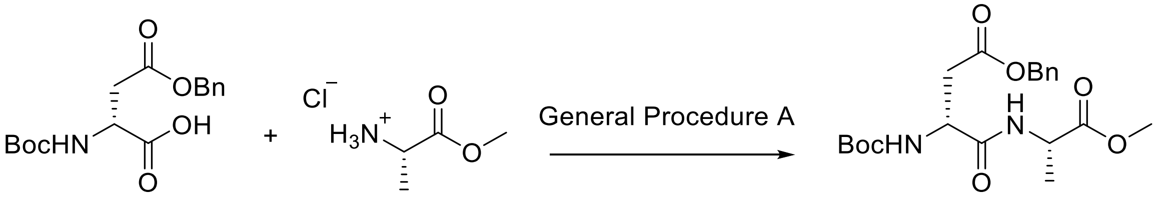

**27**

Dipeptide **27** was assembled according to General Procedure A using *N*-Boc-D-aspartic acid 4-benzyl ester (Oakwood Chemicals, 500 mg) and L-alanine methylester hydrochloride (Chem-Impex International, 238 mg). Crude **27** was isolated as a solid after work-up and purified by flash chromatography (silica, EtOAc:Hexanes 50:50 to 80:20) affording **27** as a clear, viscous oil (240 mg, 38% yield). $^{1}$H NMR (400 MHz, DMSO-$d_6$): δ (ppm) = 8.19 (d, *J* = 7.4 Hz, 1H), 7.45 – 7.25 (m, 5H), 7.06 (d, *J* = 8.5 Hz, 1H), 5.08 (s, 2H), 4.38 (dt, *J* = 8.5, 5.4 Hz, 1H), 4.26 (quint, *J* = 7.2 Hz, 1H), 3.61 (s, 3H), 2.76 (dd, *J* = 16.0, 5.4 Hz, 1H), 2.60 (dd, *J* = 16.1, 8.7 Hz, 1H), 1.38 (s, 9H), 1.24 (d, *J* = 7.2 Hz, 3H). $^{13}$C NMR (500 MHz, DMSO-$d_6$): δ (ppm) = 172.7, 170.4, 170.0, 155.1, 136.0, 128.3, 127.9, 127.8, 78.4, 65.6, 51.8, 50.6, 47.6, 36.4, 28.1, 17.1. HRMS (ESI): calcd for $C_{20}H_{29}N_2O_7$ $[M+H]^+$, 409.1975; found, 409.1993.

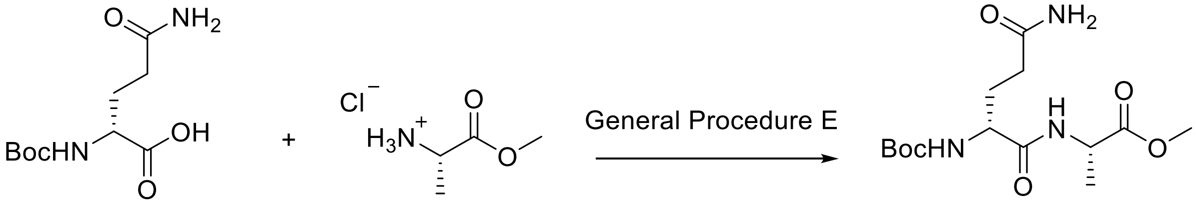

**29**

Dipeptide **29** was assembled according to General Procedure E using *N*-Boc-D-glutamine (185 mg) and L-alanine methylester hydrochloride (105 mg). The coupling was carried out at a concentration of 0.1 M *N*-Boc-D-glutamine and worked up using EtOAc to afford crude **29** as a viscous, colorless oil containing residual tripyrrolidinophosphine oxide, see note below* (127.2 mg, ~20% pure based on NMR, actual yield = 25.5 mg of **29**, 10% yield). $^{1}$H NMR (400 MHz, $CDCl_3$): δ (ppm) = 7.33 (br s, 1H), 6.95 (br s, 1H), 6.02 (br s, 1H), 5.62 (br s, 1H), 4.49 (quint, *J* = 7.2 Hz, 1H), 4.16 (br s, 1H), 3.68 (s, 3H), 3.19 – 3.07* (m, 56H), 2.42 (m, 1H), 2.36 – 2.24 (m, 1H), 2.14 – 1.85 (m, 2H), 1.79* (m, 56H), 1.44 – 1.33 (m, 12H)‡. $^{13}$C NMR (400 MHz, $CDCl_3$): δ (ppm) = 175.9, 173.3, 171.2, 156.0, 77.4, 52.4, 48.3, 48.1, 46.7*, 31.8, 31.4, 28.4, 26.4*, 17.9. HRMS (ESI): calcd for $C_{14}H_{26}N_3O_6$ $[M+H]^+$, 332.1822; found, 332.1825.

*Tripyrrolidinophosphine oxide is a known side product of PyBOP coupling reactions. This material is difficult to remove but did not interfere with the next reaction and was removed in a subsequent purification step.

‡ Predicted overlap of 9H singlet (Boc group) and 3H triplet (alanine $\beta$-methyl group)

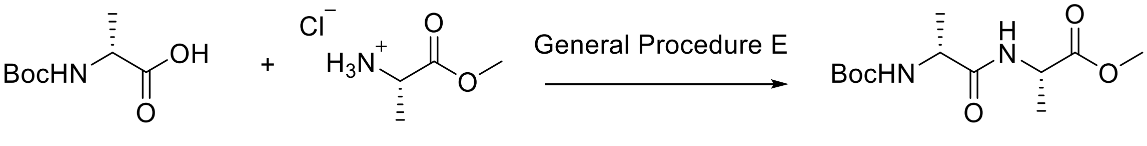

**30**

Dipeptide **30** was assembled according to General Procedure E using *N*-Boc-D-alanine (142 mg) and L-alanine methylester hydrochloride (105 mg). The reaction was worked up in EtOAc to afford **30** as a viscous, colorless oil containing residual tripyrrolidinophosphine oxide, see note below* (283.7 mg, 58% pure based on NMR; actual yield = 163.6 mg of **30**¸ 79% yield). $^1$H NMR (400 MHz, $CDCl_3$): δ (ppm) = 6.94 (br s, 1H), 5.19 (br s, 1H), 4.49 (quint, *J* = 7.4 Hz, 1H), 4.06 (q, *J* = 7.2 Hz, 1H), 3.68 (s, 3H), 3.19 – 3.07* (m, 8H), 1.79* (m, 8H), 1.39 (s, 9H), 1.35 (d, *J* = 7.3 Hz, 3H), 1.30 (d, *J* = 7.1 Hz, 3H). $^{13}$C NMR (400 MHz, $CDCl_3$): δ (ppm) = 173.3, 172.5, 155.6, 80.1, 52.4, 50.0, 48.0, 46.5*, 28.3, 26.4*, 18.3, 18.2. HRMS (ESI): calcd for $C_{12}H_{23}N_2O_5$ [M+H]$^+$, 275.1607; found, 275.1611.

* denotes a peak originating from tripyrrolidinophosphine oxide, a known side product of the PyBOP coupling used here. This contaminant is difficult to remove but does not interfere with the next reaction and was removed in a subsequent step.

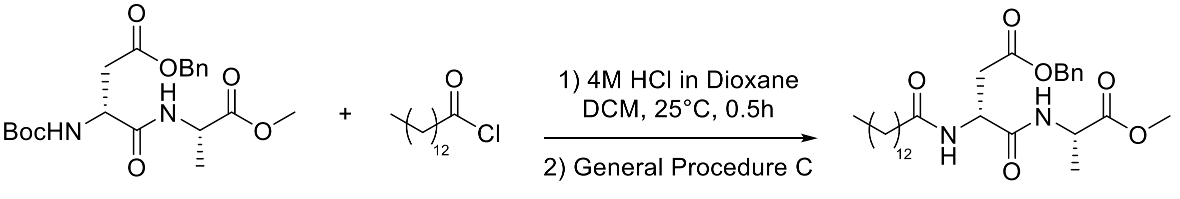

**27** **31**

To a round bottom flask equipped with a stir bar was added **27** (240 mg, 1.0 equiv), 5.1 mL of anhydrous DCM, and 3.7 mL of 4 M HCl in dioxane. The reaction mixture was stirred at room temperature until TLC indicated the Boc deprotection was complete. The reaction mixture was then sparged with $N_2$ for 10 minutes and concentrated *in vacuo* to afford the amine hydrochloride salt as a thick, clear oil. This oil was dissolved in anhydrous DMF (3.9 mL) and coupled to myristoyl chloride following General Procedure C. After stirring at room temperature overnight, **31** was precipitated out of the reaction mixture by the addition of 5% aqueous LiCl (10 mL). The precipitate was filtered and washed with saturated aqueous $NaHCO_3$, 0.1 M HCl, water, and cold diethyl ether ($Et_2O$). Some residual myristic acid produced by hydrolysis of the acyl chloride remained and removal was attempted using flash chromatography (C18 functionalized silica, CombiFlash RediSepRf system, Teledyne ISCO, Lincoln NE) eluting in water/acetonitrile. This method was unable to fully remove the residual myristic acid despite trying a variety of linear gradient conditions and the product was used in the next step without further purification (227 mg, 64 % pure based on NMR; actual yield = 145.3 mg of **31**, 48% yield). $^1$H NMR (600 MHz DMSO-$d_6$): δ (ppm) = 8.27 (d, *J* = 7.2 Hz, 1H), 8.11 (d, *J* = 8.5 Hz, 1H), 7.47 – 7.29 (m, 5H), 5.06 (s, 2H),

4.71 (td, $J$ = 8.3, 5.8 Hz, 1H), 4.24 (p, $J$ = 7.3 Hz, 1H), 3.60 (s, 3H), 2.77 (dd, $J$ = 15.8, 5.8 Hz, 1H), 2.58 (dd, $J$ = 15.9, 8.4 Hz, 1H), 2.18* (t, $J$ = 7.4 Hz, 2H), 2.07 (t, $J$ = 7.4 Hz, 2H), 1.54 – 1.40* (m, 5H), 1.29 – 1.16* (m, 45H), 0.89 – 0.81* (m, 6H). $^{13}$C NMR (500 MHz, DMSO-$d_6$) δ (ppm) = 173.1, 172.7, 170.6, 170.3, 136.5, 128.8, 128.4, 128.2, 66.1, 52.3, 49.4, 48.1, 36.9, 35.6, 31.8, 29.5, 29.5, 29.4, 29.3, 29.2, 29.1, 29.0, 28.9, 25.6, 22.6, 17.5, 14.4. HRMS (ESI): calcd for $C_{29}H_{47}N_2O_6$ [M+H]$^+$, 519.3434; found, 519.3446.

*peaks originating from or overlapping with peaks from myristic acid

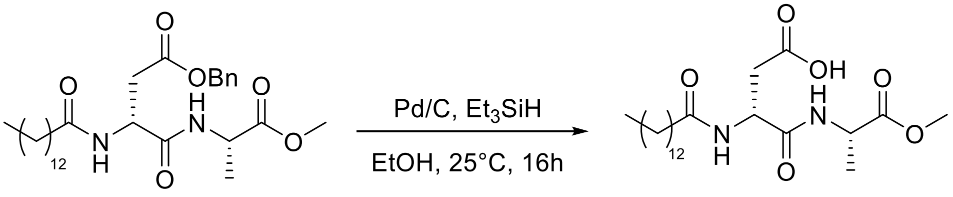

**31** **6**

To a round bottom flask equipped with a stir bar was added palladium on carbon (15 mg, 10% by weight) under $N_2$ and water (0.5 mL). Acyl dipeptide **31** (80 mg, 1.0 equiv) was suspended in ethanol (EtOH, 10 mL) in a separate vial and sonicated briefly. The slurry of **31** was added to the reaction flask and residual slurry was washed out of the vial using an additional 2 mL of ethanol and added to the reaction flask. The reaction mixture was flushed with $N_2$ while stirring for 10 minutes. Triethylsilane (246 μL, 10.0 equiv) was added dropwise, and gas evolution was observed. The reaction mixture was stirred at room temperature overnight, filtered over Celite to remove the catalyst, and the filtrate was concentrated *in vacuo*. Isopropanol (10 mL) was added and the reaction mixture concentrated *in vacuo* again. The process was repeated 2 additional times. At this point, the product was a viscous oil with some solids present. The residue was triturated with cold hexanes and filtered to isolate **6** as a white solid (71 mg, quantitative yield). This material was then dissolved in a minimum volume of DMSO and purified by preparative HPLC according to the procedure described in the General Materials and Methods to afford **6** as an off-white solid (58 mg, 82% yield). $^1$H NMR (600 MHz DMSO-$d_6$): δ (ppm) = 12.25 (br s, 1H), 8.15 (d, $J$ = 7.2 Hz, 1H), 8.03 (d, $J$ = 8.2 Hz, 1H), 4.60 (dt, $J$ = 8.1, 6.1 Hz, 1H), 4.25 (quint, $J$ = 7.3 Hz, 1H), 3.61 (s, 3H), 2.62 (dd, $J$ = 16.7, 6.1 Hz, 1H), 2.43 (dd, $J$ = 16.3, 7.9 Hz, 1H), 2.09 (t, $J$ = 7.4 Hz, 2H), 1.52 – 1.41 (m, 2H), 1.31 – 1.17 (m, 23H), 0.85 (t, $J$ = 7.0 Hz, 3H). $^{13}$C NMR (400 MHz, DMSO-$d_6$) δ (ppm) = 172.7, 172.3, 171.2*, 170.6, 51.8, 49.1, 47.6, 36.4, 35.2, 31.3, 29.05, 29.03, 29.01, 28.9, 28.8, 28.7, 28.6, 25.2, 22.1, 17.1, 13.9. HRMS (ESI): calcd for $C_{22}H_{41}N_2O_6$ [M+H]$^+$, 429.2965; found, 429.2958. $[\alpha]_D^{22}$ = + 13.2° ($c$ = 0.63, DMSO)

*low signal intensity in 1D $^{13}$C-NMR, but visible in the HMBC spectrum of this compound (see spectra below)

21 distinct $^{13}$C signals are observed for this substrate (22 are expected), which is likely due to the close overlap of peaks from the myristoyl group in the 28.5-29 ppm range.

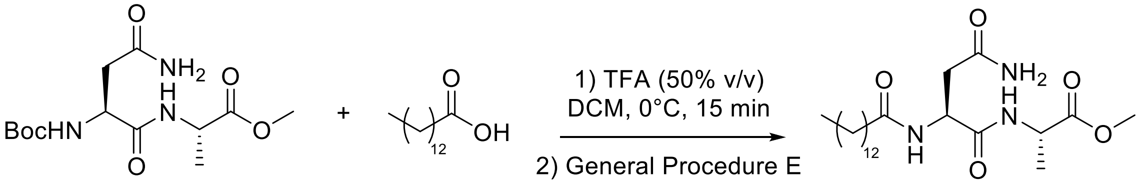

**28** **7**

To an oven-dried round bottom flask was added intermediate **28** (prepared as previously described,[10] 163 mg, 1.0 equiv) and DCM (2.5 mL). The mixture was cooled to 0 °C and trifluoroacetic acid (2.5 mL) was added dropwise. The reaction was stirred at 0 °C until TLC indicated the Boc deprotection reaction was complete (approximately 15 minutes). Anhydrous toluene (5 mL) was added and the reaction mixture concentrated *in vacuo*. An additional 5 mL of toluene was added, and the reaction mixture concentrated again to remove residual TFA. To this flask was added DMF, myristic acid, HOBt, and PyBOP according to General Procedure E. The coupling was carried out at a concentration of 0.1 M **28** and worked up using EtOAc. The crude product was isolated as a viscous, colorless oil which was subsequently purified by preparative HPLC according to the procedure described in the General Materials and Methods to afford **7** as an off-white solid (40 mg, 18% yield over two steps). $^{1}$H NMR (600 MHz DMSO-$d_6$): δ (ppm) = 8.15 (d, $J$ = 7.2 Hz, 1H), 7.93 (d, $J$ = 8.0 Hz, 1H), 7.25 (s, 1H), 6.85 (s, 1H), 4.55 (dt, $J$ = 8.3, 4.9 Hz, 1H), 4.22 (quint, $J$ = 7.3 Hz, 1H), 3.59 (s, 3H), 2.46 (dd, $J$ = 15.5, 5.1 Hz, 1H), 2.34 (dd, $J$ = 15.4, 8.7 Hz, 1H), 2.07 (t, $J$ = 7.5 Hz, 2H), 1.45 (m, 2H), 1.30 – 1.15 (m, 23H), 0.88 – 0.79 (m, 3H).$^{13}$CNMR (400 MHz, DMSO-$d_6$) δ (ppm) = 172.9, 172.2, 171.4* 171.3, 51.8, 49.4, 47.6, 37.3, 35.2, 31.3, 29.05, 29.03, 29.00, 28.9, 28.8, 28.7, 28.6, 25.2, 22.1, 16.9, 14.0. HRMS (ESI): calcd for $C_{22}H_{42}N_3O_5$ [M+H]$^+$, 428.3124; found, 428.3112. $[\alpha]_D^{22}$ = –8.1° ($c$ = 0.5, DMSO)

*low signal intensity in 1D $^{13}$C-NMR, but visible in the HMBC spectrum of this compound (see spectra below)
21 distinct $^{13}$C signals are observed for this substrate (22 are expected), which is likely due to the close overlap of peaks from the myristoyl group in the 28.5-29 ppm range.

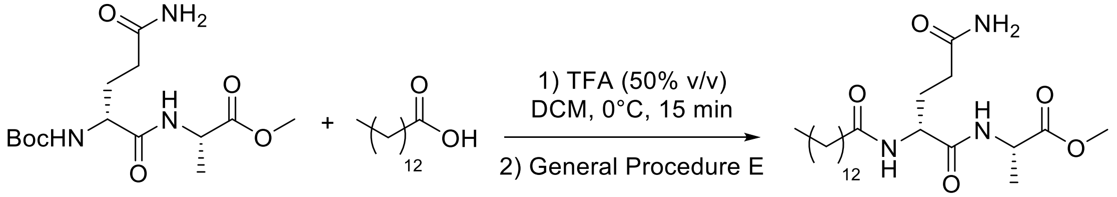

**29** **8**

Compound **8** was prepared from compound **29** (270 mg) following the same procedure as described for the preparation of compound **7**. Crude **8** was obtained as a clear viscous oil (150 mg, 42% crude yield) which was dissolved in DMSO and purified by preparative HPLC as described in the General Materials and Methods to afford pure **8** as a white solid (71 mg, 20% yield over two steps). $^{1}$H NMR (500 MHz DMSO-$d_6$): δ (ppm) = 8.27 (d, $J$ = 7.3 Hz, 1H), 7.88 (d, $J$ = 8.3 Hz, 1H), 7.25 (s, 1H), 6.74 (s, 1H), 4.27 (m, 2H), 3.62 (s, 3H), 2.11 (t, $J$ = 7.6 Hz, 2H), 2.05 (t, $J$ = 8.4 Hz, 2H), 1.88 – 1.61 (m, 2H), 1.53 – 1.42 (m, 2H), 1.26 (m, 23H), 0.86 (t, $J$ = 6.7 Hz, 3H). $^{13}$C NMR (500 MHz, DMSO-d$_6$): δ (ppm) = 173.6, 172.8, 172.1, 171.3, 51.8, 47.4, 35.2, 33.7, 31.4, 31.3, 29.04, 29.01, 28.96, 28.90, 28.8, 28.71, 28.7, 28.5, 28.2, 25.2, 22.1, 17.1, 13.9. HRMS (ESI): calcd for $C_{23}H_{44}N_3O_5$ [M+H]$^+$, 442.3281; found, 442.3284. $[\alpha]_D^{22}$ = + 2.7 ° ($c$ = 0.5, DMSO)

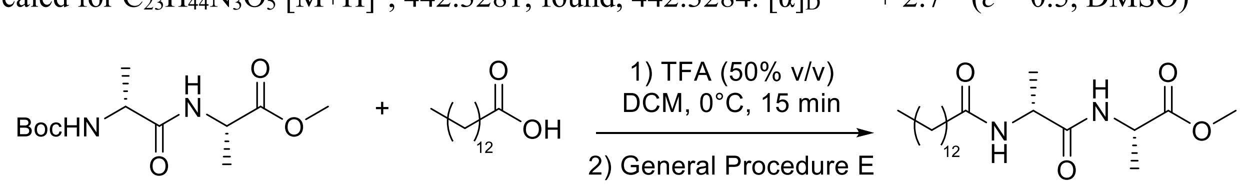

**30** **9**

Compound **9** was prepared from compound **30** (720 mg) following the same procedure as described for the preparation of compound **7**. Compound **9** was purified by preparative HPLC as described in the General Materials and Methods, but could not be fully resolved from residual myristic acid (86% pure based on $^{1}$H-NMR. 30% yield as a white solid). $^{1}$H NMR (400 MHz DMSO-$d_6$): δ (ppm) = 8.22 (d, *J* = 7.2 Hz, 1H), 7.90 (d, *J* = 7.8 Hz, 1H), 4.32 (quint, *J* = 7.4 Hz, 1H), 4.25 (quint, *J* = 7.2 Hz, 1H), 3.61 (s, 3H), 1.50 – 1.41 (m, 2H), 1.34 – 1.13 (m, 28H), 0.85 (t, J = 6.8 Hz, 3H). $^{13}$C NMR (400 MHz, DMSO-$d_6$) δ (ppm) = 173.0*, 172.9, 172.4*, 172.2, 171.83*, 171.82, 51.8, 47.6*, 47.4, 45.9*, 45.8, 35.1, 31.3, 29.1, 29.05, 29.01, 28.9, 28.8, 28.7, 28.6, 25.94*, 25.86, 25.2, 22.1, 18.5, 18.2*, 17.1, 16.8*, 13.9. HRMS (ESI): calcd for $C_{21}H_{41}N_2O_4$ [M+H]$^+$, 385.3066; found, 385.3058. $[\alpha]_D^{22}$ = – 2.8 ° (*c* = 1.0, DMSO)

* denotes a rotamer

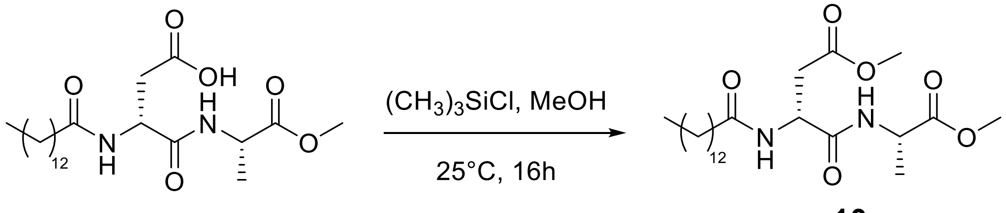

**6** **10**

To an oven-dried round bottom flask was added compound **6** (30 mg, 1.0 equiv) and trimethylsilyl chloride (20 µL, 2 equiv) was added under $N_2$. The suspension was then dissolved in anhydrous methanol (0.5 mL, 42 equiv) and stirred at room temperature overnight. The reaction was concentrated *in vacuo* and the resulting solid was purified by preparative HPLC according to the procedure described in the General Materials and Methods to afford **10** as a white solid (13 mg, 43% yield after prep HPLC). $^{1}$H NMR (600 MHz DMSO-$d_6$): δ (ppm) = 8.26 (d, *J* = 7.3 Hz, 1H), 8.12 (d, *J* = 8.3 Hz, 1H), 4.66 (td, *J* = 8.2, 6.1 Hz, 1H), 4.25 (p, *J* = 7.2 Hz, 1H), 3.61 (s, 3H), 3.56 (s, 3H), 2.70† (dd, *J* = 15.8, 6.1 Hz, 1H), 2.09 (t, *J* = 7.4 Hz, 2H), 1.55 – 1.39 (m, 2H), 1.33 – 1.11 (m, 23H), 0.85 (t, *J* = 6.8 Hz, 3H). $^{13}$C NMR (500 MHz, DMSO-$d_6$) δ (ppm) = 172.6, 172.3, 170.4, 170.2, 51.8, 51.4, 48.9, 47.6, 36.2, 35.1, 31.3, 29.04, 29.00, 28.9, 28.8, 28.7, 28.5, 25.2, 22.1, 17.0, 13.9. HRMS (ESI): calcd for $C_{23}H_{43}N_2O_6$ [M+H]$^+$, 443.3121; found, 443.3106. $[\alpha]_D^{22}$ = + 14.4 ° (*c* = 0.7, DMSO)

† see "Note on NMR spectra" at the beginning of this section

21 distinct $^{13}$C signals are observed for this substrate (23 are expected), which is likely due to the close overlap of peaks from the myristoyl group in the 28.5-29 ppm range.

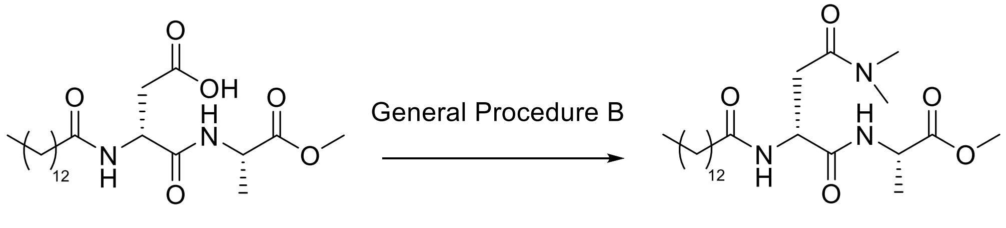

**6** **11**

Compound **11** was prepared from compound **6** (30 mg, 1.0 equiv) and dimethylamine (as a 2 M solution in THF, 52.5 µL, 1.5 equiv) according to General Procedure B. Crude **11** was isolated as an off-white solid after work-up (38 mg, quantitative crude yield). This solid was dissolved in

DMSO and purified by preparative HPLC according to the procedure described in the General Materials and Methods to afford pure **11** as a white solid (5.7 mg, 18% final yield). $^{1}$H NMR (500 MHz DMSO-d$_6$): δ (ppm) = 8.11 (d, *J* = 7.2 Hz, 1H), 7.99 (d, *J* = 8.1 Hz, 1H), 4.62 (dt, *J* = 8.2, 6.7 Hz, 1H), 4.22 (quint, *J* = 7.2 Hz, 1H), 3.58 (s, 3H), 2.93 (s, 3H), 2.77 (s, 3H), 2.64 (dd, *J* = 15.9, 6.5 Hz, 1H), 2.56 (dd, *J* = 15.9, 6.8 Hz, 1H), 2.06 (t, *J* = 7.4 Hz, 2H), 1.44 (m, 2H), 1.26 – 1.18 (m, 23H), 0.83 (t, *J* = 6.9 Hz, 3H). $^{13}$C NMR (500 MHz, DMSO-*d*$_6$) δ (ppm) = δ 172.8, 172.1, 171.0, 169.1, 51.8, 49.3, 47.6, 36.70, 36.68, 35.2, 34.8, 31.3, 29.03, 29.0, 28.9, 28.8, 28.7, 28.5, 25.2, 22.1, 17.1, 13.9. HRMS (ESI): calcd for $C_{24}H_{46}N_3O_5$ $[M+H]^+$, 456.3437; found, 456.3423. $[\alpha]_D^{22}$ = + 18.2 ° (*c* = 0.48, DMSO)

21 distinct $^{13}$C signals are observed for this substrate (23 are expected), which is likely due to the close overlap of peaks from the myristoyl group in the 28.5-29 ppm range.

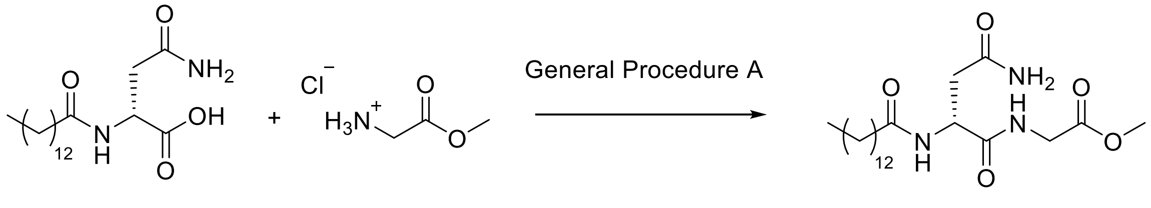

**19** **12**

Substrate **12** was prepared from **19** (14.2 mg) and glycine methylester hydrochloride (6.3 mg) using General Procedure A. Crude **12** was isolated as a white solid after work-up (18 mg, quantitative). This solid was dissolved in DMSO and purified by preparative HPLC according to the procedure described in the General Materials and Methods to afford pure **12** as a white solid (12 mg, 70%). $^{1}$H NMR (400 MHz DMSO-*d*$_6$): δ (ppm) = 8.11 (t, *J* = 5.9 Hz, 1H), 7.99 (d, *J* = 8.1 Hz, 1H), 7.27 (s, 1H), 6.84 (s, 1H), 4.59 (td, *J* = 8.1, 5.4 Hz, 1H), 3.89 – 3.73 (m, 2H), 3.61 (s, 3H), 2.35† (dd, *J* = 15.4, 8.3 Hz, 1H), 2.09 (t, *J* = 7.5 Hz, 2H), 1.46 (t, *J* = 7.1 Hz, 2H), 1.23 (d, *J* = 2.4 Hz, 20H), 0.90 – 0.81 (m, 3H).$^{13}$C NMR (400 MHz, DMSO-*d*$_6$) δ (ppm) = 172.2, 171.8, 171.3, 170.1, 51.6, 49.3, 40.7, 37.2, 35.2, 31.3, 29.1, 29.05, 29.01, 28.93, 28.9, 28.7, 28.6, 25.1, 22.1, 13.9. HRMS (ESI): calcd for $C_{21}H_{40}N_3O_5$ $[M+H]^+$, 414.2968; found, 414.2951. $[\alpha]_D^{22}$ = + 8.5° (*c* = 0.98, DMSO)

† see "Note on NMR spectra" at the beginning of this section

20 distinct $^{13}$C signals are observed for this substrate (21 are expected), which is likely due to the close overlap of peaks from the myristoyl group in the 28.5-29 ppm range.

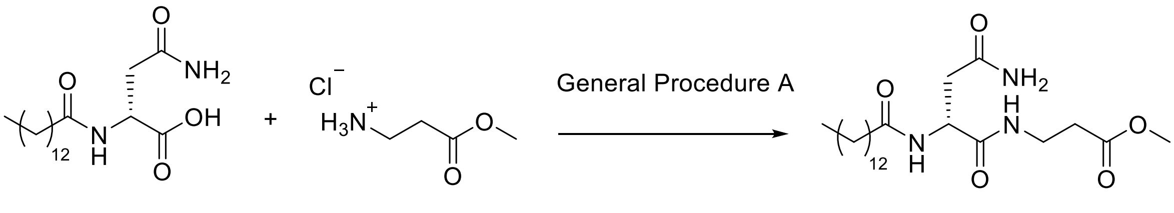

**19** **13**

Substrate **13** was prepared from **19** (13.5 mg) and β-alanine methylester hydrochloride (6.6 mg) using General Procedure A. Crude **13** was isolated as a white solid after work up (18.9 mg, quantitative). This solid was dissolved in DMSO and purified by preparative HPLC according to the procedure described in the General Materials and Methods to afford **13** as a white solid (14 mg, 59%). $^1$H NMR (400 MHz DMSO-$d_6$): δ (ppm) = 7.88 (d, $J$ = 8.0 Hz, 1H), 7.74 (t, $J$ = 5.8 Hz, 1H), 7.21 (s, 1H), 6.78 (s, 1H), 4.43 (td, $J$ = 7.8, 5.9 Hz, 1H), 3.55 (s, 3H), 3.27 – 3.16 (m, 2H), 2.41 (dd, $J$ = 15.3, 6.0 Hz, 1H), 2.39 (t, $J$ = 6.9 Hz, 2H), 2.27 (dd, $J$ = 15.3, 7.6 Hz, 1H), 2.04 (t, $J$ = 7.5 Hz, 2H), 1.42 (q, $J$ = 7.1 Hz, 2H), 1.20 (s, 20H), 0.87 – 0.74 (m, 3H). $^{13}$C NMR (400 MHz, DMSO-$d_6$) δ (ppm) = 172.1, 171.7, 171.4, 171.2, 51.3, 49.6, 37.3, 35.2, 34.7, 33.5, 31.3, 29.1, 29.04, 29.01, 29.0, 28.9, 28.8, 28.7, 28.6, 25.1, 22.1, 14.0. HRMS (ESI): calcd for $C_{22}H_{42}N_3O_5$ $[M+H]^+$, 428.3124; found, 428.3107. $[\alpha]_D^{22}$ = + 5.5° ($c$ = 0.43, DMSO)

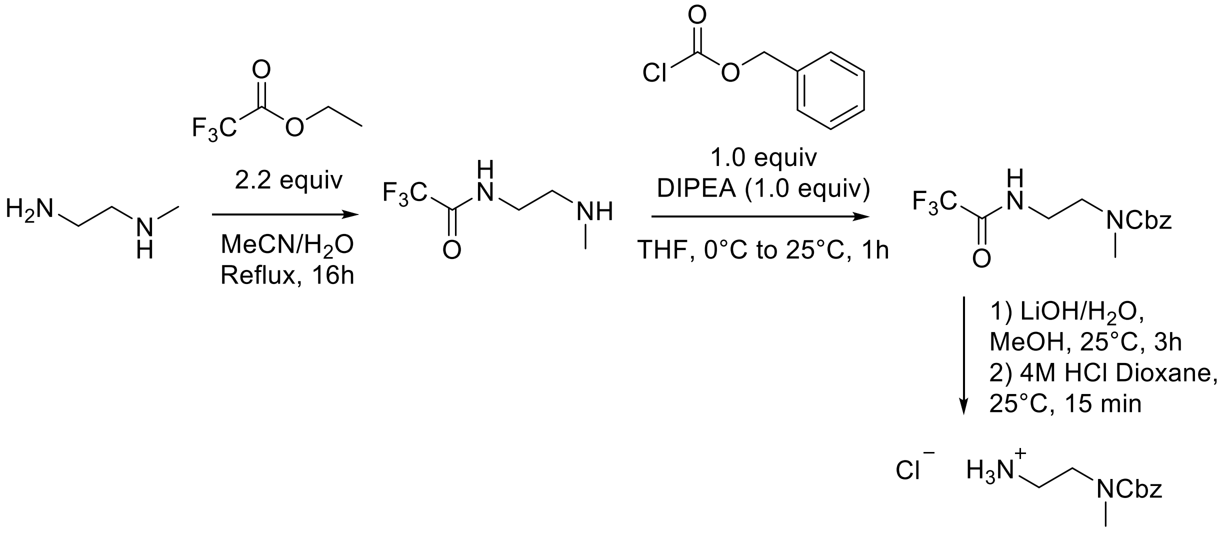

**32**

Intermediate **32** was synthesized has previously reported.[13] $N$-Methylethylenediamine (2.6 g, 1.0 equiv), ethyl trifluoroacetate (3.5 mL, 2.2 equiv) and water (0.76 mL, 1.2 equiv) were combined in acetonitrile (35 mL) and refluxed with stirring overnight. Solvents were evaporated under vacuum and the residue was re-evaporated with i-PrOH three times. The trifluoroacetamide product was recrystallized from DCM and filtered. This product (4.0 g, 1.0 equiv) was then dissolved in THF (26 mL) with DIPEA (4.09 mL, 1.0 equiv) and cooled to 0 °C, at which point benzyloxycarbonyl chloride (3.35 mL, 1.0 equiv) was added dropwise. The reaction was warmed to room temperature and stirred for an additional 1 hour. Solvent was then evaporated in vacuo and the resulting residue dissolve in EtOAc. The organic layer was washed with 5% aqueous $NaHCO_3$ twice and brine. The organic layer was evaporated to provide the benzyloxy-protected trifluoroacetamide product as a yellowish oil. This oil was then dissolved in MeOH (78 mL) and a solution of LiOH (1.12 g, 2.0 equiv) in water (9.5 mL) was added and the mixture stirred for 3 hours at room temperature. Solvents were evaporated to 75% of the initial volume and then diluted with water (30 mL). The solution was extracted twice with EtOAc. The combined organic layers were then washed with brine (50 mL), dried over $Mg_2SO_4$, filtered, and the filtrate concentrated *in vacuo*. The resulting residue was then dissolved in $Et_2O$ (25 mL) and treated with 4N HCl in dioxane (7.7 mL). After 15 minutes, the solvent was removed *in vacuo* to afford HCl salt **32**.

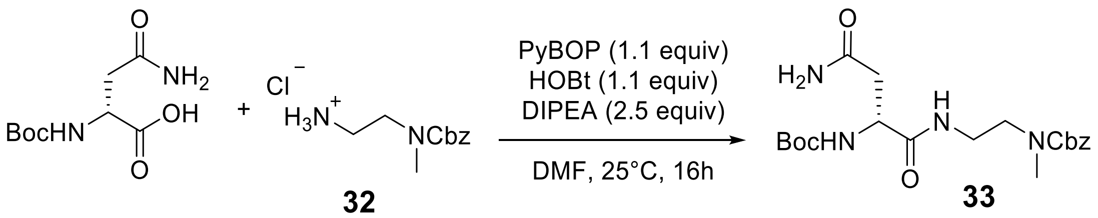

To a flame-dried round bottom flask containing a stir bar was added amine hydrochloride salt **32** (1.65 g, 1.0 equiv), *N*-Boc-D-Asparagine (1.57 g, 1.0 equiv), PyBOP (3.87 g, 1.1 equiv), HOBt (1.00 g, 1.1 equiv), and dry DMF (13.5 mL, 0.5 M). The reaction mixture was cooled to 0 °C and DIPEA (2.93 mL, 2.5 equiv) was added dropwise under $N_2$. The resulting reaction mixture was then warmed to room temperature over 12 h. The reaction was then quenched with aqueous 1M HCl (10 mL) and diluted with EtOAc (20 mL). The layers were separated and the aqueous layer extracted 3 times with EtOAc. The combined organic layers were then washed with satatuarated aqueous $NaHCO_3$, twice with water, and brine. The organic layer was dried over $Na_2SO_4$, filtered, and the resulting filtrate concentrated *in vacuo* to give a crude solid. The solid was triturated with $Et_2O$ (20 mL) and dried under vacuum to afford carbamate **33** as a light brown solid (2.08 g, 73%). $^1$H NMR (600 MHz DMSO-*d*$_6$): δ (ppm) = 7.92 (br s, 1H); 7.39-7.29 (m, 5H); 7.24 (br s, 1H); 6.87 (br s, 1H); 6.83 – 6.77 (m, 1H); 5.06 (s, 2H); 4.23 – 4.14 (m, 1H); 3.30 – 3.24 (m, 2H); 3.24 – 3.14 (m, 2H); 2.87 (s, 1.5H)*; 2.84 (s, 1.5H)*; 2.42 – 2.31 (m, 2H); 1.37 (s, 9H). 13C NMR (500 MHz, DMSO-*d*$_6$): δ (ppm) = 171.6, 171.5, 155.5, 155.3*, 155.0, 137.1, 137.0*, 128.4, 127.72, 127.69*, 127.40, 127.37*, 78.1, 66.1, 51.4, 48.0, 47.6*, 37.3, 37.0, 36.99*, 35.2, 34.5*, 28.2. HRMS (ESI): calcd for $C_{20}H_{31}N_4O_6$ $[M+H]^+$, 423.2238; found, 423.2247. $[\alpha]_D^{23}$ = + 4.8° (*c* = 1.0, DMSO)

* = denotes a rotamer

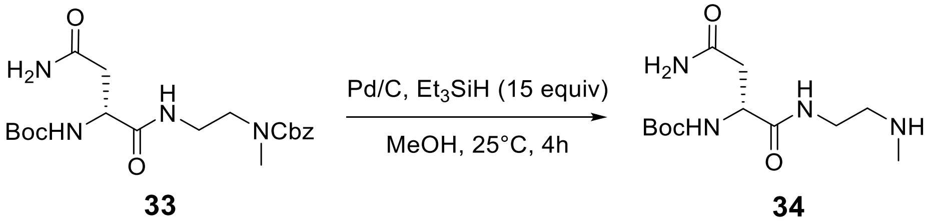

To a round bottom flask equipped with a stir bar was added palladium on carbon (22 mg, 10% Pd by weight) and water (0.5 mL) under $N_2$. Separately, carbamate **33** (318 mg, 1.0 equiv) was suspended in MeOH (6 mL) in a vial. The mixture was sonicated in order to uniformly suspend the solid and the resulting slurry was transferred to the reaction flask via syringe. The vial was rinsed with additional MeOH (2 mL) which was added to the reaction vessel. Triethylsilane (1.1 mL, 15.0 equiv) was added dropwise while the reaction mixture was stirred at room temperature and gas evolution was observed. The reaction was stirred for 4 hours, during which time the mixture went from a gray slurry to a clear solution with only the black catalyst visible. The reaction was then filtered over Celite to remove the catalyst and the filtrate concentrated in vacuo. The resulting residue was then dissolved in isopropanol and concentrated *in vacuo* twice to afford crude amine **34**, which was used in the subsequent step without further purification.

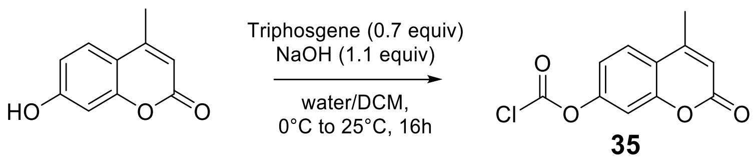

Intermediate **35** was prepared as previously described by Hesserodt and coworkers.[14] To a round bottom flask under $N_2$ was added triphosgene (1.26 g, 0.7 equiv) under $N_2$. Dry DCM (30mL) was added and the solution cooled to 0 °C. 7-Hydroxy-4-methylcoumarin (1.08 g, 1.0 equiv) was then added in one portion. To the resulting white suspension was added 2M aqueous NaOH (3.3 mL) dropwise. The reaction was stirred at 0 °C for 1 hour, then warmed to room temperature and stirred overnight. The precipitated white solid was filtered and washed with cold DCM affording **35** as a white solid (0.570 g, 40%). $^1$H NMR (500 MHz, DMSO-$d_6$) δ 7.91 (d, $J$ = 8.7 Hz, 1H), 7.59 (d, $J$ = 2.3 Hz, 1H), 7.48 (dd, $J$ = 8.7, 2.4 Hz, 1H), 6.44 (d, $J$ = 1.5 Hz, 1H), 2.46 (s, 3H).

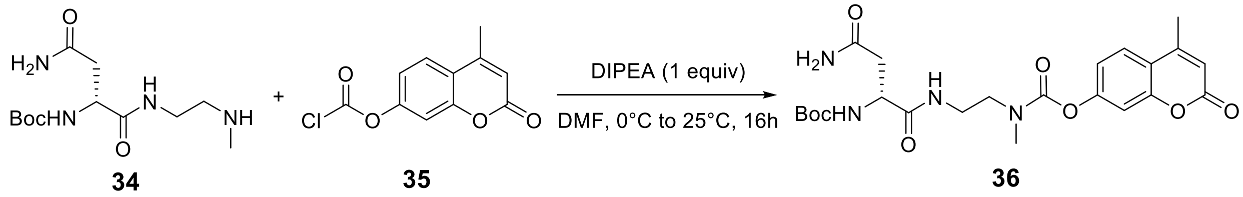

To a flame-dried round bottom flask equipped with a stir bar was added **34** (181 mg, 1.0 equiv) and **35** (165 mg, 1.1 equiv) under $N_2$. Dry DMF (6.3 mL) was added and the flask cooled to 0 °C. DIPEA (121 µL, 1.1 equiv) was added dropwise and the reaction stirred overnight at room temperature. The reaction was diluted with water (30 mL) and extracted twice with DCM (30 mL). The combined organic layers were washed with 5% aqueous LiCl, 1M HCl, 0.1M NaOH, and brine. The organic layer was then concentrated *in vacuo* and purified by flash chromatography (silica, EtOAc:MeOH 90:10, isocratic). Fractions containing the desired product were concentrated *in vacuo* to afford **36** as a white solid (175 mg, 57%). $^1$H NMR (500 MHz DMSO-$d_6$): δ (ppm) = 8.12 – 7.95 (m, 1H), 7.78 (d, $J$ = 8.6 Hz, 1H), 7.32 – 7.23 (m, 2H), 7.20 (ddd, $J$ = 8.8, 6.5, 2.3 Hz, 1H), 6.88 (t, $J$ = 6.7 Hz, 2H), 6.37 (s, 1H), 4.21 (m, 1H), 3.51 – 3.21 (m, 4H), 3.05 (s, 1.5H)*, 2.92 (s, 1.5H)*, 2.44 (s, 3H), 2.42 – 2.29 (m, 2H), 1.37 (s, 9H). $^{13}$C NMR (500 MHz, DMSO-$d_6$) δ 171.8, 171.6, 159.7, 155.1, 153.9, 153.5, 153.3, 153.0, 126.0, 118.4, 116.9, 113.4, 109.9, 109.8*, 78.2, 51.5, 48.5, 48.46*, 37.40*, 37.3, 36.6, 35.7, 35.3*, 28.2, 18.2. HRMS (ESI): calcd for $C_{23}H_{31}N_4O_8$ $[M+H]^+$, 491.2142; found, 491.2124.

* denotes a rotamer

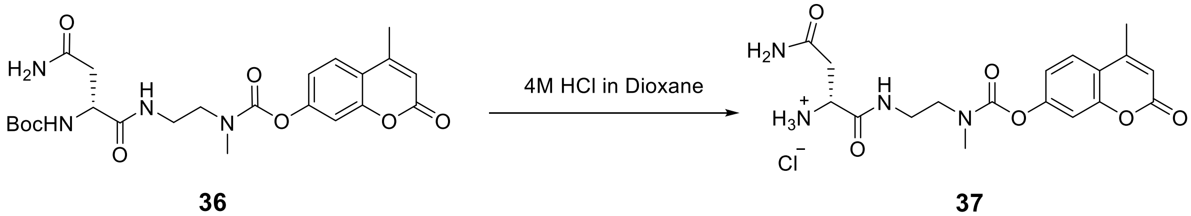

To a round bottom flask with a stir bar was added **36** (190 mg, 1.0 equiv). The starting material was dissolved in DCM (4 mL) and 4M HCl in dioxane (1.94 mL) was added. After stirring for approximately 10 minutes at room temperature, a white precipitate appeared. After 30 minutes, the reaction was sparged with $N_2$ for 10 minutes while stirring to drive off residual HCl. The reaction was then concentrated in vacuo, affording the amine HCl salt **37** as a white solid, which was used without further purification.

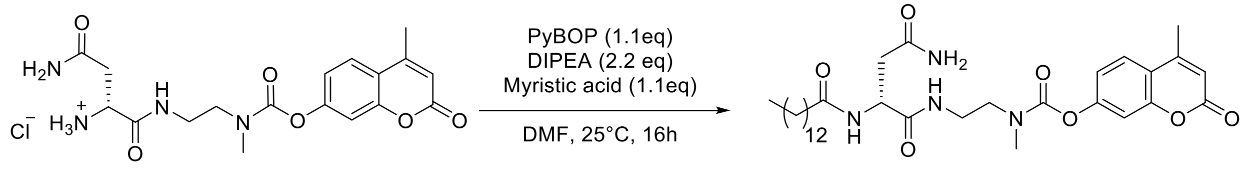

**37** **15**

To an oven dried round-bottom flask were added crude **37** (97 mg, 1.0 equiv), myristic acid (52 mg, 1.2 equiv), PyBOP (143 mg, 1.2 equiv), HOBt (42 mg, 1.2 eq) and anhydrous DMF (2.3 mL) under $N_2$. DIPEA (84 µL, 2.1 equiv) was added dropwise and the reaction allowed to stir overnight at room temperature. The reaction was diluted with water, extracted twice with DCM (5-10 mL), and the combined organic layers were washed with 5% LiCl, 1M HCl, sat. aq. $NaHCO_3$, and brine. The organic layer was dried over $Na_2SO_4$, filtered, and the filtrate concentrated *in vacuo*. The resulting solid was dissolved in a minimum volume of DMSO and purified by preparative HPLC as described in the General Materials and Methods to afford pure **15** as a white solid (18 mg, 13% yield over three steps). $^1$H NMR (400 MHz, DMSO-$d_6$): δ (ppm) = 8.06 – 7.88 (m, 2H), 7.78 (d, $J$ = 8.7 Hz, 1H), 7.33 – 7.24 (m, 2H), 7.20 (dt, $J$ = 8.7, 2.6 Hz, 1H), 6.83 (s, 1H), 6.37 (d, $J$ = 1.4 Hz, 1H), 4.59 – 4.45 (m, 1H), 3.48-3.39 (m, 1H), 3.39 – 3.22 (m, 10H)§, 3.04* (s, 1.5H), 2.93* (s, 1.5H), 2.44 (s, 3H), 2.35† (dd, $J$ = 15.2, 7.7 Hz, 1H), 2.13 – 2.00 (td, $J$ = 7.5, 5.0 Hz, 2H), 1.44 (m, 2H), 1.31-1.10 (m, 20H), 0.85 (t, $J$ = 6.6 Hz, 3H). $^{13}$C NMR (400 MHz, DMSO-$d_6$): δ (ppm) = 172.1*, 172.0, 171.49*, 171.47, 171.42*, 171.4, 166.0, 159.7, 153.9, 153.5, 153.4, 153.03*, 153.0, 126.0, 118.43*, 118.42, 116.86*, 116.85, 113.4, 109.92*, 109.87, 49.8*, 49.7, 48.5*, 48.4, 37.3*, 37.2, 36.7, 35.6, 35.3*, 35.2, 31.3, 29.1, 29.04, 29.01, 28.9, 28.8, 28.70, 28.66, 25.10*, 25.07, 22.1, 18.2, 13.9. HRMS (ESI): calcd for $C_{32}H_{49}N_4O_7$ [M+H]$^+$, 601.3601; found, 601.3612. $[\alpha]_D^{22}$ = + 35.1° ($c$ = 1.0, DMSO)

§ the high integration value of this peak results from overlap with residual water. HSQC data (next section) shows the expected C and H signals for the ethylenediamine linker in this region
* denotes a rotamer
† see "Note on NMR spectra" at the beginning of this section

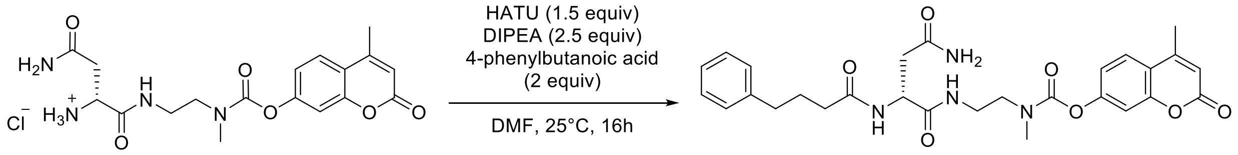

**37** **16**

To an oven-dried round bottom flask equipped with a stir bar were added **37** (44.9 mg, 1.0 equiv), 4-phenylbutanoic acid (38.4 mg, 2.0 equiv), HATU (65.5 mg, 1.5 equiv) and anhydrous DMF (1.14 mL) under $N_2$. DIPEA (50 µL, 2.5 equiv) was added and the reaction allowed to stir at room temperature overnight. The reaction was diluted with water, extracted twice with DCM (5 reaction volumes). The combined organic layers were washed with 5% LiCl, 1M HCl, sat. aq. $NaHCO_3$, and brine. The organic layer was dried over $Na_2SO_4$, filtered, and the filtrate concentrated *in vacuo* to give a white solid. This solid was then dissolved in a minimum volume of DMSO and purified by preparative HPLC as described in the General Materials and Methods to afford **16** as a white solid (38.9 mg, 63% yield over two steps). $^1$H NMR (600 MHz, DMSO-$d_6$): δ (ppm) = δ 8.12 – 7.93 (m, 2H), 7.77 (d, $J$ = 8.6 Hz, 1H), 7.36 – 7.10 (m, 7H), 6.87 (s, 1H), 6.36 (s, 1H), 4.60 – 4.47

(m, 1H), 3.46 – 3.23 (m, 6H) §, 3.04 (s, 1.5H)*, 2.91 (s, 1.5H)*, 2.56 – 2.48 (m, 4H) §, 2.47 (dd, *J* = 9.6, 5.5 Hz, 1H), 2.43 (s, 3H), 2.37† (dd, *J* = 15.2, 8.2 Hz, 1H), 2.17 – 2.06 (m, 2H), 1.82 – 1.68 (m, 2H). $^{13}$C NMR (500 MHz, DMSO-$d_6$): δ (ppm) =172.27, 172.25*, 172.0*, 171.95, 171.92*, 171.9, 160.2, 154.3, 153.9, 153.82*, 153.8, 153.5*, 153.48, 142.3, 128.78, 128.75*, 128.7, 126.5, 126.1, 118.9, 117.3, 113.8, 110.4, 110.3*, 50.33, 50.28*, 48.94*, 48.9, 37.8, 37.75*, 37.7, 37.1, 36.1*, 35.8, 35.2*, 35.1, 27.42, 27.4*, 18.6. HRMS (ESI): calcd for $C_{28}H_{33}N_4O_7$ $[M+H]^+$, 537.2349; found, 537.234. $[\alpha]_D^{23}$ = + 7.2 ° (*c* = 0.54, DMSO)

* denotes a rotamer
† see "Note on NMR spectra" at the beginning of this section
§ The integral values of these peaks are higher than expected due to overlaps with the water and DMSO solvent residuals. 2D NMR experiments (shown in the following section) show the expected connectivities.

### 7. NMR spectra

Compound **2** $^{1}$H NMR (500 MHz DMSO-$d_6$):

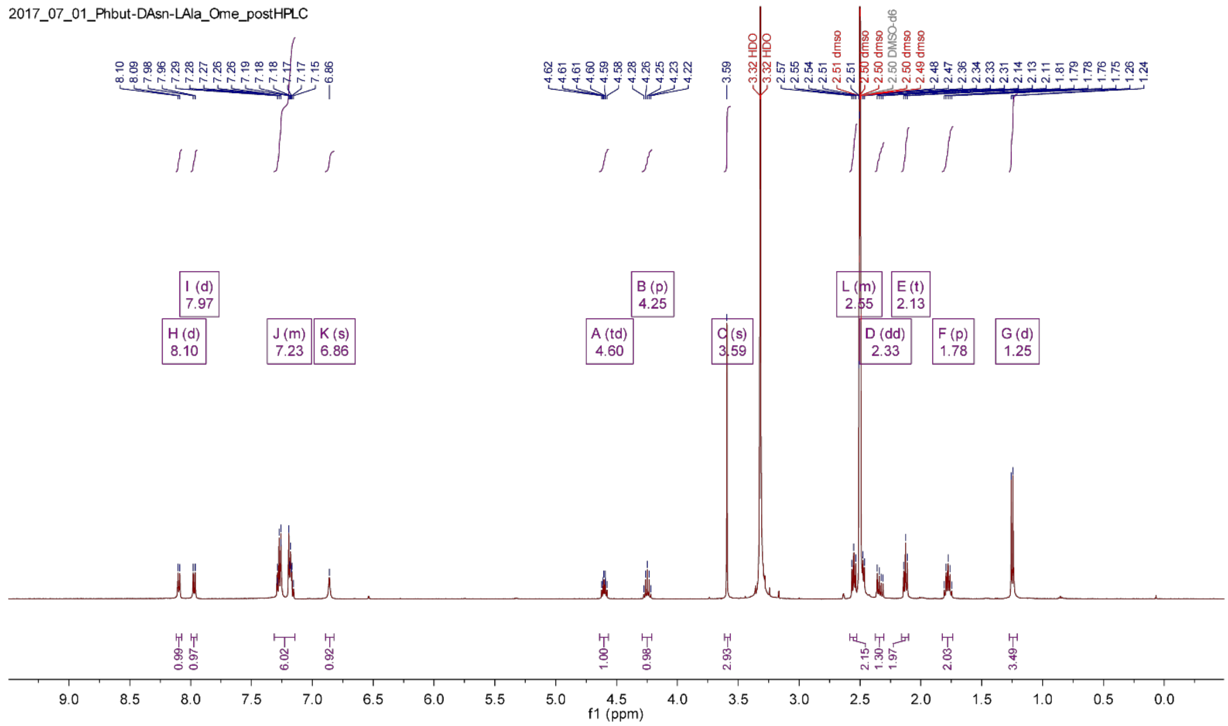

Compound **2** $^{13}$C NMR (500 MHz DMSO-$d_6$):

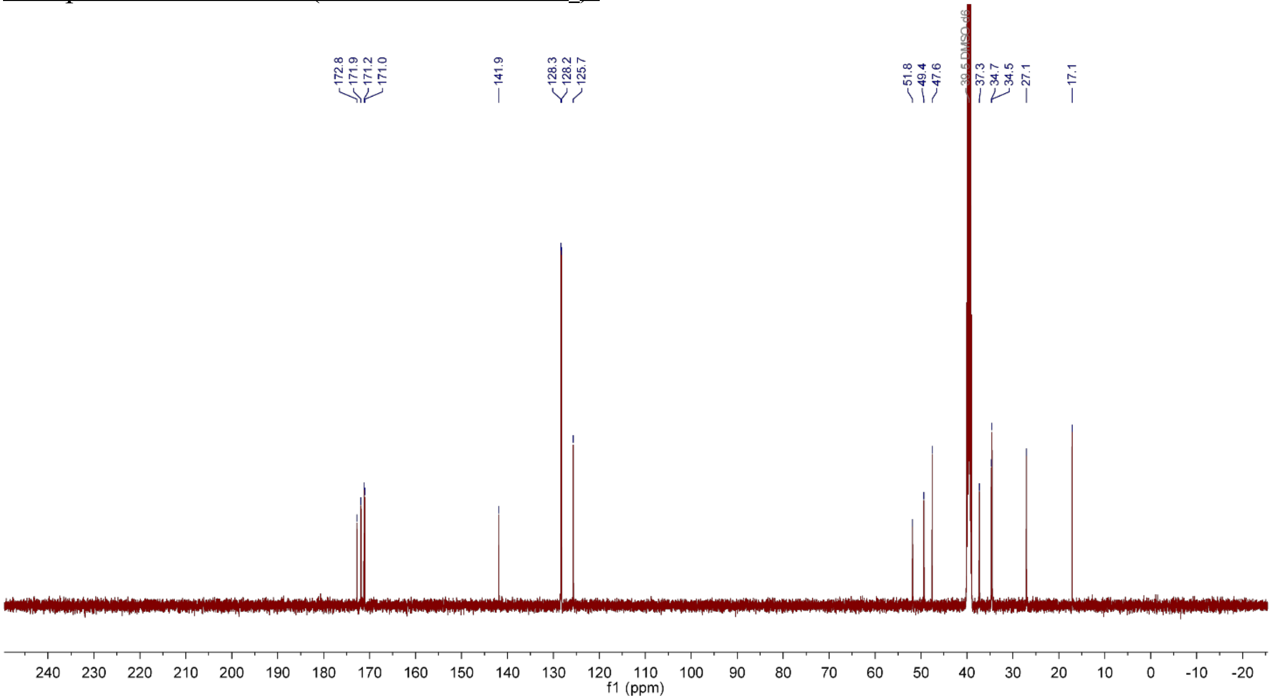

Compound **3** $^{1}$H NMR (500 MHz DMSO-$d_6$):

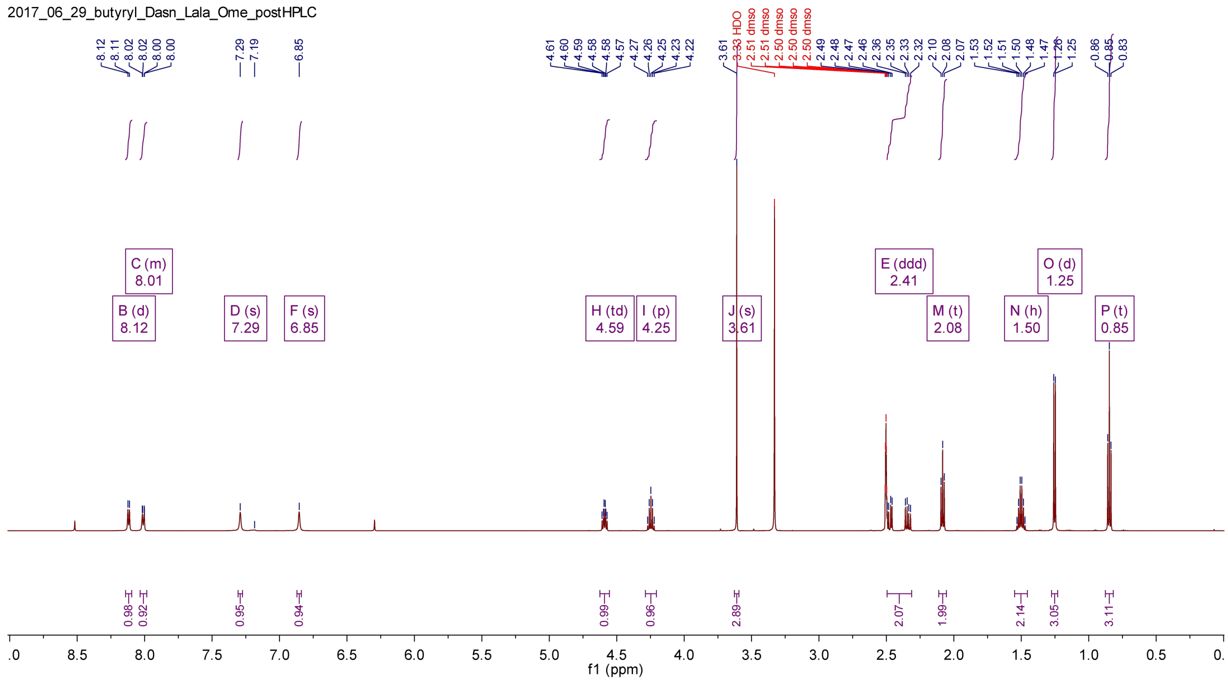

Compound **3** $^{13}$C NMR (500 MHz DMSO-$d_6$):

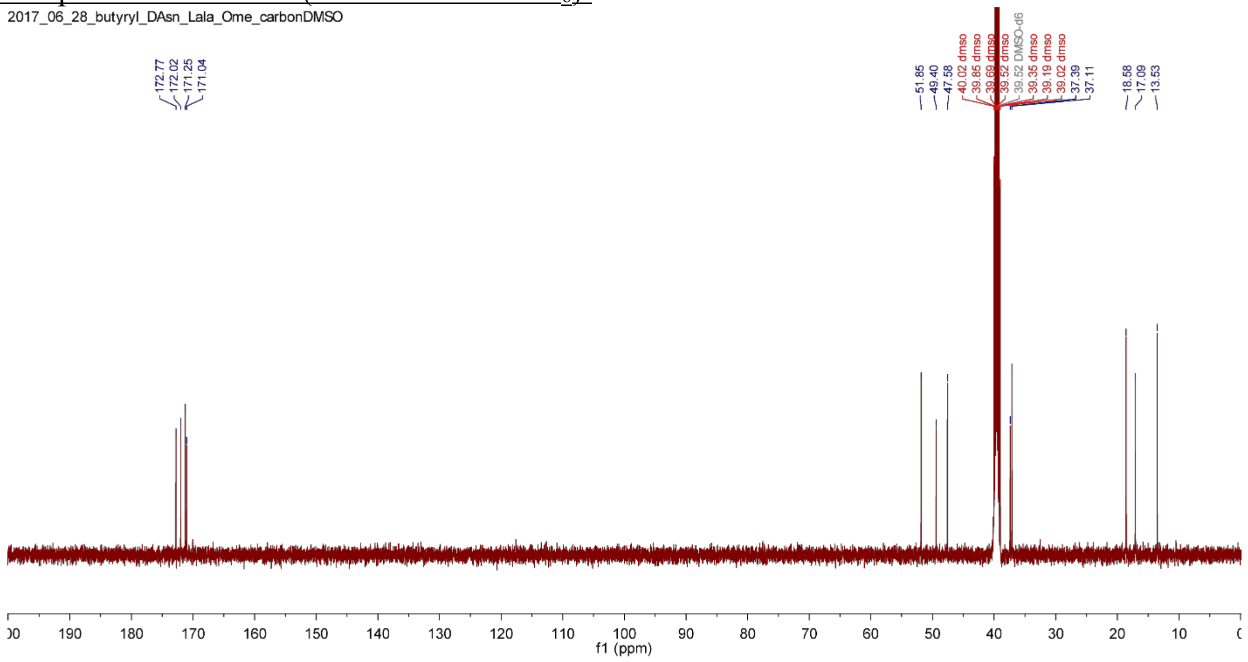

Compound **3** $^1$H-$^1$H COSY (500 MHz DMSO-$d_6$):

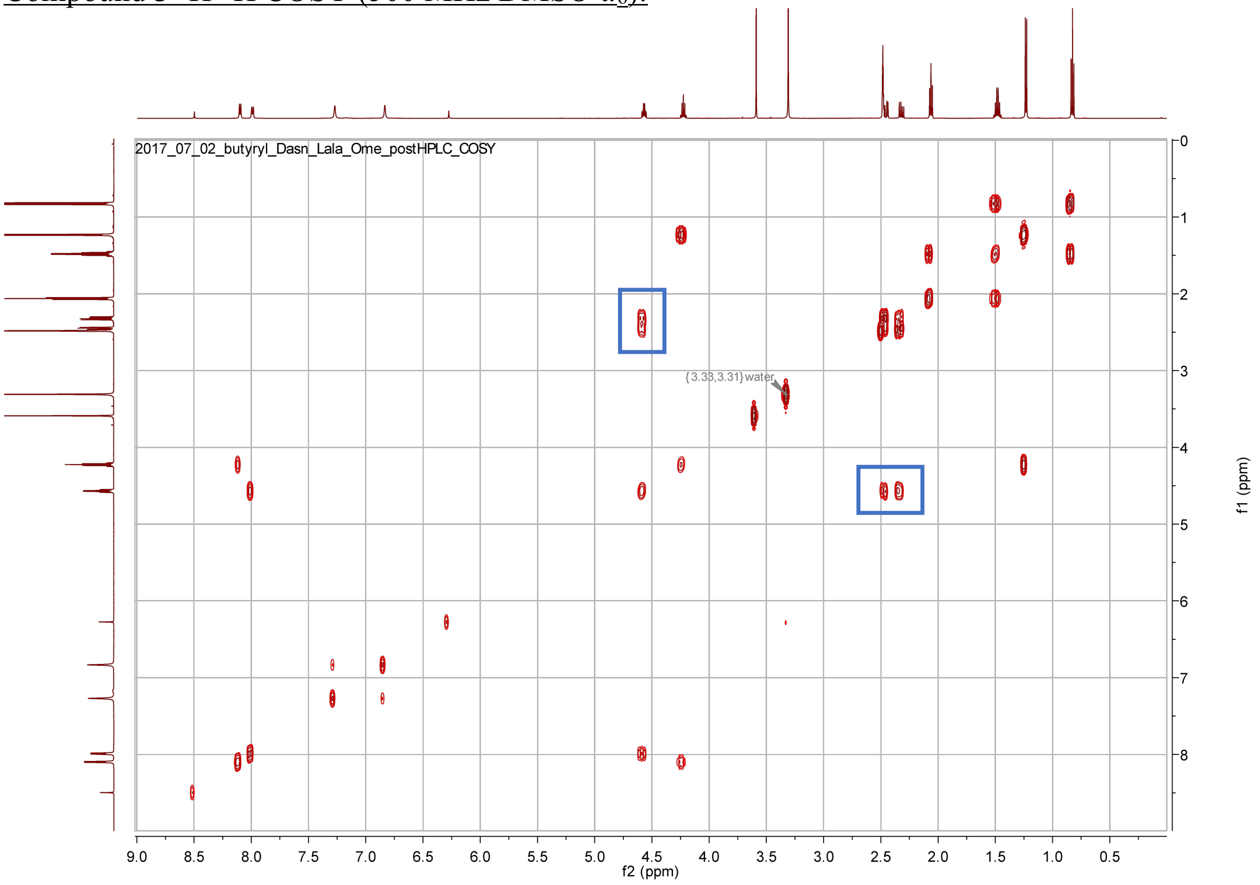

Boxed peaks indicate that there are two distinct signals for the D-Asn β position protons (one of which overlaps partially with the DMSO solvent residual), both of which show a COSY coupling to the D-Asn α-proton

Compound **4** $^1$H NMR (500 MHz DMSO-$d_6$):

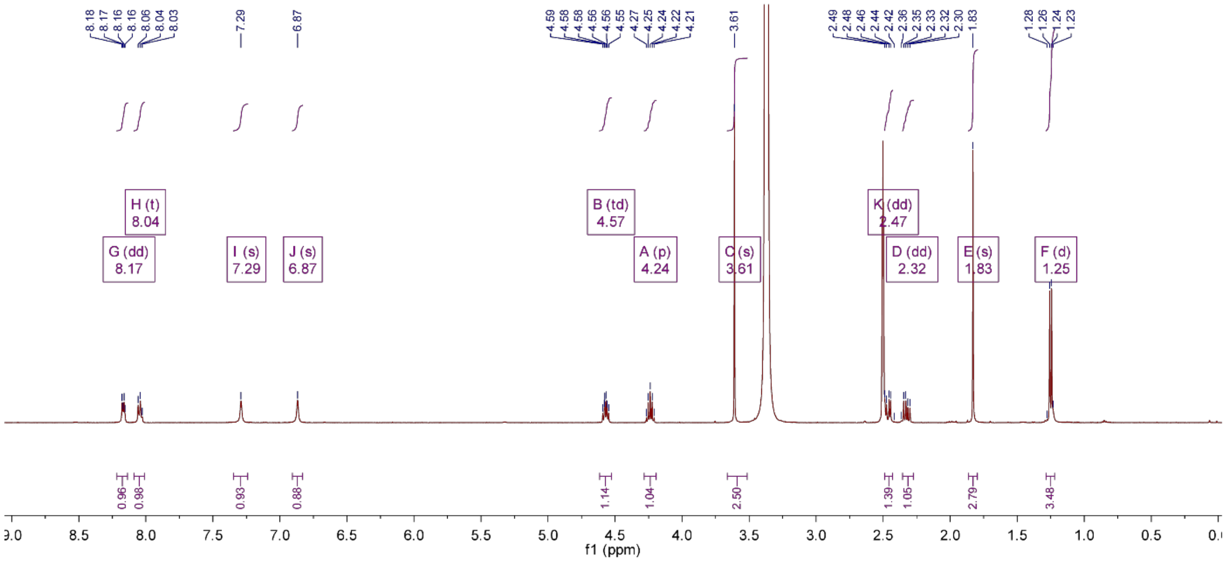

Compound **4** $^{13}C$ NMR (500 MHz DMSO-$d_6$):

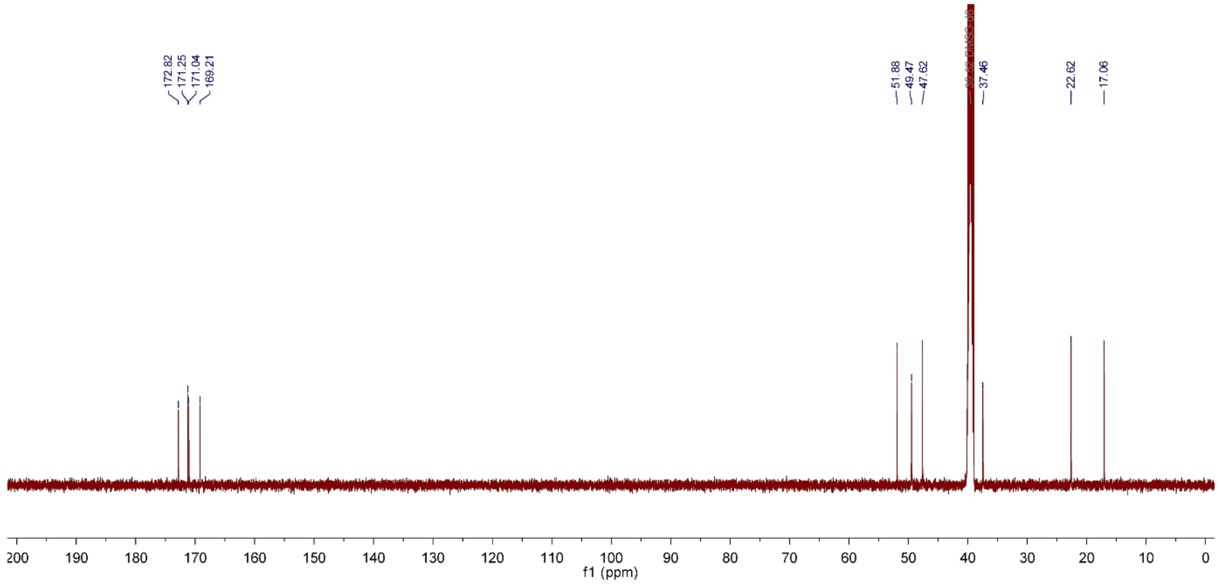

Compound **5** $^{1}H$ NMR (400 MHz DMSO-$d_6$):

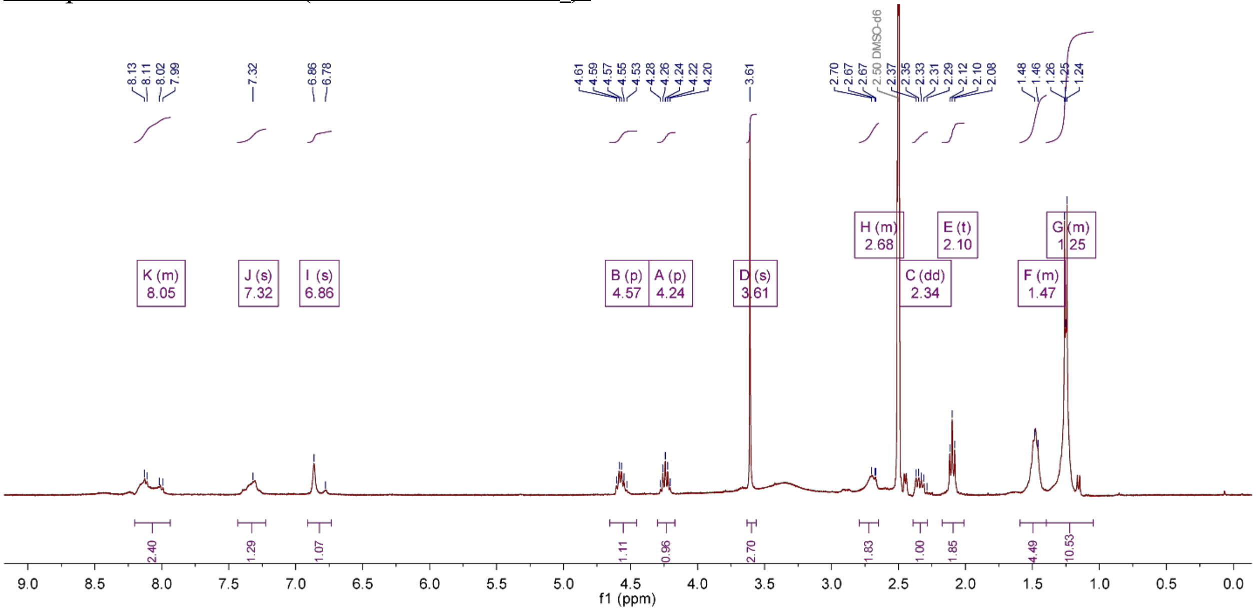

Compound **5** $^{13}$C NMR (500 MHz DMSO-$d_6$):

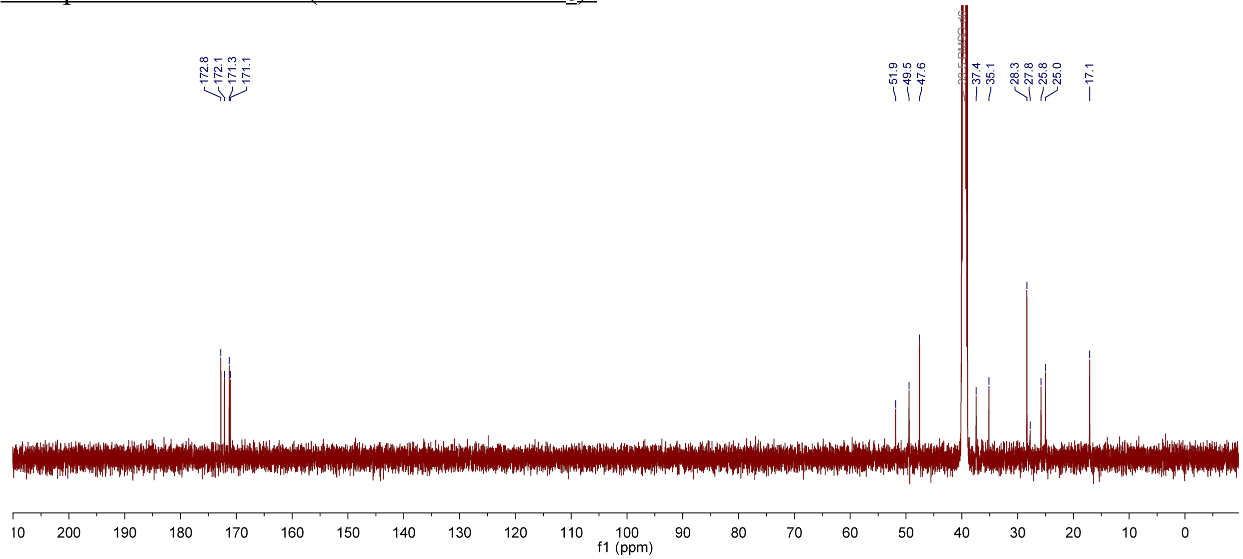

Compound **5** $^{1}$H-$^{13}$C HSQC (400 MHz DMSO-$d_6$):

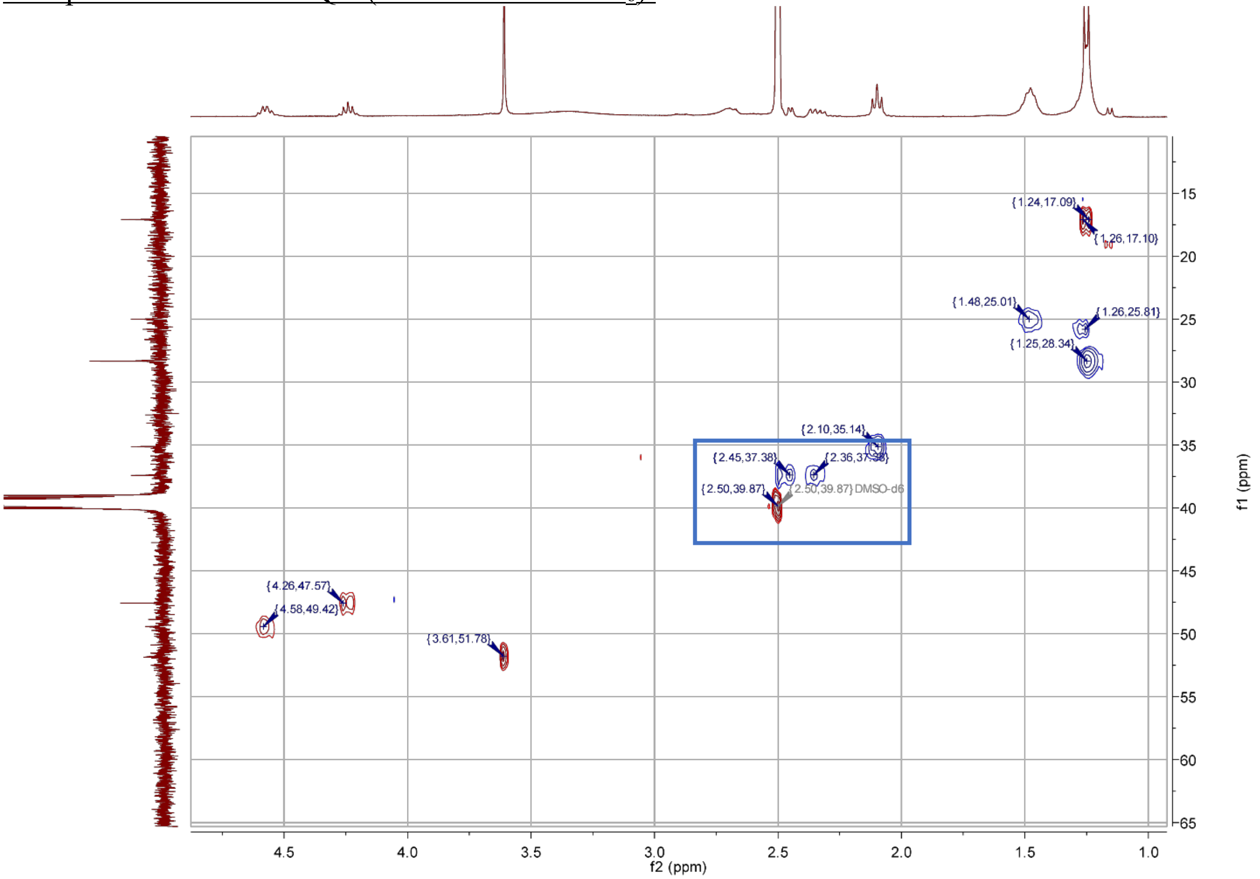

Boxed peaks indicate that there are two distinct signals for the D-Asn β position protons (one of which overlaps partially with the DMSO solvent residual), both of which show an HSQC coupling to the D-Asn β-carbon

Compound **6** $^{1}$H NMR (600 MHz DMSO-$d_6$):

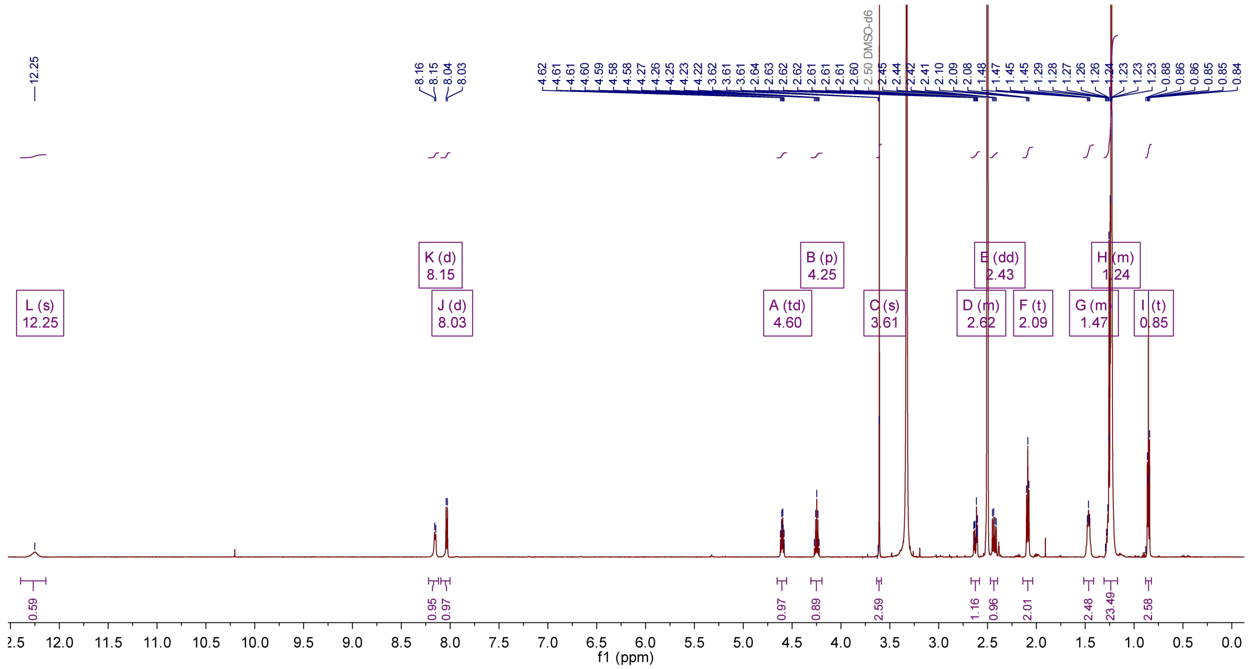

Compound **6** $^{13}$C NMR (400 MHz DMSO-$d_6$):

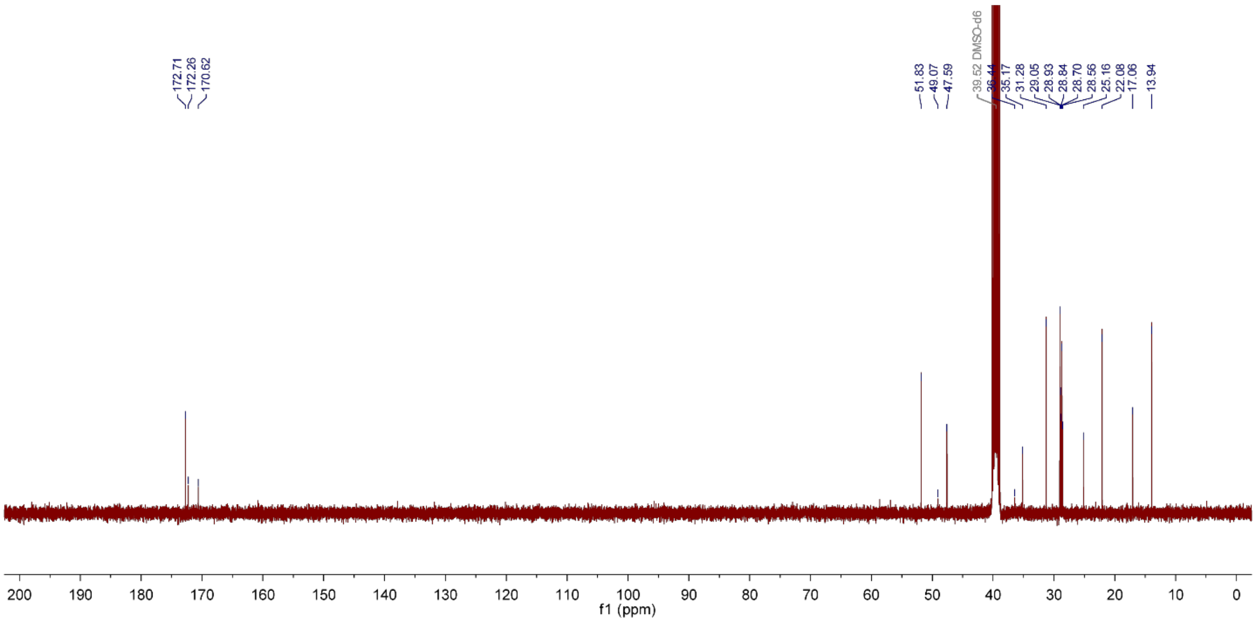

Compound **6** HMBC (400 MHz DMSO-$d_6$):

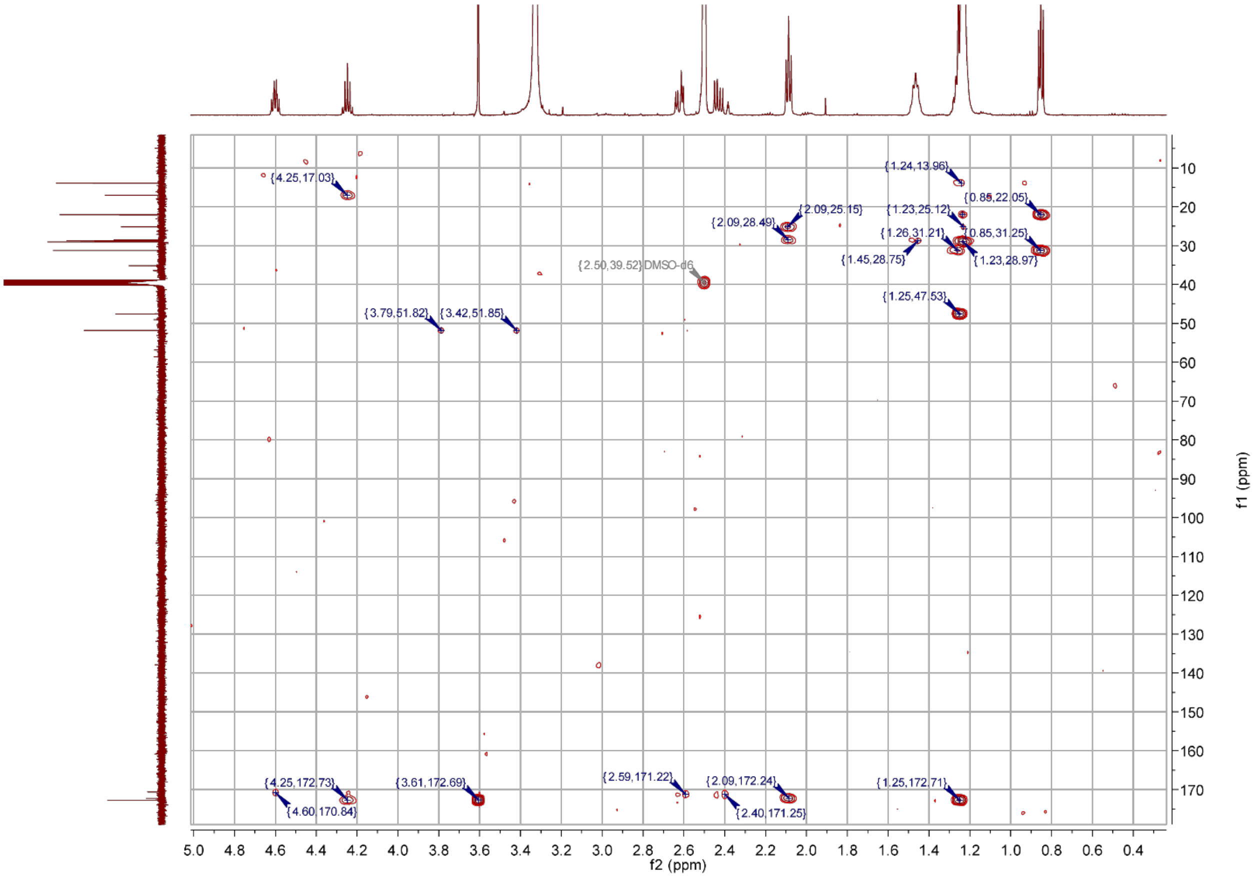

Compound **7** $^{1}$H NMR (600 MHz DMSO-$d_6$):

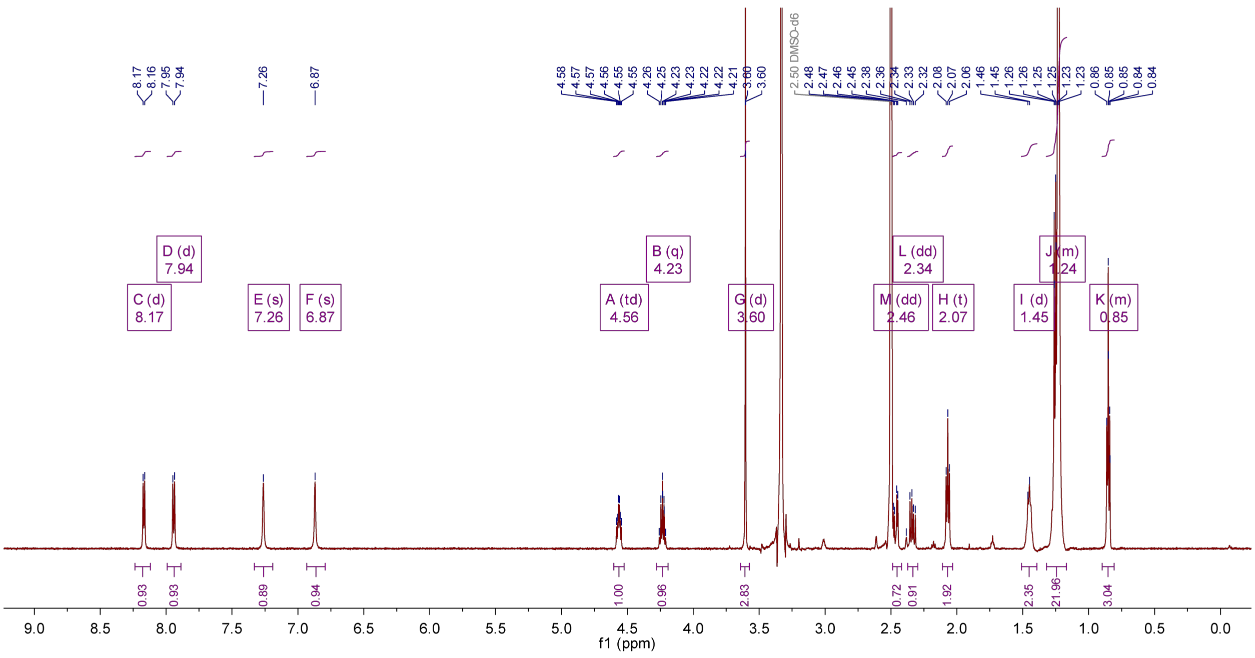

Compound **7** $^{13}$C NMR (400 MHz DMSO-$d_6$):

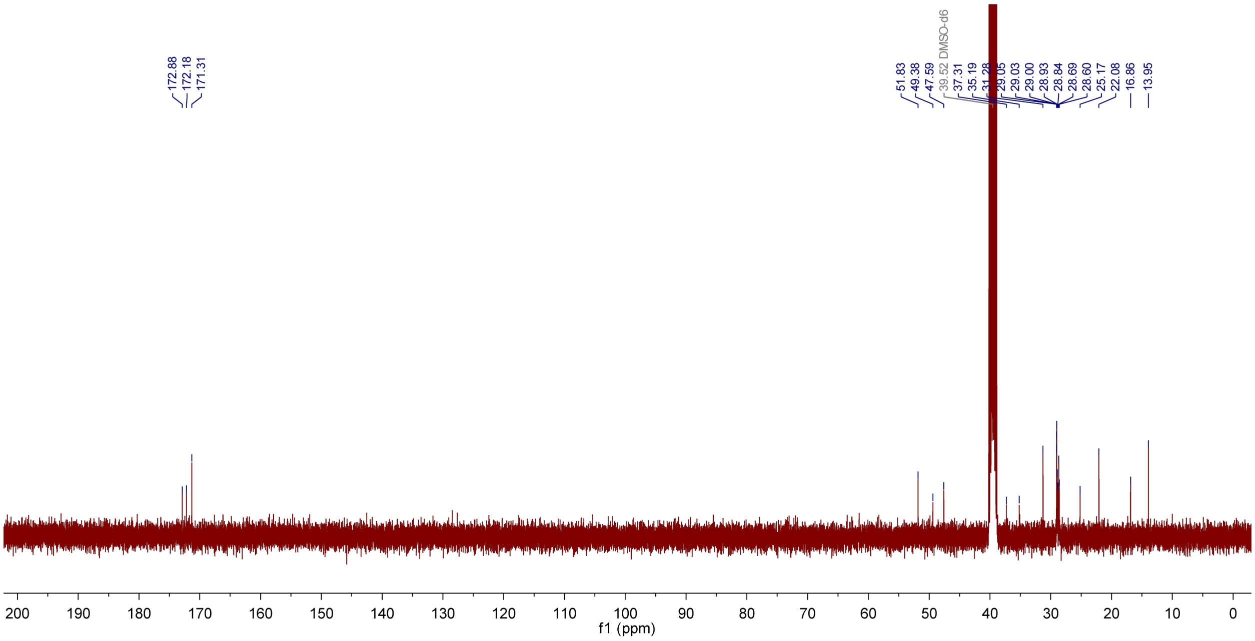

Compound **8** $^{1}$H NMR (500 MHz DMSO-$d_6$):

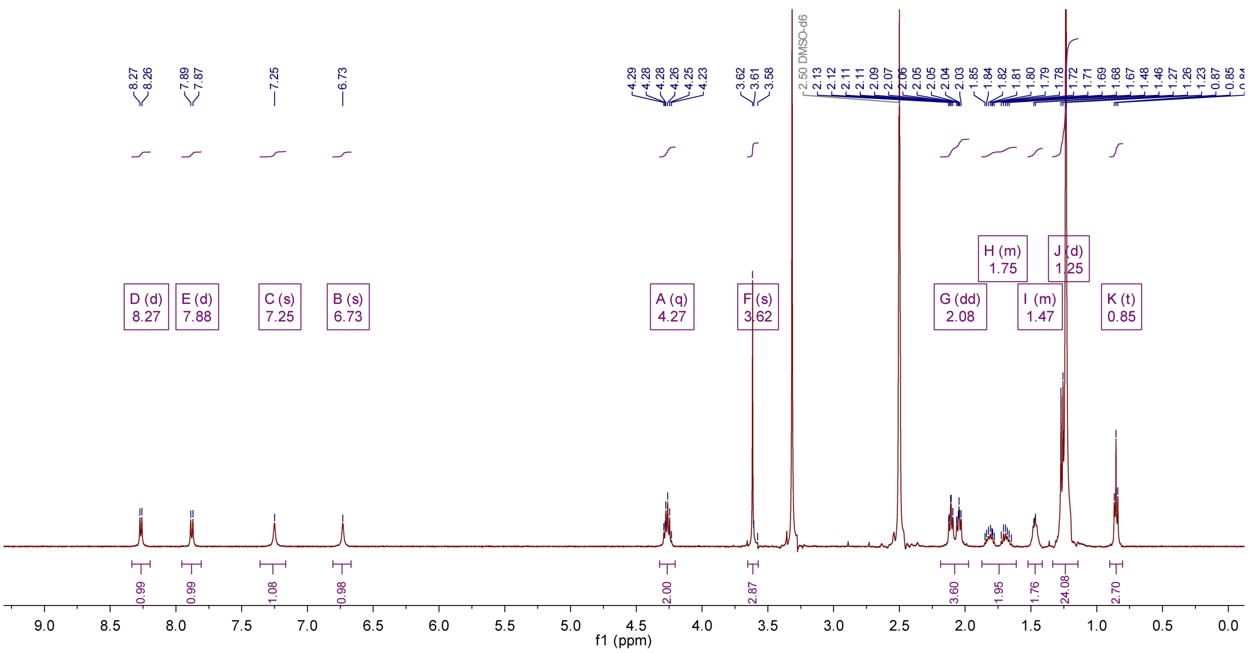

Compound **8** $^{13}$C NMR (500 MHz DMSO-$d_6$):

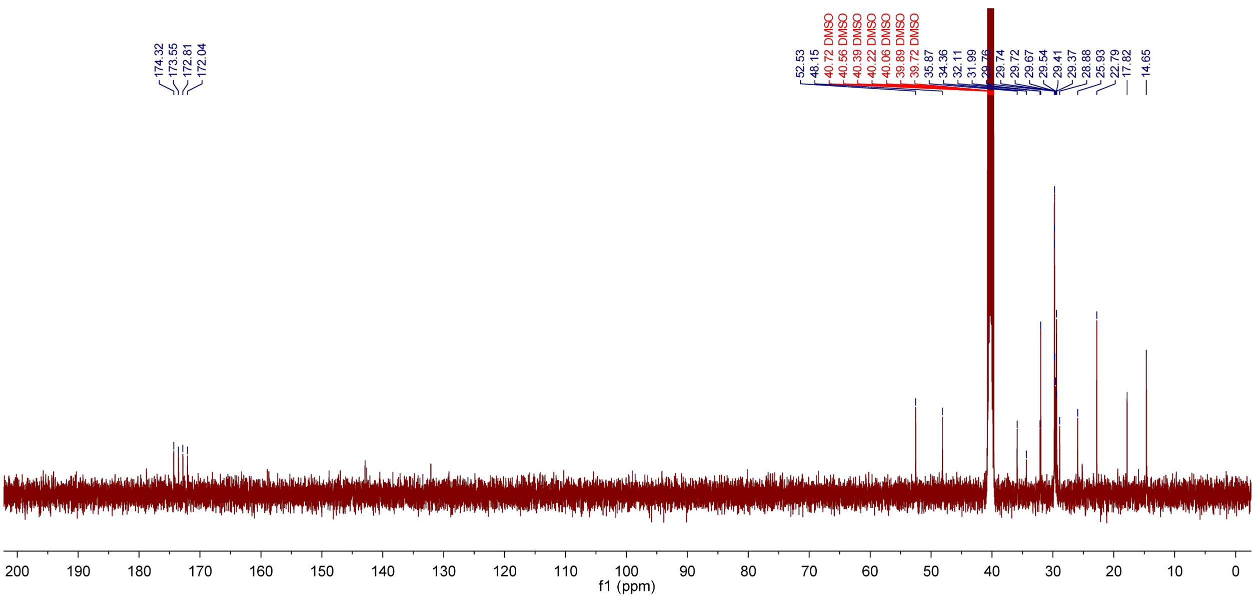

Compound **9** $^{1}$H NMR (400 MHz DMSO-$d_6$):

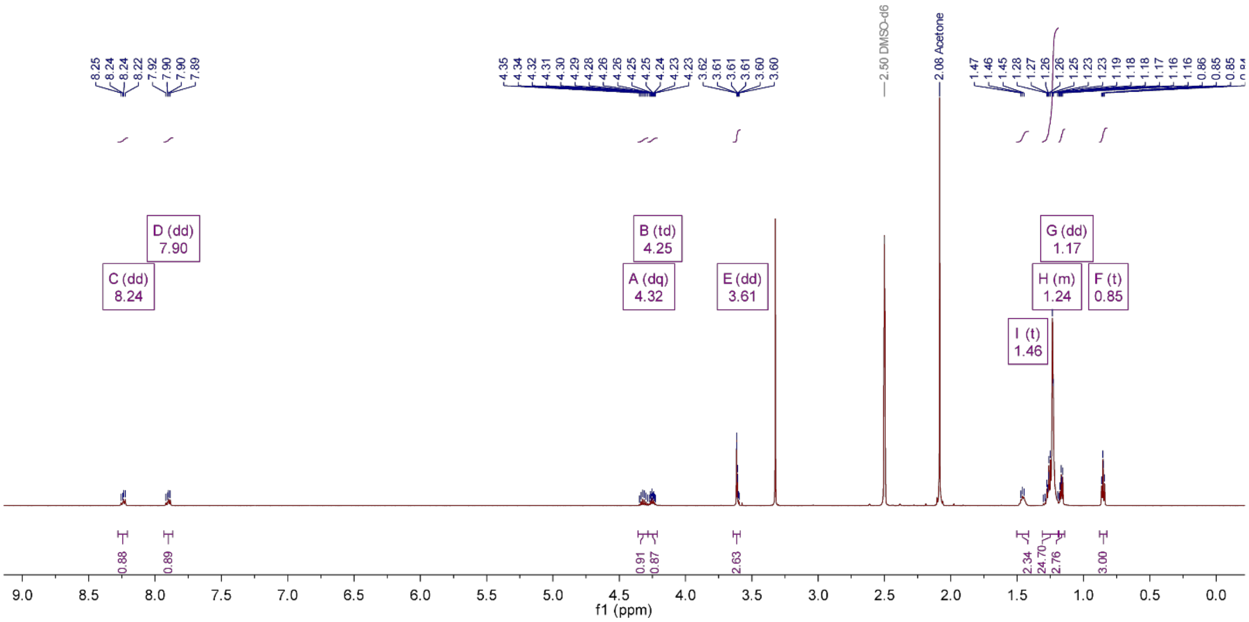

Compound **9** $^{13}$C NMR (400 MHz DMSO-$d_6$):

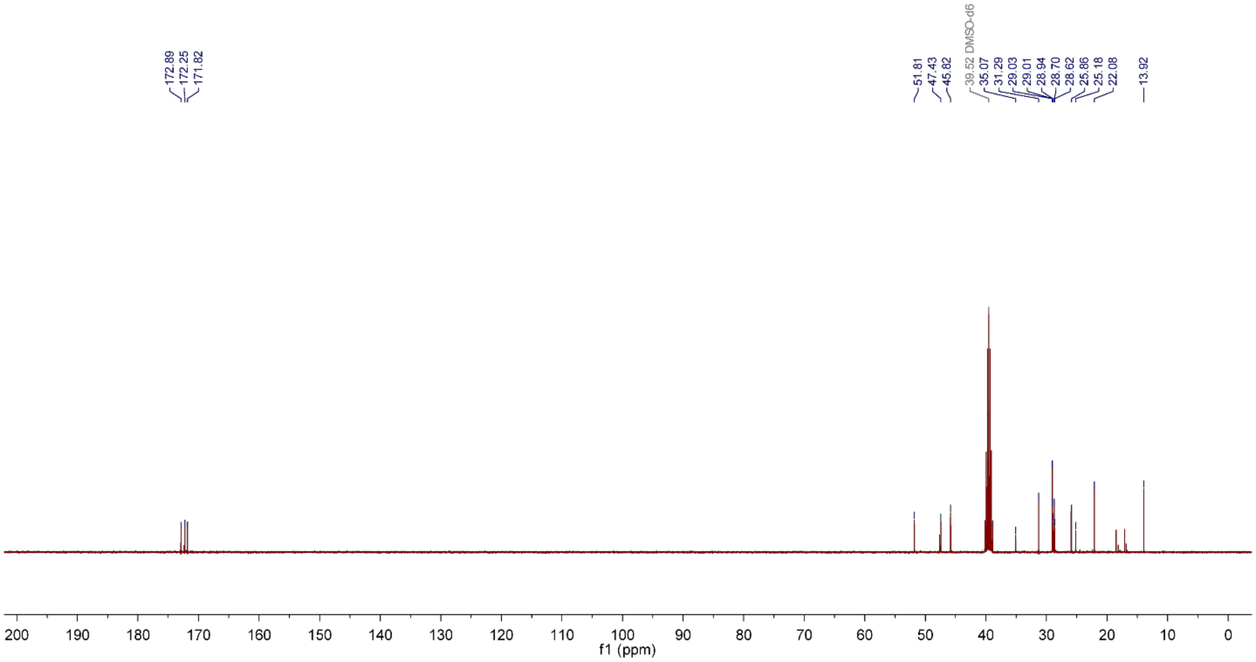

Compound **10** $^{1}$H NMR (600 MHz DMSO-$d_6$):

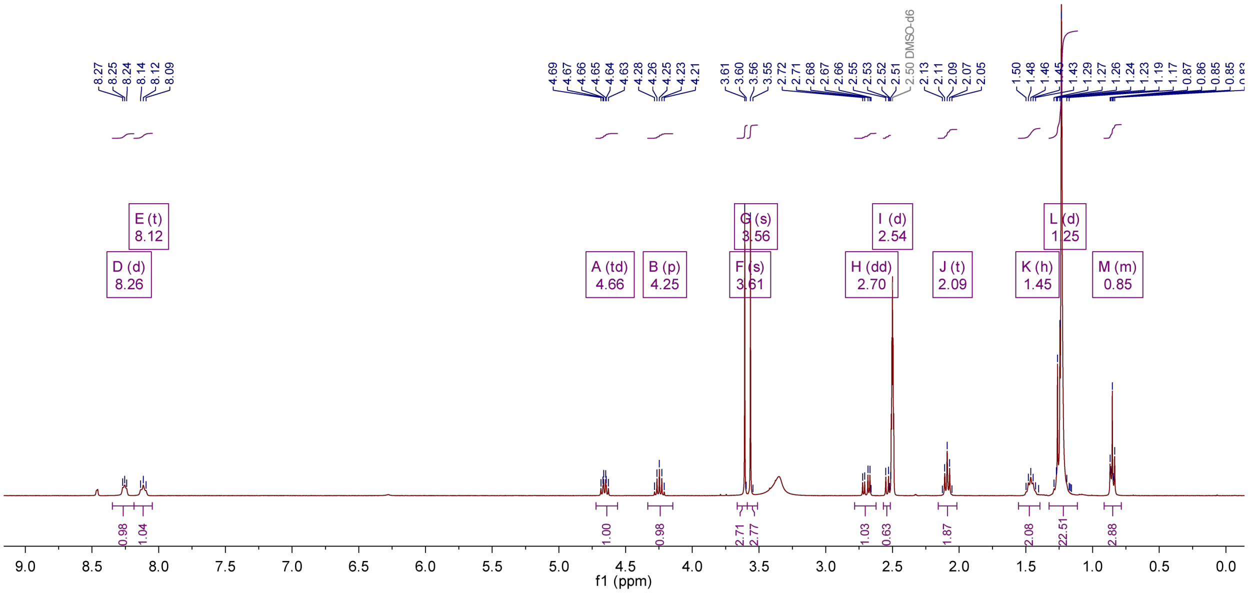

Compound **10** $^{13}$C NMR (500 MHz DMSO-$d_6$):

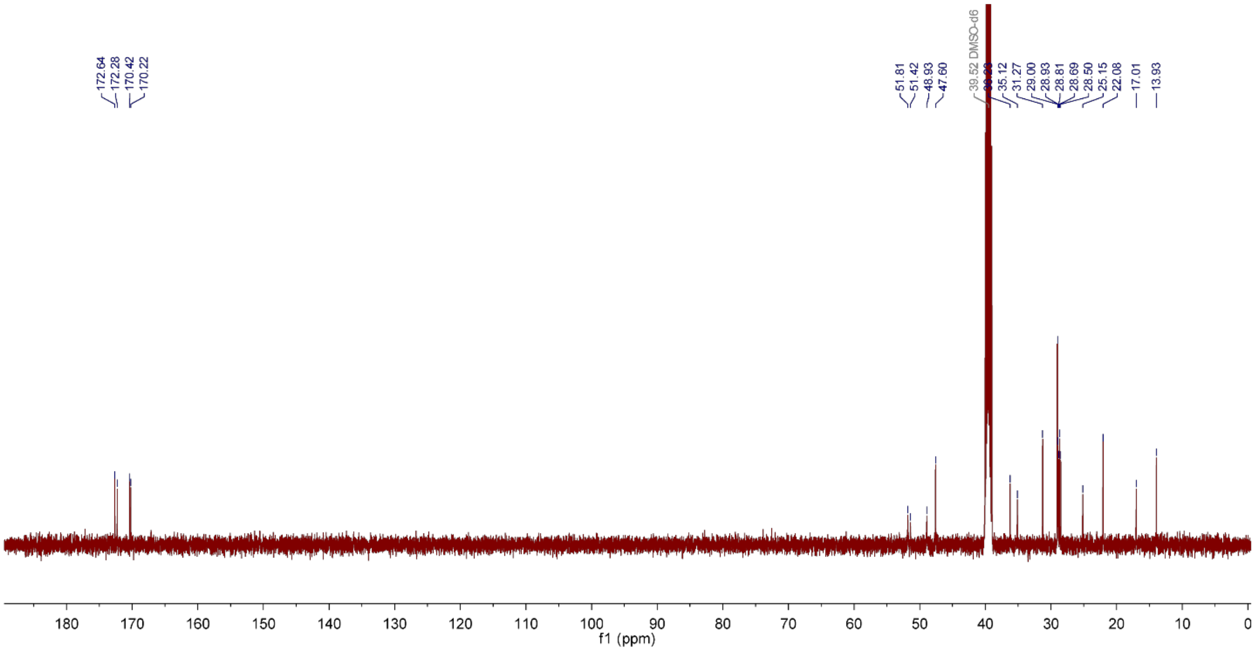

Compound **11** $^{1}$H NMR (500 MHz DMSO-$d_6$):

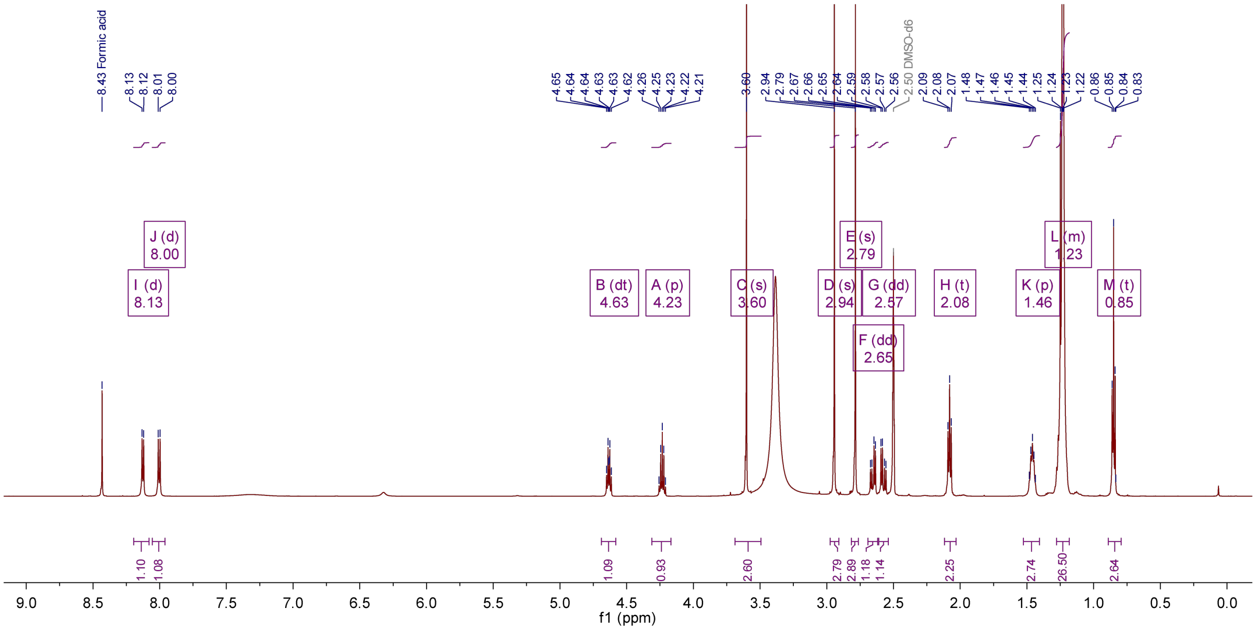

Compound **11** $^{13}$C NMR (500 MHz DMSO-$d_6$):

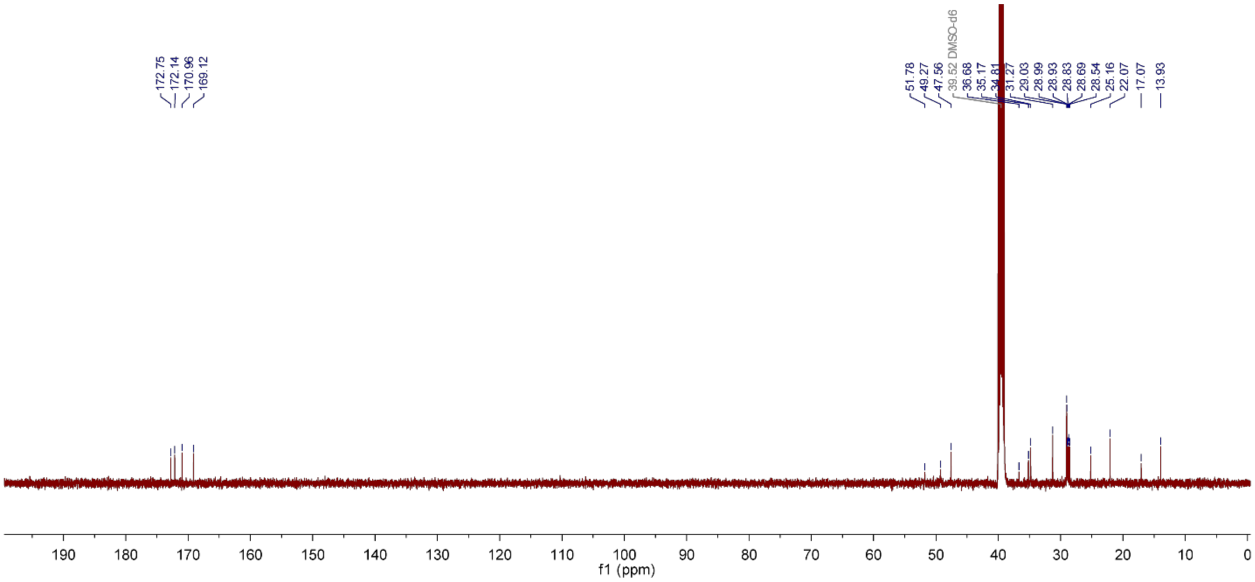

Compound **11** $^{13}C$-$^{1}H$-HSQC (400 MHz DMSO-$d_6$):

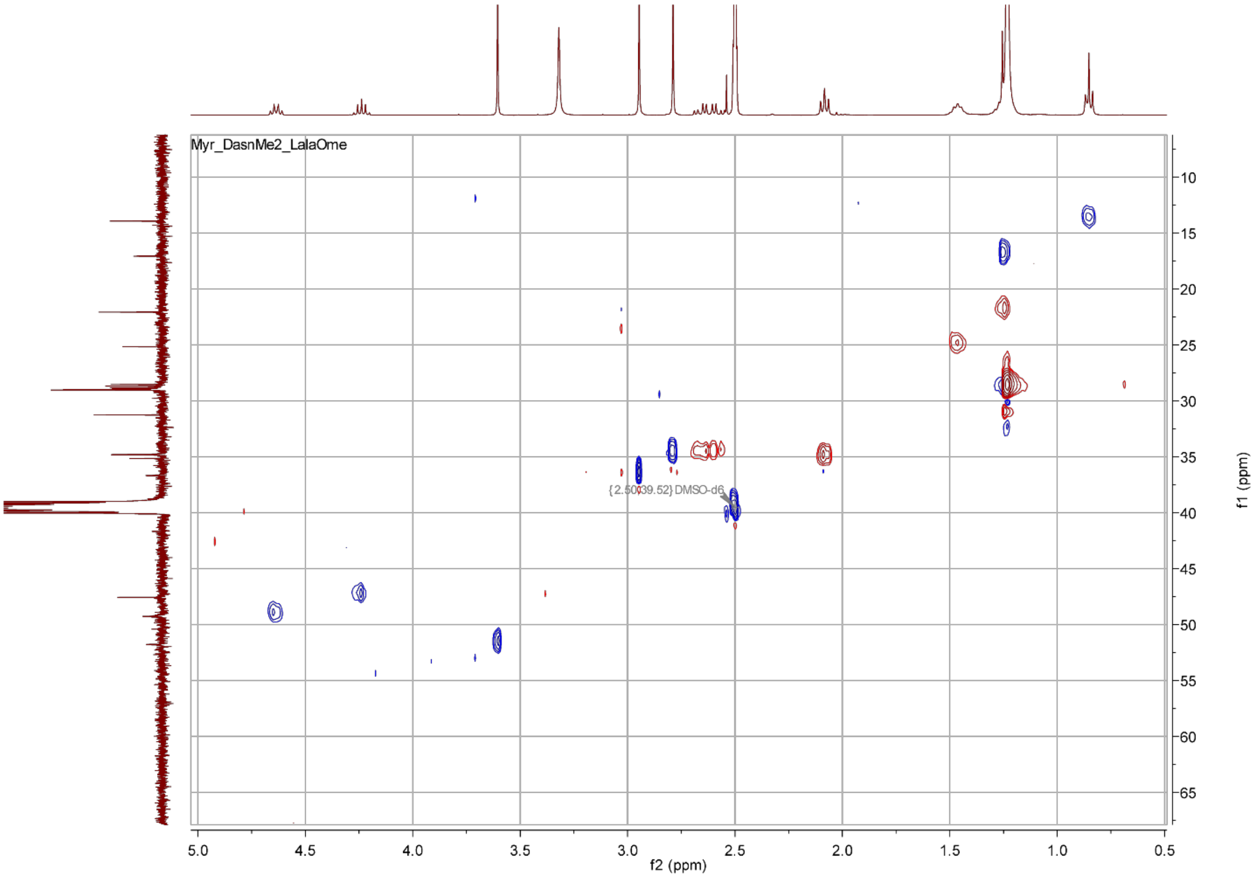

Compound **12** $^{1}$H NMR (400 MHz DMSO-$d_6$):

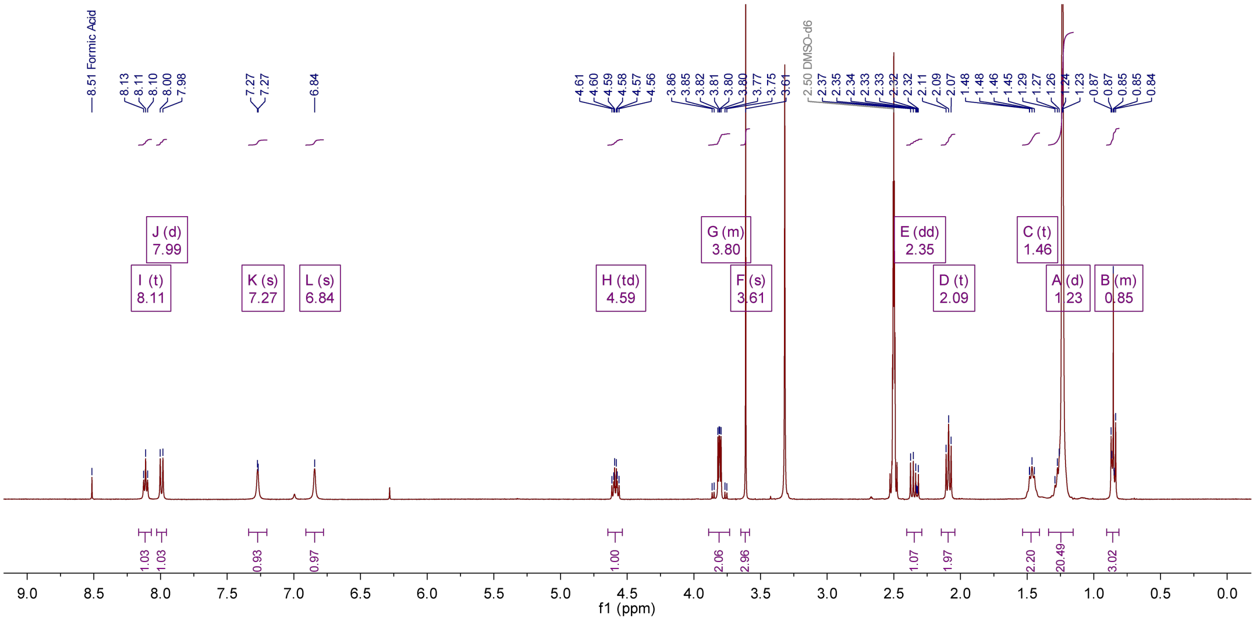

Compound **12** $^{13}$C NMR (400 MHz DMSO-$d_6$):

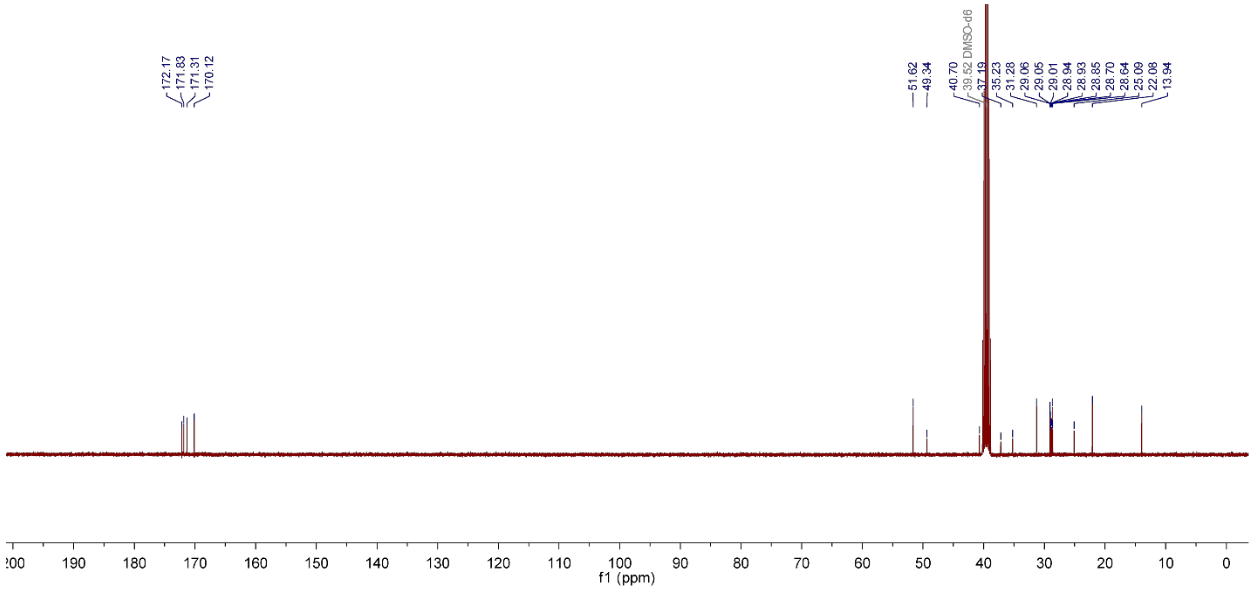

Compound **12** $^{13}C$-$^{1}H$ HSQC (400 MHz DMSO-$d_6$):

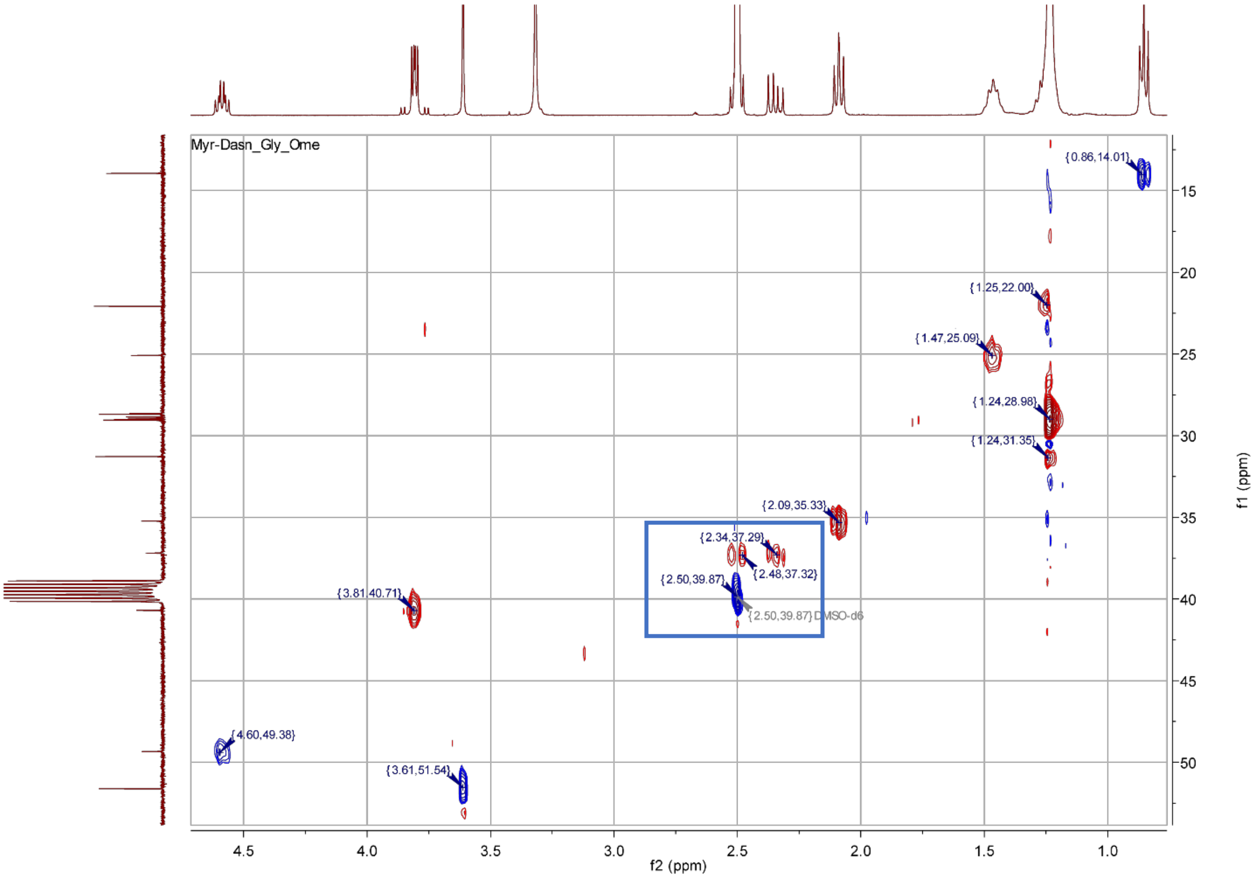

Boxed peaks indicate that there are two distinct signals for the D-Asn β position protons (one of which overlaps partially with the DMSO solvent residual), both of which show an HSQC coupling to the D-Asn β-carbon

Compound **13** $^{1}$H NMR (400 MHz DMSO-$d_6$):

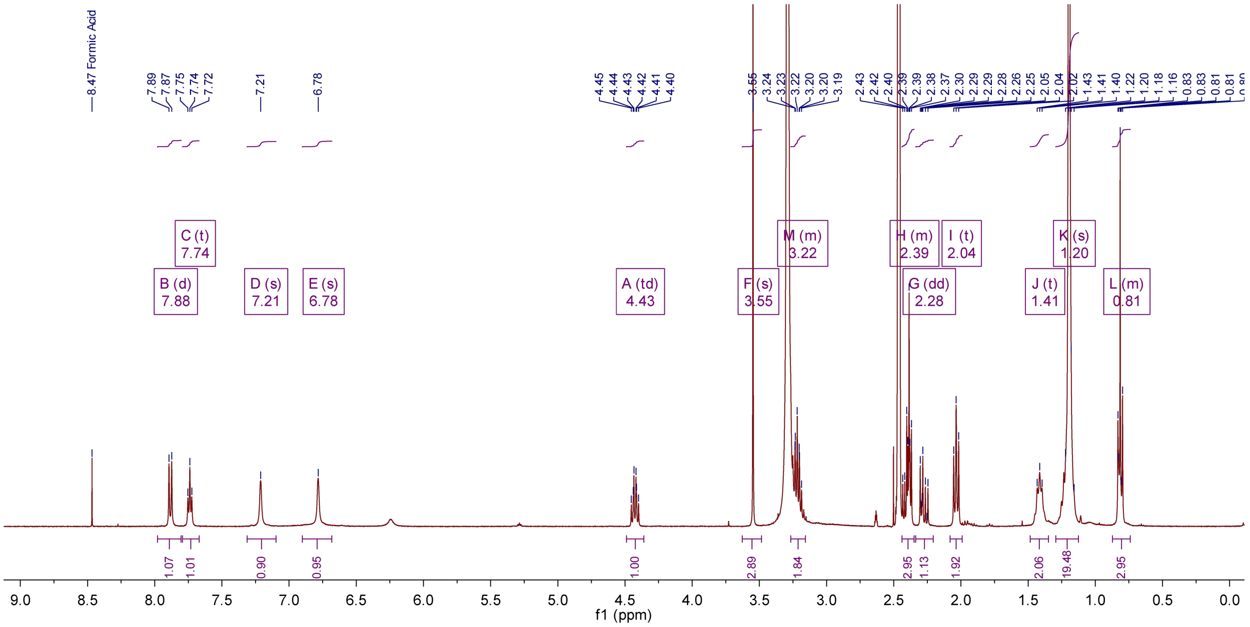

Compound **13** $^{13}$C NMR (400 MHz DMSO-$d_6$):

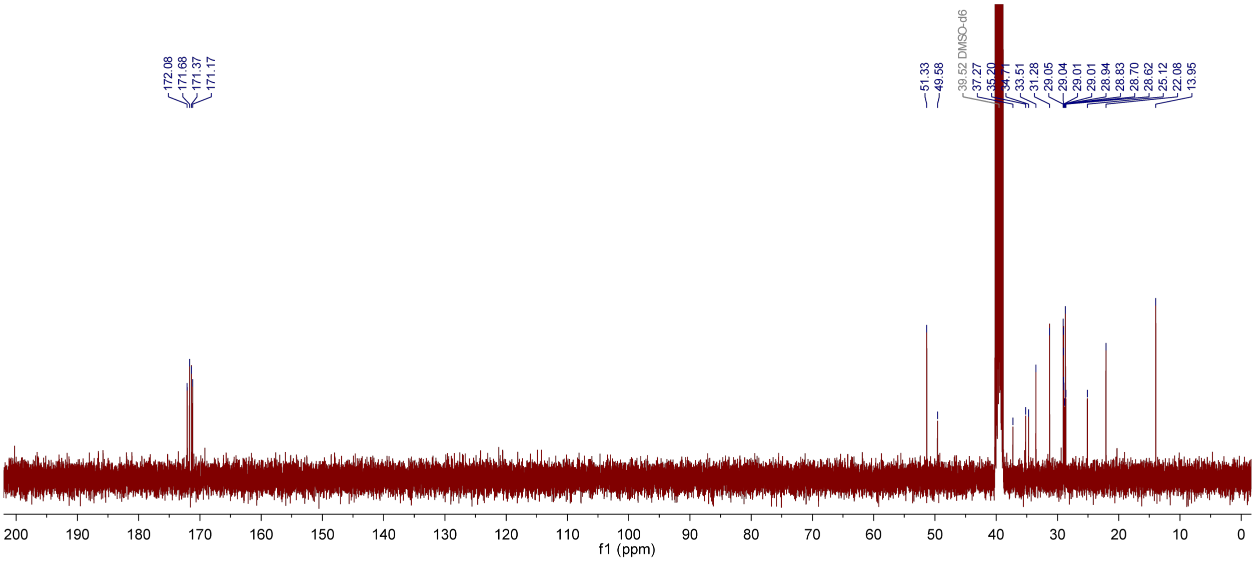

Compound **15** $^{1}$H NMR (400 MHz DMSO-$d_6$):

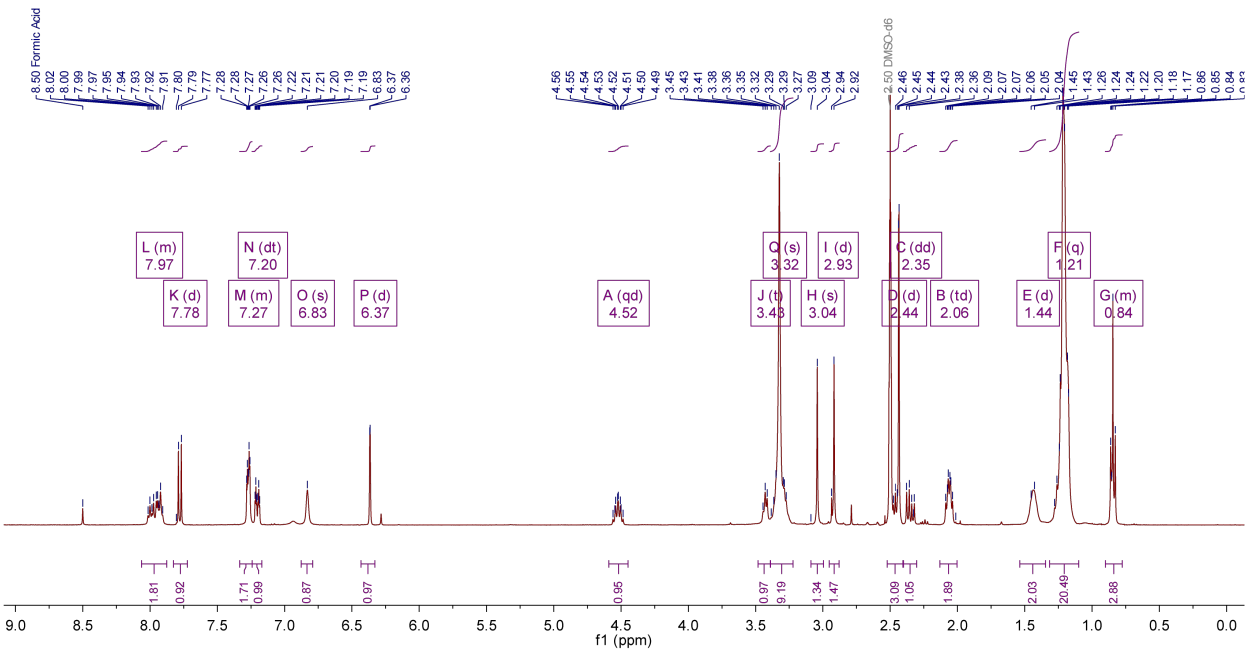

Compound **15** $^{13}$C NMR (400 MHz DMSO-$d_6$):

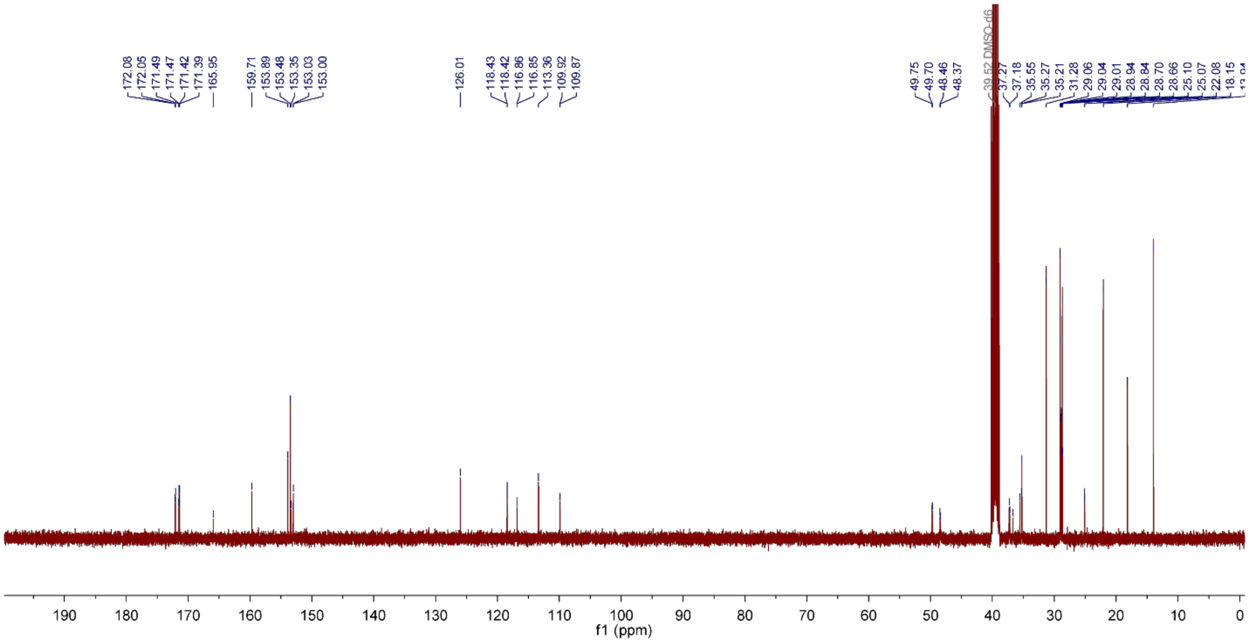

Compound **15** $^{13}C$-$^{1}H$ HSQC (400 MHz DMSO-$d_6$):

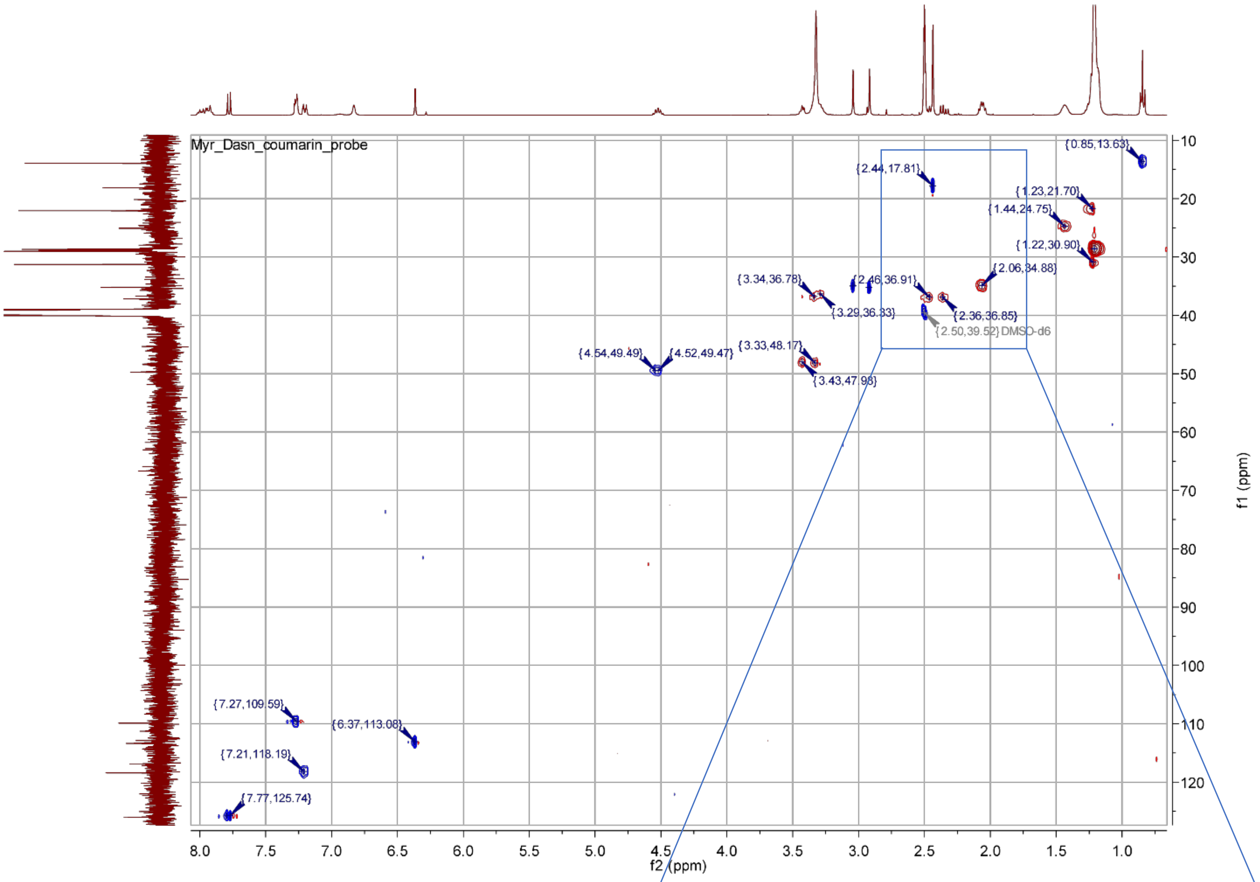

Probe **15** exhibits the same splitting pattern for the proton signals at the D-Asn β position as exhibited by many of the other substrates shown above (one peak partially overlaps with the DMSO signal). There is an additional signal in this group originating from the methyl group attached to the coumarin ring system (see inset at right)

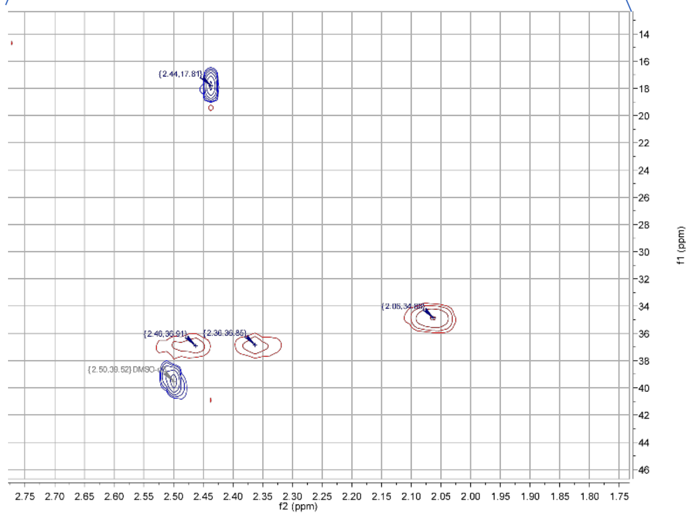

Compound **16** $^{1}$H NMR (600 MHz, DMSO-$d_6$):

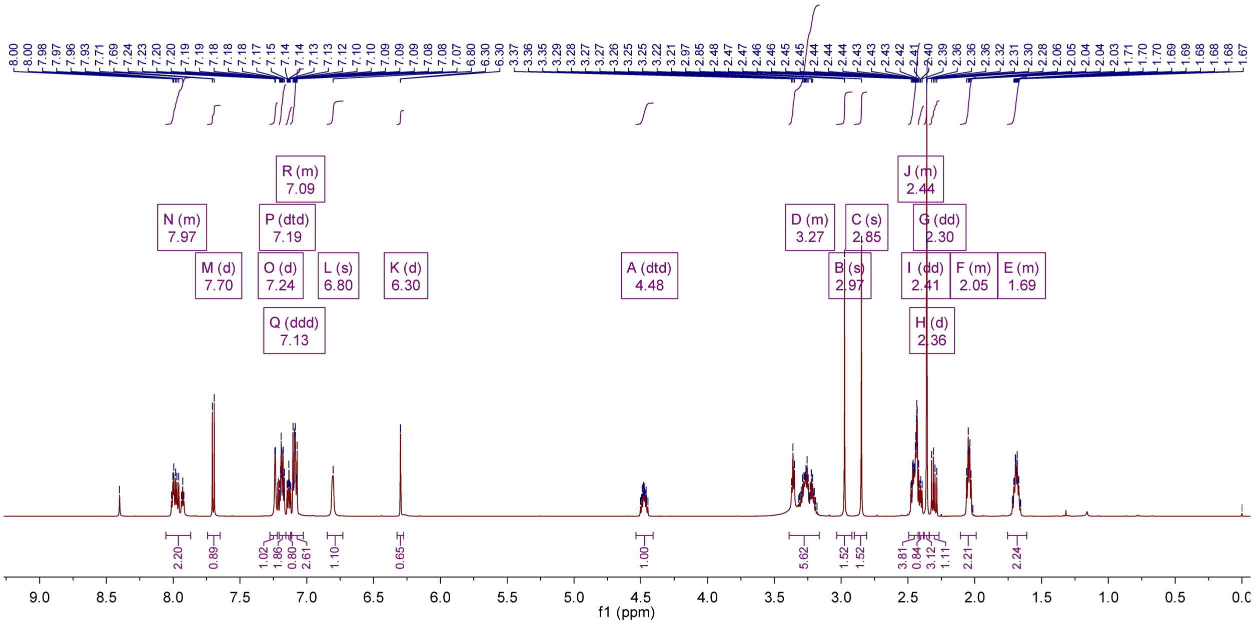

Compound **16** $^{13}$C NMR (500 MHz, DMSO-$d_6$):

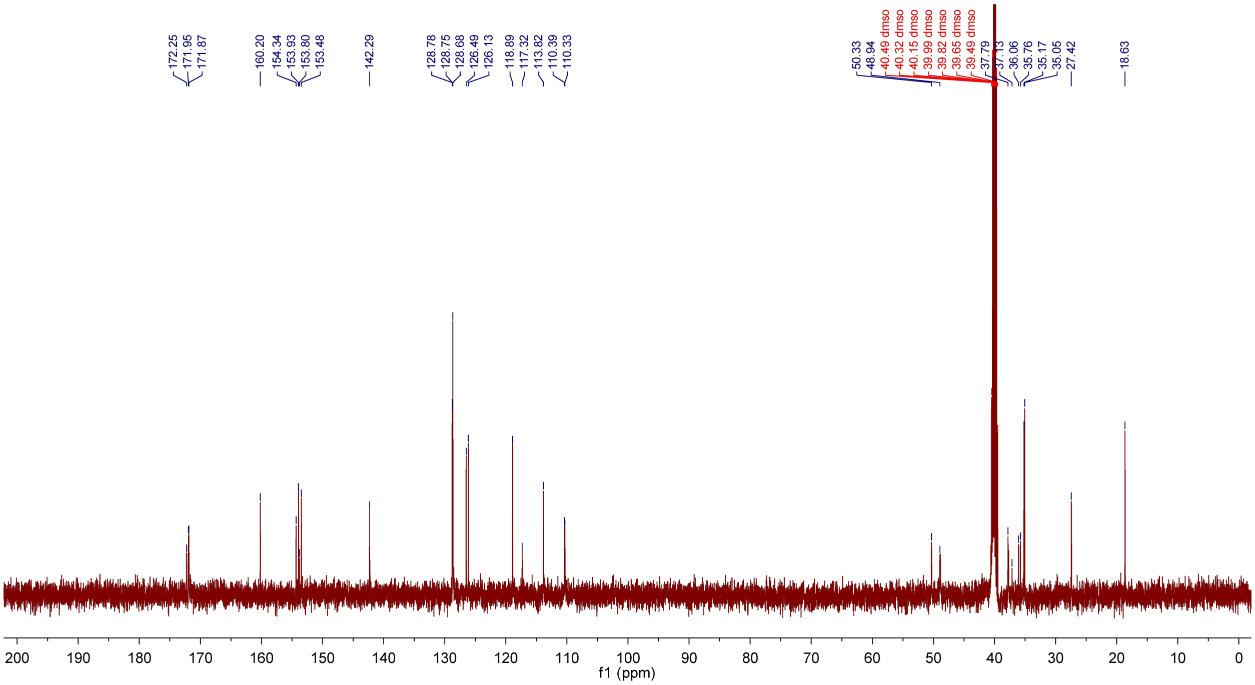

Compound **16** COSY (600 MHz, DMSO-$d_6$):

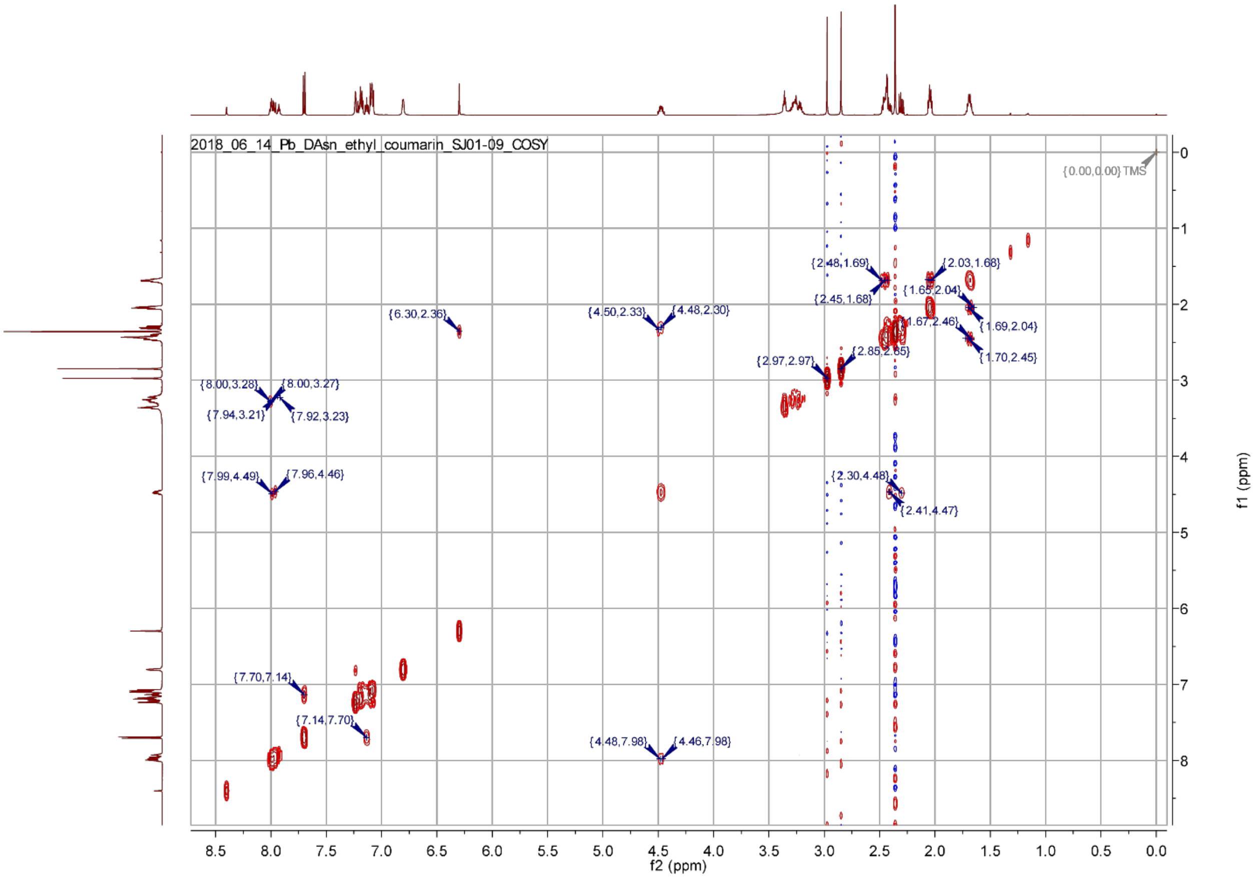
